# Determinants of QTL mapping power in the realized Collaborative Cross

**DOI:** 10.1101/459966

**Authors:** Gregory R. Keele, Wesley L. Crouse, Samir N. P. Kelada, William Valdar

## Abstract

The Collaborative Cross (CC) is a mouse genetic reference population whose range of applications includes quantitative trait loci (QTL) mapping. The design of a CC QTL mapping study involves multiple decisions, including which and how many strains to use, and how many replicates per strain to phenotype, all viewed within the context of hypothesized QTL architecture. Until now, these decisions have been informed largely by early power analyses that were based on simulated, hypothetical CC genomes. Now that more than 50 CC strains are available and more than 70 CC genomes have been observed, it is possible to characterize power based on realized CC genomes. We report power analyses based on extensive simulations and examine several key considerations: 1) the number of strains and biological replicates, 2) the QTL effect size, 3) the presence of population structure, and 4) the distribution of functionally distinct alleles among the founder strains at the QTL. We also provide general power estimates to aide in the design of future experiments. All analyses were conducted with our R package, SPARCC (Simulated Power Analysis in the Realized Collaborative Cross), developed for performing either large scale power analyses or those tailored to particular CC experiments.

## Introduction

The Collaborative Cross (CC) is a multiparental population (MPP) recombinant inbred (RI) strain panel of laboratory mice derived from eight inbred founder strains (letter abbreviation in parentheses): A/J (A), C57BL/6J (B), 129S1/SvImJ (C), NOD/ShiLtJ (D), NZO/H1LtJ (E), CAST/EiJ (F), PWK/PhJ (G), and WSB/EiJ (H) (Threadgill *et al.* 2002; Churchill *et al.* 2004; Chesler *et al.* 2008; Threadgill and Churchill 2012). This set of founder strains represents three subspecies of the house mouse *Mus musculus* (Yang *et al.* 2011) and, in large part due to the inclusion of three wild-derived founders (F-H), imbues the CC panel with far greater genetic variation than previous RI panels derived solely from pairs of classical inbred strains. As an RI panel, the CC thus provides a diverse set of reproducible genomes and represents a powerful tool for genetic analysis (Collaborative Cross Consortium 2012; Srivastava *et al.* 2017). Indeed, although the CC RI panel has only become available in the last six years (Welsh *et al.* 2012), it has already yielded new insights into human disease and basic mouse biology (Shusterman *et al.* 2013; Rogala *et al.* 2014; Rasmussen *et al.* 2014; Lorè *et al.* 2015; Levy *et al.* 2015; Gralinski *et al.* 2015; Venkatratnam *et al.* 2017; Orgel *et al.* 2019; Molenhuis *et al.* 2018).

As originally envisaged, a key use of the CC was as a resource for QTL mapping (Threadgill *et al.* 2002; Churchill *et al.* 2004). In theory, its broad genetic diversity makes it ideal for this purpose, and its replicability permits the mapping of phenotypes such as drug-response that are otherwise hard to measure in outbreds (Mosedale *et al.* 2017). Its utility for QTL mapping in practice was also predicted by studies in the incipient CC lines (pre-CC) (Aylor *et al.* 2011; Durrant *et al.* 2011; Philip *et al.* 2011; Mathes *et al.* 2011; Kelada *et al.* 2012; Ferris *et al.* 2013; Ram *et al.* 2014; Rutledge *et al.* 2014; Kelada 2016; Donoghue *et al.* 2017; Phillippi *et al.* 2014)

Nonetheless, QTL mapping power depends in part on the number of strains available, and the number strains available in the CC is, and will remain, far less than the 1,000 proposed in Churchill *et al.* (2004): At the time of this work, mice were available for 59 CC strains from the UNC Systems Genetics Core, with a subset from these 59 and an additional 11 expected to be offered through the Jackson Laboratory (JAX), a total of 70 CC strains potentially.

A reduction in strain numbers as a function of allelic incompatibilities between subspecies (Shorter *et al.* 2017) was expected, and winnowed the number of resulting CC strains down to 50-70. Although smaller than originally intended, this population size reflects the biological and financial realities of maintaining a sustainable mammalian genome reference population. [Whereas cost grows proportional to the the number of strains, demand does not, and a much larger number of strains would threaten the economic viability of the operation (F. Pardo-Manuel de Villena, *pers. comm*.).] Nonetheless, subsets of the available CC strains have already been used to map QTL, as evidenced by a growing list of studies (Vered *et al.* 2014; Mosedale *et al.* 2017; Graham *et al.* 2017). Beyond these successes, however, it is unclear how much the reduction has affected the ability to map QTL in the CC in general.

The initially proposed figure of 1,000 CC strains in Churchill *et al.* (2004) was more formally justified in Valdar *et al.* (2006a) as being necessary to provide enough power both to map single QTL and for robust, genome-wide detection of epistasis. That estimate was based on simulations involving larger numbers (500-1,000) of hypothetical CC genomes. Those simulations, performed before any CC strains existed and with the goal of guiding the CC’s design, had a broad scope, exploring the effect of varying strain numbers, alternative mapping approaches [association of single nucleotide polymorphisms (SNPs) vs association of inferred haplotypes], and alternative breeding strategies. As such, the power estimates that were reported do not reflect the number of CC strains now available, nor their actual, realized founder mosaic genomes. An updated, more focused power analysis that both exploits and works within the constraints of the realized genomes is therefore timely.

Power analyses have been performed previously for a number of RI panels. For biparental RIs, they have been performed analytically in plants (*e.g*., Kaeppler 1997), animals [*e.g*., the BXD lines in mice (Belknap *et al.* 1996; Peirce *et al.* 2004)], and in general (Cowen 1988; Soller and Beckmann 1990; Knapp and Bridges 1990), as well as through simulation (Falke and Frisch 2011; Takuno *et al.* 2012). For MPP RIs, they have most often been reported as those resources are introduced to the community. This includes, in plants: *Arabidopsis* (Kover *et al.* 2009; Klasen *et al.* 2012), nested association mapping (NAM) populations (Li *et al.* 2011) in maize (Yu *et al.* 2008) and sorghum (Bouchet *et al.* 2017), and multigenerational advanced intercross (MAGIC) populations of rice (Yamamoto *et al.* 2014) and maize (Dell’Acqua *et al.* 2015). In animals, other than aforementioned prospective study of Valdar *et al.* (2006a): Noble *et al.* (2017) assessed mapping power of SNP association while introducing a 507-strain nematode resource, the *Caenorhabditis elegans* Multi-parental Experimental Evolution (CeMEE) panel; and King *et al.* (2012) estimated haplotype-based association power while introducing the *Drosophila* Synthetic Population Resource (DSPR), a fly panel with more than 1,600 lines. In a follow-up DSPR power analysis, King and Long (2017) compared the DSPR with the related *Drosophila* Genetic Reference Panel (DGRP) (Mackay *et al.* 2012). They illustrated how QTL effect size differs between a population whose allele frequencies are balanced (DSPR) vs one whose allele frequencies are less balanced (DGRP) and explored implications for cross-population validation; they also compared mapping power for bi-allelic QTL, based on single SNPs, and multi-allelic QTL constructed from actual adjacent SNPs within genes.

Here we examine related topics on QTL mapping power in the realized CC, including: 1) how power is affected by the number of strains and replicates; 2) how it is affected by the number of functional alleles and their distributions among the founders; and 3) how the QTL effect size is specific to a particular population or sample and how that influences a power estimate’s interpretation.

To allow researchers to repeat our analyses, but tailored to their own specific requirements or with updated CC genome lists, we provide an R package SPARCC (Simulated Power Analysis of the Realized Collaborative Cross), a tool that evaluates the power to map QTL by performing efficient haplotype regression-based association analysis of simulated QTL using the currently available CC genomes. SPARCC is highly flexible, allowing QTL to be simulated with any possible allele-to-founder pattern and scaled with respect to different reference populations. As a reusable resource, researchers could estimate power calculations based on the CC strains available to them and potentially incorporate prior knowledge about the genetic architecture of the likely QTL or the phenotype as whole.

## Methods

Our power calculations are based on three main processes:

1. Simulation of CC data, including selection of CC strains from a fixed set of realized CC genomes, and QTL location, and simulation of phenotypes.
2. QTL mapping, including determination of significance thresholds.
3. Evaluation of QTL detection accuracy, power and false positive rate (FPR).

These are described in detail below, after a description of the genomic data that serves as the basis for the simulations.

### Data on realized CC genomes

#### CC strains

Genome data was obtained for a set of 72 CC strains (listed in **Appendix C**) available at the time of writing from http://csbio.unc.edu/CCstatus/index.py?run=FounderProbs. Genome data was in the form of founder haplotype mosaics (see below) for each strain, this based on genotype data from the MegaMUGA genotyping platform (Morgan *et al.* 2016) applied to composites of multiple mice per strain. Since genotyping, some of the 72 strains have become extinct, and more may do so in the future (Darla Miller *pers. comm*.), although it is also possible that more may be added. At the time of writing, however, these were all genomes that had been observed by workers at UNC.

Of the 72 CC strains used in the simulations, it is planned that 54 will be maintained and distributed by The UNC Systems Genetics Core, along with another 5 whose genome data were not available in time for this study (see **Discussion**) to give a UNC total of 59 strains (listed in **Appendix C**). A subset of the UNC 59 will also eventually be maintained by The Jackson Laboratory, which will also potentially maintain 11 of the 72 not among the UNC 59.

The 72 strains used in the simulations included two that were more closely related than others: CC051 and CC059. These strains, which are among the UNC 59, were derived from the same breeding funnel; the number of independent strains available from UNC is thus arguably 58. This relatedness, though not explicitly modeled in the simulations, is nonetheless marked in the figures, which include an indicator denoting 58 as a currently realistic maximum for strain number in CC studies.

#### Reduced dataset of haplotype mosaics

The genomes of the CC, as with other MPPs, can be represented by inferred mosaics of the original founder haplotypes (Mott *et al.* 2000). Founder haplotype mosaics were inferred previously by the UNC Systems Genetics Core (http://csbio.unc.edu/CCstatus/index.py?run=FounderProbs) using the hidden Markov model (HMM) of Fu *et al.* (2012) applied to genotype calls from MegaMUGA, a genotyping platform providing data on 77,800 SNP markers (Morgan *et al.* 2016). The HMM inference provides a vector of 36 diplotype probabilities for each CC strain for each of 77,551 loci (each defined as the interval between adjacent, usable SNPs) across the genome. Rather than using all of the available data for our simulations, we used a reduced version: since adjacent loci often have almost identical descent, mapping using all loci is both computationally expensive and—at least for the purposes of the power analysis—largely redundant. Thus, prior to analysis the original dataset was reduced by averaging adjacent genomic intervals whose diplotype probabilities were highly similar. Specifically, adjacent genomic intervals were averaged if the maximum L2 norm between the probability vectors of all individuals is less than 10% of the maximum possible L2 norm 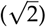; this reduced the file storage from 610 MB to 288 MB, and the genome from 77,551 to 17,900 intervals (76.9% reduction in positions to be evaluated in a scan).

### Phenotype simulation

Phenotypes for CC strains were simulated based on effects from a single QTL, plus effects of polygenic background (“strain effects”), and noise. Within our simulation framework, we specified: 1) the QTL location, which randomly was sampled from the genome; 2) the sample size in terms of both strains and replicates; 3) how the eight possible haplotypes at that location are grouped into eight or fewer functional alleles (the “allelic series”; see below); and 4) how those alleles, along with strain information, are used to generate phenotype values (see below).

#### Underlying phenotype model

Simulated phenotypes were generated according to the following linear mixed model. For given QTL with *m* ≤ 8 functional alleles, phenotype values 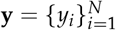 for *N* individuals in *n* ≤ *N* strains were generated so that

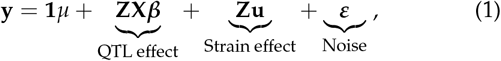

where **1** is an *N*-vector of 1’s, *μ* is an intercept, **Z** is an *N* × *n* incidence matrix mapping individuals to strains, **X** is an *n* × *m* allele dosage matrix mapping strains to their estimated dosage of each of the *m* alleles, ***β*** is an *m*-vector of allele effects, **u** is an *n*-vector of strain effects (representing polygenic background variation), and ***ε*** is an *N*-vector of unstructured, residual error. The parameter vectors ***β***, **u** and ***ε*** were each generated as being equivalent to independent normal variates rescaled to have specific variances: the strain effects **u** and residual ***ε*** were rescaled to have population (rather than sample) variances 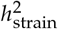 and *σ*^2^ respectively; the allele effects ***β*** were rescaled so that the QTL contributes a variance 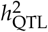, with this latter rescaling performed in one of three distinct ways (described later).

The relative contributions of the QTL, polygenic background, and noise were thus controlled through three parameters: the QTL effect size, 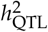, the strain effect size, 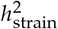, and the residual variance *σ*^2^. By convention, these were specified as fractions summing to exactly 1.

The allele dosage matrix **X** was generated by collapsing functionally equivalent haplotypes according to a specified allelic series. Let **D** be an *n* × 36 incidence matrix describing the haplotype pair (diplotype) state of of each CC strain at the designated QTL, with columns corresponding to AA,…, HH, AB, …, GH, such that, for example, {*D*}_3,1_ = 1 implies CC strain 3 has diplotype AA. Then

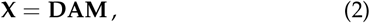

where **A** is an 36 × 8 additive model matrix that maps diplotype state to haplotype dosage (*e.g*., diplotype AA equals 2 doses of A), and **M** is an 8 × *m* “merge matrix” [after Yalcin *et al.* (2005)] that encodes the allelic series, mapping the 8 haplotypes to *m* alleles, such that if haplotypes A and B were both in the functional group “allele 1”, then diplotype AB in **D** would correspond to 2 doses of allele 1 in **X** (see examples in **Appendix D**).

#### QTL allelic series

The specification of an allelic series, rather than assuming all haplotype effects are distinct, acknowledges that for many QTL we would expect the same functional allele to be carried by multiple founder haplotypes. For our main set of simulations, the allelic series was randomly sampled from all possible configurations (examples in **Figure 1**); in a smaller, more focused investigation of the effects of allele frequency imbalance, we sampled from all possible configurations of bi-alleles.

**Figure 1.**
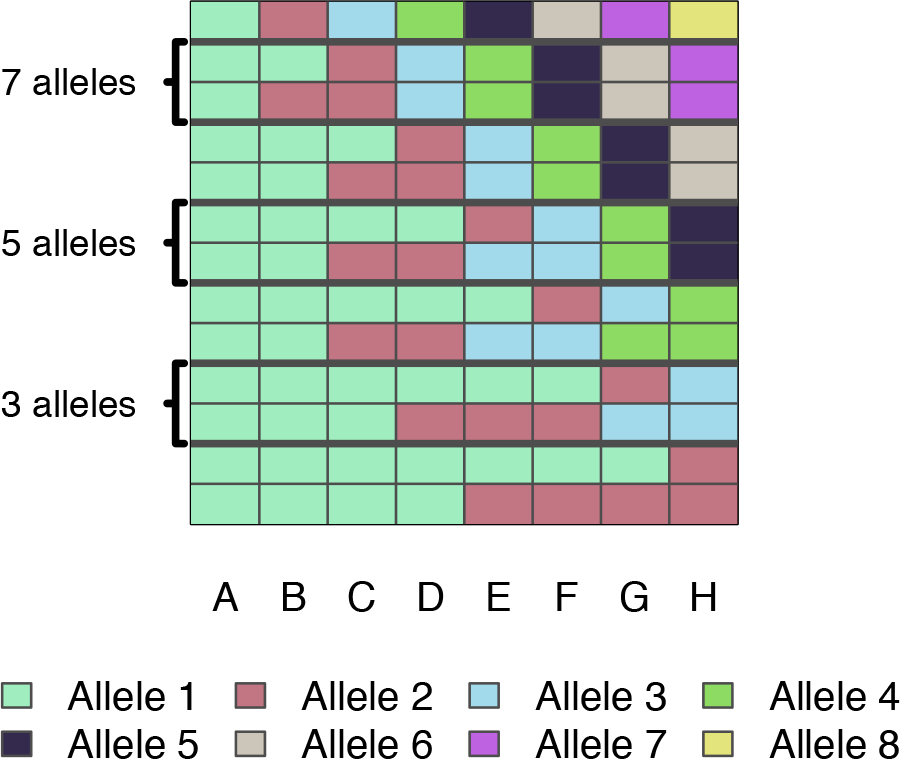
Example allelic series with differing numbers of functional alleles. Each row is an allelic series, each column of the grid is a CC founder, and colors correspond to functional allele. Two examples of allelic series are provided for each number of functional alleles: a balanced series and an imbalanced series. The entire space of allelic series are not shown here; however, the full space of series with two alleles is shown in **Figure 9A**.

#### Alternative definitions of QTL effect size: B and DAMB

The QTL effect size 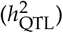 is a critical determinant of mapping power; yet its precise definition and its corresponding interpretation often varies between studies and according to what question is being asked. We used two alternative definitions, “B” and “DAMB”, described below. These alternatives acknowledge that the proportion of variance explained by a particular QTL, and thus the power to detect that QTL, is not determined solely by 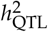, but rather depends on several additional factors, namely: the variance of the finite sample of allele effects ***β***; the allelic series configuration **M**; and the particular set of CC strains and their locus diplotypes **D**.

Definition B scales the allele effects so that 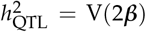, where V() denotes the population variance (rather than the sample variance). The QTL effect size is interpretable as the variance that would be explained by the QTL in a theoretical population that is balanced with respect to the functional alleles. As such, the proportion of variance explained by the QTL in the mapping population will deviate from 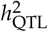 due to imbalance in both **M** and **D**. Conversely, for a given 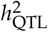, the allelic values at a QTL will be constant across populations. (Note: the 2 multiplier ensures proper scaling since **X** from Eq 2 includes dosages of founder haplotypes at the QTL, ranging from 0 to 2.)

Definition DAMB scales the QTL effect so that 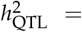 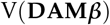. The QTL effect size is exactly the variance explained by the QTL in the mapping population, essentially the *R*^2^. As such, it depends on both **M** and **D**. Correspondingly, for a given 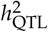, the allelic values will adjust depending on which population they are in. [In the **Supplement**, for completeness, we also describe a further, intermediate option, Definition MB, where 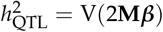, corresponding to balanced founder contributions.]

The earlier power study of Valdar *et al.* (2006a), which considered only bi-allelic QTL, defined effect size in a manner comparable to Definition B.

#### Averaging over strains and causal loci

The previous subsections described simulation of a single phenotype conditional on a set of strains and a causal genomic locus. For each of *S* simulations, *s* = 1,…, *S*, we averaged over these variables by uniformly sampling 1) the set of strains included in the experiment (for a specified number of strains), 2) the causal locus underlying the QTL, and 3) the allelic series (for a specified number of functional alleles). This was intended to produce power estimates that take into account many sources of uncertainty and are thus broadly applicable.

### QTL detection and power estimation

#### QTL mapping model

QTL mapping of the simulated data was performed using a variant of Haley-Knott (HK) regression (Haley and Knott 1992; Martínez and Curnow 1992) that is commonly used in MPP studies (Mott *et al.* 2000; Liu *et al.* 2010; Fu *et al.* 2012; Gatti *et al.* 2014; Zheng *et al.* 2015) whereby association is tested between the phenotype and the local haplotype state, the latter having been inferred probabilistically from genotype (or sequence data) and represented as a set of diplotype probabilities or, in the case of an additive model, a set of haplotype dosages then used as predictors in a linear regression. Specifically, we used HK regression on the strain means (Valdar *et al.* 2006a; Zou *et al.* 2006) via the linear model

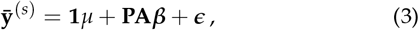

where 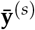 is the s^th^ simulated *n*-vector of strain means, **P** is an *n* × 36 matrix of inferred diplotype probabilities for the sampled CC genomes at the QTL [*i.e*., **P =** *p*(**D**|genotype data); see Zhang *et al.* (2014)], and ***ϵ*** is the *n*-vector of residual error on the means, distributed as 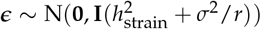. The above implies an eight-allele model (cf Eq 1 with **M** = **I**). Although this could lead to reduced power when there are fewer functional alleles, particularly at loci in which the functional alleles are not well represented, it is most common in practice, in accordance with the fact that the allelic series of an unmapped QTL would typically be unknown in advance [*e.g*., Mott *et al.* (2000); Valdar *et al.* (2006a,b); Svenson *et al.* (2012); Gatti *et al.* (2014)]. Additional factors that might contribute to variation in an experiment, such as covariates or batch effects, are neither simulated nor modeled; it is assumed that such factors would be adequately accounted for by, for instance, addition of suitable covariates, pre-processing (*e.g*., residualizing) of phenotype values or similar, and ultimately lead to a more-or-less equivalent analysis to that described here. The fit of Eq 3 was compared with that of an intercept-only null model via an F-test, and produced a p-value, reported as its negative base 10 logarithm, the logP. This procedure was performed for all loci across the genome, resulting in a genome scan for **y**^(*s*)^.

#### Genome-wide significance thresholds and QTL detection

Genome-wide significance thresholds were determined empirically by permutation. The CC panel is a balanced population with respect to founder genomic contributions and, by design, has minimal population structure. These features support the assumption of exchangeability among strain genomes: that under a null model in which the genetic contribution to the phenotype is entirely driven by infinitesimal (polygenic) effects, all permutations of the strain labels (or equivalently, of the strain means vector **y**^(*s*)^) are equally likely to produce a given configuration of **y**^(*s*)^. Permutation of the strain means, **y**^(*s*)^, was therefore used to find the logP critical value controlling genome-wide type I error rate (GWER) (Doerge and Churchill 1996). Briefly, we sampled 100 permutations and perform genome scans for each; this was done efficiently using a standard matrix decomposition approach (**Appendix A**). The maximum logPs per genome scan and simulation s were then recorded, and these are fitted to a generalized extreme value distribution (GEV) (Dudbridge and Koeleman 2004; Valdar *et al.* 2006a) using R package evir (Pfaff and McNeil 2018). The upper *α* = 0.05 quantile of this fitted GEV was then taken as the *α*-level significance threshold, 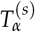. If the maximum observed logP for **y**^(*s*)^ in the region of the simulated QTL exceeded 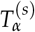, then the corresponding locus was considered to be a (positively) detected QTL (see immediately below).

#### Performance evaluation

For a given simulation, we declared a true positive if the detected QTL was within ±5Mb of the true (simulated) QTL. The 5Mb window size was used to approximate a QTL support interval, which is partly a function of linkage disequilibrium (LD) in the CC. (LD has been characterized in the CC previously but not summarized with a single point estimate (Collaborative Cross Consortium 2012); our choice of 5Mb is therefore an approximation, but we find that it only marginally increased mapping power relative to using smaller window widths.) A false positive was declared if one or more QTL were detected on chromosomes other than the chromosome harboring the simulated QTL. Simulations in which a QTL was detected on the correct chromosome but outside the 5Mb window were disregarded; although this was potentially wasteful of data and biased FPR slightly downward due to loss of false positives on the chromosome with the simulated QTL, it avoided the need for arbitrary rules to handle edge cases in which it was ambiguous whether the simulated signal had been detected or not. Power for a given simulation setting was then defined as the proportion of true positives among all simulations at that setting, and the FPR was defined as the proportion of false positives.

As a measurement of mapping resolution, for true positive such that any permutation of the values is equally strain likely (the detection, we recorded the mean and the 95% quantile of the genomic distance from the true QTL. Given our criterion for calling true positives, the maximum distance was necessarily 5Mb, and experimental settings that correspond to low power would be expected to have fewer data points, yielding estimates that are unstable. In order to obtain more stable estimates, we used a regularization procedure, estimating the mean distance and 95% quantiles as weighted averages of the observed values and prior pseudo-observations. Specifically, for an arbitrarily small but detected true positive QTL, it is reasonable to expect the peak signal to be distributed uniformly within the ±5Mb window. This implies a mean location error of 2.5Mb and a 95% quantile of 4.75Mb. Thus, when calculating the regularized mean location error we assumed 10 prior pseudo-observations of 2.5Mb, and when calculating the regularized 95% quantile we assume 10 prior pseudo-observations of 4.75Mb. This number of pseudo-observations represents 1% of the maximum number of possible data points.

### Overview of the simulations

#### Simulation settings

Simulations for all combinations of the following parameter settings:

- Number of strains: [(10-70 by 5), 72]
- QTL effect size (%): [1, (5-95 by 5)]
- Number of functional alleles: [2, 3, 8]

The number of observations per strain were fixed at *r* = 1 and the background strain effect size was fixed at 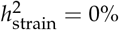 with the understanding that results from these simulations provide information on other numbers of replicates and strain effect sizes implicitly. Specifically, a simulated mapping experiment on strain means that assumes *r* replicates, strain effect 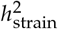, and QTL effect size 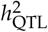 is equivalent to a single-observation mapping experiment with no strain effect and QTL effect size 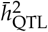, where

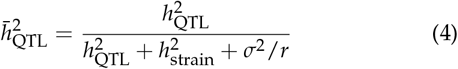

[Valdar *et al.* (2006a), after Soller and Beckmann (1990); Knapp and Bridges (1990); Belknap (1998)]. For example, a mapping experiment on strain means with QTL effect size 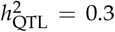,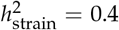, *σ*^2^ = 0.3, and *r* = 10, is equivalent to our simulation of a single-observation with no strain effect but QTL effect size 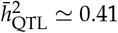 (**Supplement**).

We conducted *s* = 1, 000 simulation trials per setting. CC strains and the position of the QTL were sampled for each simulation, providing estimates of power that are effectively averaged over the CC population. We ran these settings for QTL effect sizes specified with respect to the observed mapping population (Definition DAMB) and a theoretical population that is balanced in terms of the functional alleles (Definition B). Confidence intervals for power were calculated based on Jeffreys interval (Brown *et al.* 2001) for a binomial proportion. A description of the computing environment and run-times are provided in **Appendix B**.

### Examining FPR when accounting for non-exchangeability of CC strain genomes

In the simulations and mapping procedures described above, strain effects are modeled under the assumption that all CC strains are (at least approximately) equally related. That is, the effects **u** = *u*_1_,…, *u*_72_ in Eq 1 are simulated as 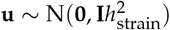 such that any permutation of the values is equally likely (the effects are exchangeable), and this same assumption is made in both the mapping model of Eq 3 and the permutation-based estimation of significance thresholds.

An assumption of equal relatedness among CC strains is commonplace: it is suggested by the exchangeable random funnel design used in the CC, is supported by the results of Valdar *et al.* (2006a), and has been made in every CC or pre-CC mapping analysis to our knowledge. Making this assumption simplifies QTL mapping analysis by obviating the need for an explicit modeling of genomic similarity [as in, *e.g*., Kang *et al.* (2008)], since, when those similarities are approximately equal and the analysis is performed on strain means, the strain effects are absorbed into the residual error.

Nonetheless, CC strains are equally related only in expectation. Much like the “equal” relatedness of siblings, realized relatedness will depart from expectation due to chance at the point of mixing, and, in the case of the CC, due to selection [*e.g*., arising from male sterility (Shorter *et al.* 2017)] and genetic drift during inbreeding [as reflected in unequal founder contributions by Srivastava *et al.* (2017)]. This combination of stochastic forces can produce unequal relatedness, correlated effects among strains, and population structure, at least at some level.

To quantify population structure in the realized CC, we compared the eigenvalues of the realized genetic relationship matrix **K**, calculated from the founder mosaic probabilities [after Gatti *et al.* (2014)], with those from an idealized **K** that reflects equal relatedness of the CC strains, whose off-diagonal elements were set to the mean value observed for the off-diagonal elements in the realized **K**. We observed that slightly fewer principal components are required to explain 95% of the variation in the realized **K** than are required for the balanced K (64 vs 68 components, respectively; **Figure S5A**). This reduction was attenuated with the omission of CC059, one of the two cousin strains, but not completely (64 vs 67 components; **Figure S5B**). This suggested that the realized CC strains have mild population structure.

To evaluate to what degree the population structure in the realized CC genomes could inflate FPR when mapping using an analytic model and threshold procedure that ignores it (*i.e*., that assumes exchangeability), we performed an additional set of null simulations in which strain effects were generated according to additive infinitesimal model (Lynch and Walsh 1998) based on the actual genomic similarities. Specifically, we set 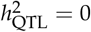 and 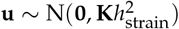 but left our mapping protocol unchanged. We conducted 10,000 such null simulations with r=1 for each setting of strain effect size (%): [0-100 by 20]. These simulations were performed using either all 72 founder strains or 71 strains with the omission of CC059, one of the two highly-related cousin strains. A false positive was declared if any QTL were detected based on the permutation-based significance threshold.

### Measuring the Beavis effect

The “Beavis effect” (Beavis 1994) refers to an upward bias in estimated effect sizes for detected QTL. This phenomenon, also known as the “winner’s curse” (Zollner and Pritchard 2007), arises because the data used for effect estimation has already been substantially selected during QTL discovery; the resulting (post-selection) estimates are thus inflated due to ascertainment bias. The Beavis effect was evaluated theoretically in Xu (2003) and found to be most pronounced in studies of smaller sample size (*n <* 100), suggesting that it could be a significant feature of CC mapping studies.

To assess the extent of the Beavis effect in CC mapping experiments, we performed simulations (*s =* 1, 000) mapping a bi-allelic QTL, with one replicate (*r =* 1) and zero background strain effect 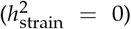 for all combinations of simulated QTL effect size under Definition DAMB 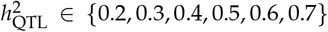 and numbers of strains *n* ∈ {40, 50, 60, 72}. If an association was detected within the 10Mb window (using permutation-based thresholds as above), then we recorded the QTL effect size as the *R*^2^ of the model fit at the peak locus (which may or may not be the locus at which the QTL was simulated).

### Availability of data and software

#### R package

All analyses were conducted in the statistical programming language R (R Core Team 2018). SPARCC is available as an R package on GitHub at https://github.com/gkeele/sparcc. Specific arguments that control the phenotype simulations, the strains used, genomic position of simulated QTL, and allelic series, are listed in the **Supplement**. A static version of SPARCC is also provided there (File S2).

Also included within the SPARCC R package are several results datasets. These include data tables of power summaries from our simulations, as well as table summaries from simulations of a bi-allelic QTL that is balanced in the founders, maximally unbalanced in the founders, and the distance between detected and simulated QTL. Further details are provided in File S1 of the **Supplement**, an account of all the supplemental files. These files are available at figshare, including data, and scripts to run the analysis and produce the figures. File S3 contains the founder haplotype mosaics required for the SPARCC package. Files S4, S5, and S6 can be used to perform the large-scale power analysis. File S7 describes options in the SPARCC package, and also provides two simple tutorials. File S8 produces the figures in this paper and **Supplement**. File S9 is the supplemental tables and figures.

#### CC strains

The 72 CC strains with available data that were included in the simulations are described in **Appendix C**. Founder diplotype probabilities for each CC strain are available on the CC resource website (http://csbio.unc.edu/CCstatus/index.py?run=FounderProbs). We used probabilities corresponding to build 37 (mm9) of the mouse genome, though build 38 (mm10) is also available at the same website.

We store the founder haplotype data in a directory structure that SPARCC is designed to use, and was initially established by the HAPPY software package (Mott *et al.* 2000). The reduced data are available on GitHub at https://github.com/gkeele/sparcc_cache.

## Results

Power simulations were performed for varying numbers of strains, replicates and functional alleles, and for a ladder of QTL effect sizes. QTL effect size was defined in two ways: as the variance explained in a hypothetical populations that is balanced with respect to the alleles (Definition B; see **Methods**), or as the variance explained in the realized population (Definition DAMB). In this section we focus on results using the first of these, Definition B, owing to its more consistent theoretical interpretation. Under that definition, plots of power against numbers of strains are shown in **Figure 3**, and power across a representative selection of conditions is shown in **Table 1**. For comparison, these numbers are also provided for simulations under Definition DAMB in **Table S1**. Throughout these simulations the false positive rate was controlled at the target 0.05 level (**Figure S2**).

**Figure 2.**
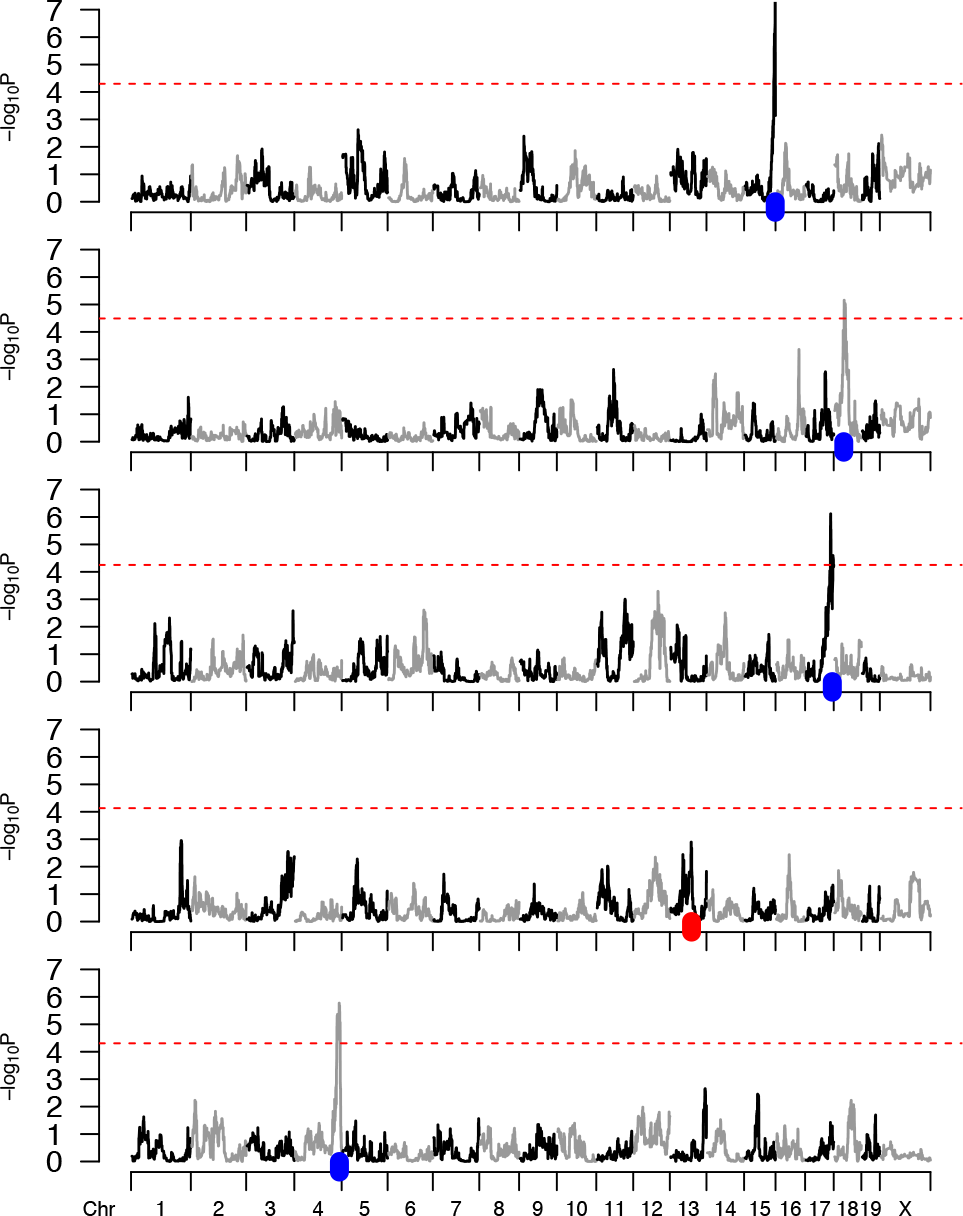
Simulated CC data and resulting genome scans. Five simulated genome scans are generated by the code provided in a simple example using our package SPARCC. Red dashed lines represent 95% significance thresholds based on 100 permutation scans. A blue tick represents the simulated position for a QTL that was successfully detected, whereas a red tick marks a QTL that was missed. These simulations were based on a specified set of 65 CC strains, five replicates of each strain, two functional alleles, 10% QTL effect size, and no background strain effect. The QTL is not mapped in the fourth simulation, ranked top to bottom, resulting in a power of 80%. Actual power calculations are based on a greater number of simulations.

### Large effect QTL usually detected by 50 or more strains

As a baseline for describing mapping power in the CC, an experiment using one replicate (*r* = 1) of all 72 strains is well-powered to detect QTL explaining >40% of phenotypic variance but moderately or low powered for QTL explaining 30% or less (**Table 1**). Specifically, assuming eight functional alleles, there is 96.4% power to detect a 50% QTL, 79.2% for a 40% QTL, 44.1% for a 30% QTL, and 12.4% for a 20% QTL.

More broadly, simulations across different allele effect types and numbers of strains showed that studies without replicates and with large numbers of strains (>50) were found to be well-powered to detect large effect QTL (>40%) (Figure 3 **[top]**).

Identifying smaller effect QTL is feasible, however, using replicates. Replicates improve power by reducing the individual noise variance; as such the extent of the power improvement diminishes as more variance is attributable to background strain effects than noise. Assuming no background strain effect, and using 50 strains, the power to detect a 20% effect-size QTL with a single replicate is near zero; with 5 replicates it approaches 80%; detecting QTL with effect sizes ≤ 10% is challenging. For example, achieving 80% power to detect an effect size of 10% when all 72 CC strains were used required more than 5 replicates per strain (Figure 3 **[middle right]**). Assuming a background strain effect, as would be expected with a complex trait, can reduce the QTL mapping power of small effect QTL substantially (Figure 3 **[bottom]**).

### Additional strains improve power more than additional replicates

We investigated the relationship between power and the total number of mice, evaluating whether power gains were greater with additional CC strains or additional replicate observations. Power was interpolated over a grid of values for number of replicates and total number of mice from simulations based on a single observation per strain (**Figure 5**). This showed that additional CC strains improved mapping power more than additional replicates; this is indicated by higher power values for lower numbers of replicates while holding number of mice constant (see **Figure 5**, bordered vertical section at 250 mice).

### Location error of detected QTL

To obtain an approximation of mapping resolution, for all true positive detections we recorded the location error, or the genomic distance between simulated and detected QTL. The mean and the 95% quantile of the location error are reported as stabilized estimates for different numbers of strains and QTL effect sizes, but averaged over all other conditions, in **Figure 4**. (The stabilization procedure is described in **Methods**; raw, unstabilized estimates provided **Figure S3**.) The location error statistics require careful interpretation: for a detection to be classed as a true positive it had to be within 5Mb of the simulated QTL; therefore, location error was artificially capped at 5Mb. Mediocre performance thus corresponds to when that location seems uniformly (and therefore arbitrarily) distributed over the ±5Mb interval, that is, having a mean of 2.5Mb and a 95% quantile of 4.8Mb.

Location error was improved (reduced) by increasing the number of strains, increasing the QTL effect size, or both. In particular, as with power, location error was improved by increasing the number of strains even when while holding the total number of mice constant (**Figure S4**), consistent with mapping resolution being improved by an increased number of recombination events in the QTL region. Distributions of raw location error, stratified by levels of the number of strains, the number of functional alleles, and the QTL effect size can be found in **Figure S6**.

### False positive rate

The FPR for the QTL power simulations was estimated as the percentage of scans (per setting) that produced a statistically significant signal on a chromosome without a QTL, shown in **Figure S2**. As expected, FPR was not elevated from 5% when the strain effects were simulated independently, as the effects were exchangeable by construction. The FPR did not vary with the number of strains or the number of alleles.

In additional null simulations that where strain effects were correlated due to realized genomic similarity, QTL scans assuming independent strain effects (and thus, exchangeability) had elevated FPR (**Figure 6** and **Table S2**). Using all 72 CC strains, the FPR varied from a maximum of 14.5% when strain effects explain all variability to the well controlled FPR of 5.5% when the strain effects were relatively small. Omitting CC059, one of the highly-related cousin strains (CC053 and CC059), because of its obvious violation of equal relatedness, reduced the FPR, although it was still elevated (12.9% for maximum strain effect). This demonstrates that, when strain effects are large relative to individual error (i.e. highly heritable trait, or the use of many replicates), failure to account for population structure due to realized imbalance in founder contributions can increase the risk of false positives.

### Beavis effect

It is an expected feature of QTL mapping studies that estimates of QTL effect size, when calculated only for detected QTL, will be biased upwards. This phenomenon, known as the Beavis effect, is a form of selection bias and as such is expected to be most extreme under low power conditions, *e.g*., when detection rates are low and/or estimates have high variance.

We explored the Beavis effect in our simulations. Assuming a one-replicate (*r* = 1) experiment, we found that, for example, the estimated effect size of a simulated 20% QTL was inflated by 3-fold when mapping in 40 CC strains, and by 2-fold when mapped in 72 CC strains. More generally, and as expected, the Beavis effect was reduced with larger numbers of strains and larger QTL effect sizes (**Figure 7**).

These results also imply that the Beavis effect is reduced by replication, at least to the extent that replication boosts effective QTL effect size. For example, consider again the mapping of a 20% QTL effect in 40 strains, which with *r* = 1 replicates implies 3-fold effect size inflation. Although this inflation could be reduced to 2-fold by increasing the number of strains to 72, the same reduction could be achieved by replication: assuming no background strain effect, increasing replicates to a theoretical *r* = 1.8 (so as to give a total sample size of *N* = 40 × 1.8 = 72) would boost the QTL effect size to an effective ≈31% (according to Eq 4) and, as shown in **Figure 7**, have approximately the same result. The ability of replicates to reduce the Beavis effect, however, will diminish to the extent that there is a significant background strain effect, following the general relationship of replicates and QTL effect size described in Eq 4.

### Allele frequency imbalance reduces power

For a fixed set of QTL allele effects, it is expected that power will always be greatest when allele frequencies are balanced. Accordingly, when QTL effect size was defined in terms of the variance that would be explained in a theoretical population with balanced allele frequencies (Definition B), deviations from balance in the mapping population—either from imbalance in functional alleles among the founders or imbalance of the founders among the CC strains—inevitably reduce power (**Figure 8A**). This reduction in power under Definition B is most evident for bi-allelic QTL (pink), in which the potential imbalance in allelic series is most extreme, namely when a single founder carries one functional allele and the other seven possess the alternative allele (7v1).

**Figure 3.**
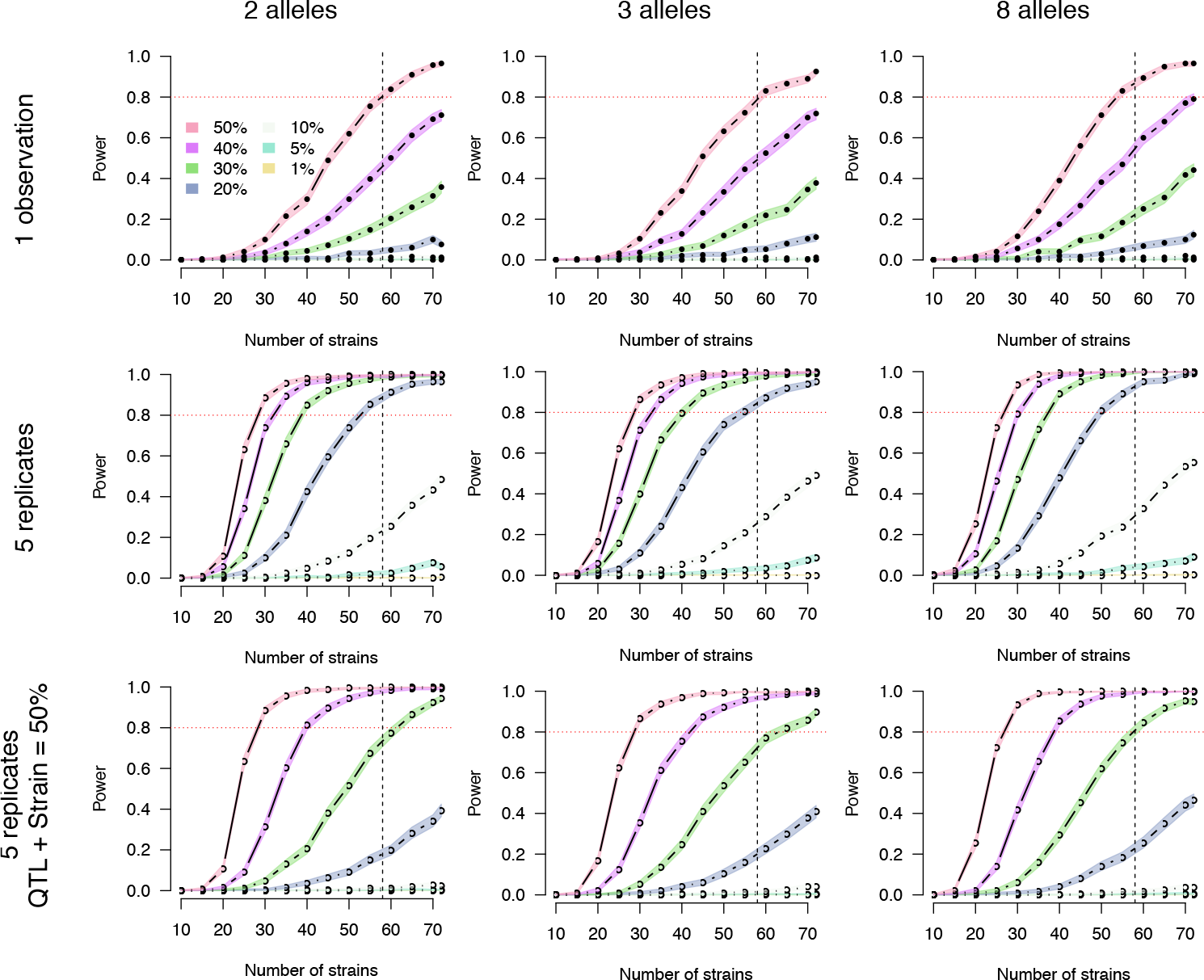
Power curves by number of CC strains. Results are stratified by a number of replicates, background strain effect size, and the number of functional alleles. The **[top]** row is based on a single observation per strain and no background strain effect. The **[middle]** row corresponds to five replicates per strain and no background strain effect. For the **[bottom row]**, five replicates are observed and the QTL effect size and background strain effect size sum to 50%, thus penalizing smaller QTL more harshly. The horizontal red dotted line marks 80% power. The vertical black dashed line marks 58 strains, which is currently the number of unrelated strains available from UNC. The columns, left to right, correspond to two, three, and eight functional alleles. Closed circles represent power estimates that were directly assessed, whereas open circles were interpolated. Simulations are based on Definition B.

**Table 1.** QTL mapping power in the Collaborative Cross based on QTL effect sizes in a balanced population (Definition B)

| QTL |  |  | Power |  |  |  |  |  |  |  |  |
| --- | --- | --- | --- | --- | --- | --- | --- | --- | --- | --- | --- |
|  |  |  | 30 strains |  |  | 50 strains |  |  | 72 strains |  |  |
| 1 obs<sup>a</sup> | 3 rep<sup>b</sup> | 5 rep<sup>b</sup> | 2 alleles | 3 alleles | 8 alleles | 2 alleles | 3 alleles | 8 alleles | 2 alleles | 3 alleles | 8 alleles |
| 0.01 | 0.003 | 0.002 | 0.001 | 0.000 | 0.000 | 0.000 | 0.001 | 0.001 | 0.001 | 0.000 | 0.000 |
| 0.05 | 0.017 | 0.010 | 0.001 | 0.001 | 0.002 | 0.004 | 0.000 | 0.001 | 0.007 | 0.000 | 0.003 |
| 0.1 | 0.036 | 0.022 | 0.001 | 0.001 | 0.001 | 0.006 | 0.003 | 0.004 | 0.013 | 0.013 | 0.014 |
| 0.15 | 0.056 | 0.034 | 0.001 | 0.003 | 0.002 | 0.009 | 0.011 | 0.014 | 0.035 | 0.054 | 0.041 |
| 0.2 | 0.077 | 0.048 | 0.006 | 0.009 | 0.003 | 0.032 | 0.026 | 0.030 | 0.077 | 0.110 | 0.124 |
| 0.25 | 0.100 | 0.062 | 0.002 | 0.011 | 0.015 | 0.076 | 0.061 | 0.066 | 0.207 | 0.231 | 0.252 |
| 0.3 | 0.125 | 0.079 | 0.011 | 0.014 | 0.010 | 0.105 | 0.118 | 0.116 | 0.357 | 0.377 | 0.441 |
| 0.35 | 0.152 | 0.097 | 0.018 | 0.024 | 0.034 | 0.194 | 0.207 | 0.261 | 0.553 | 0.564 | 0.633 |
| 0.4 | 0.182 | 0.118 | 0.035 | 0.038 | 0.056 | 0.298 | 0.335 | 0.383 | 0.711 | 0.717 | 0.792 |
| 0.45 | 0.214 | 0.141 | 0.048 | 0.063 | 0.078 | 0.456 | 0.467 | 0.539 | 0.858 | 0.857 | 0.905 |
| 0.5 | 0.250 | 0.167 | 0.098 | 0.102 | 0.114 | 0.620 | 0.630 | 0.712 | 0.964 | 0.924 | 0.964 |
| 0.55 | 0.289 | 0.196 | 0.156 | 0.180 | 0.208 | 0.789 | 0.784 | 0.860 | 0.977 | 0.961 | 0.993 |
| 0.6 | 0.333 | 0.231 | 0.272 | 0.251 | 0.304 | 0.914 | 0.896 | 0.935 | 0.990 | 0.984 | 0.998 |
| 0.65 | 0.382 | 0.271 | 0.387 | 0.412 | 0.486 | 0.953 | 0.934 | 0.985 | 0.993 | 0.992 | 0.999 |
| 0.7 | 0.438 | 0.318 | 0.603 | 0.582 | 0.635 | 0.983 | 0.965 | 0.994 | 0.998 | 0.993 | 1.000 |
| 0.75 | 0.500 | 0.375 | 0.780 | 0.746 | 0.818 | 0.990 | 0.986 | 0.999 | 0.998 | 0.999 | 1.000 |
| 0.8 | 0.571 | 0.444 | 0.890 | 0.851 | 0.923 | 0.995 | 0.991 | 1.000 | 0.999 | 1.000 | 1.000 |
| 0.85 | 0.654 | 0.531 | 0.932 | 0.927 | 0.983 | 0.997 | 0.995 | 0.999 | 1.000 | 1.000 | 1.000 |
| 0.9 | 0.750 | 0.643 | 0.970 | 0.955 | 0.994 | 0.999 | 0.999 | 1.000 | 1.000 | 0.999 | 1.000 |
| 0.95 | 0.864 | 0.792 | 0.976 | 0.966 | 1.000 | 0.999 | 0.998 | 1.000 | 1.000 | 1.000 | 1.000 |

<sup>a</sup> Convert QTL effect sizes from experiments with replicates to mean scale with Eq 4.
<sup>b</sup> Based on no background strain effect.

Conversely, when the QTL effect size is defined in terms of variance explained in the mapping population (Definition DAMB, which is similar to an *R*^2^ measure), power remains constant across different allelic series and degrees of balance. Although note that this definition carries with it the (possibly unrealistic) implication that allele effects vary depending what population they are in.

When averaged over many allelic series, QTL mapping power based on Definition B is reduced relative to Definition DAMB, with the greatest reduction occurring for bi-allelic QTL (**Figure 8B**). Though this modest reduction in power may seem to suggest that simulating with respect to a balanced population (Definition B) versus the mapping population (Definition DAMB) is unimportant in terms of designing a robust mapping experiment in the CC, we reiterate the value of using Definition B. Specifically, simulating with respect to Definiton DAMB is overly optimistic regarding mapping power for QTL with imbalanced allelic series.

We performed additional simulations to evaluate bi-allelic QTL in more detail, these being more prone to drastic imbalance under Definition B. All 127 possible bi-allelic series are visualized as a grid in **Figure 9A**, ordered from balance and high power to imbalance and low power. The corresponding power estimates are shown in **Figure 9B**. Power was maximized when the bi-allelic series is balanced (4v4; 35/127 possible allelic series) and minimized when imbalanced (7v1; 8/127 possible allelic series). Uniform sampling of bi-allelic series, the approach in the more general simulations described earlier, slightly reduced power relative to balanced 4v4 allelic series due to averaging over many cases of balance and some cases of extreme imbalance. These latter, more focused simulations highlight the extent that the reduction in QTL effect size, and thus mapping power, when simulating based on Definition B, is highly dependent on the allelic series. This could be of particular importance when considering QTL that result from a causal variant inherited from a wild-derived founder, such as CAST, which will present as both imbalanced and bi-allelic.

## Discussion

Now that the CC strains have been largely finalized, it is possible to investigate more deeply how, in potential mapping experiments, power is affected by factors such as the number of strains, the number of replicates, and the allelic series at the QTL. We find that the CC can powerfully map large effect QTL (≥ 50%) with single observations of > 60 strains. Through the use of replicates, the power to map QTL can be greatly improved, potentially mapping QTL ≥ 20% in 60 strains with 5 replicates per strain with no background strain effect. To guide the design of new CC experiments, we provide broad power curves and tables in **Figure 3** and **Tables 1** and **S1**.

**Figure 4.**
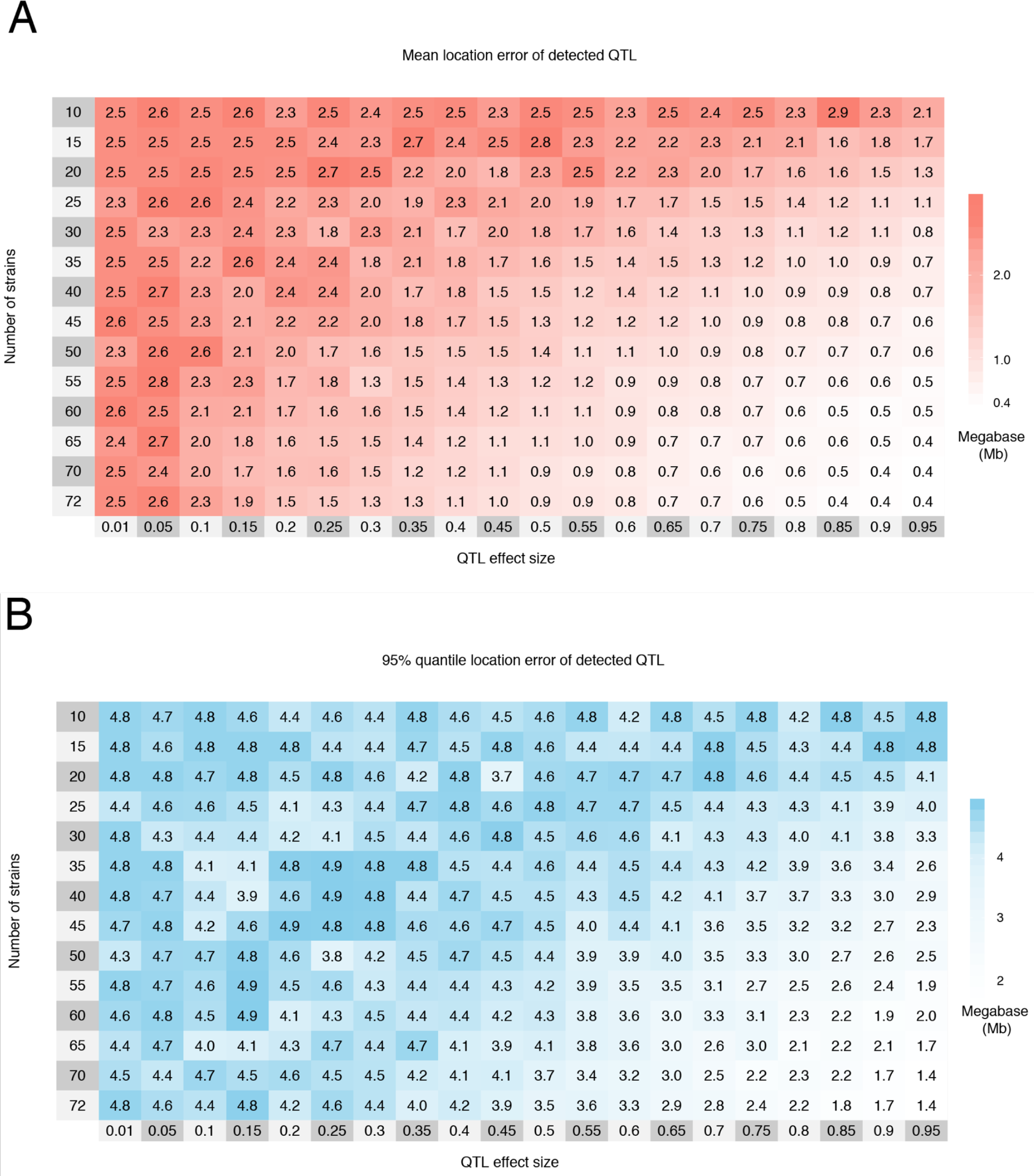
The mean (A) and 95% quantile (B) of location error, the distance in Mb between the detected and simulated QTL, by effect size and number of strains for 1,000 simulations of each setting. The simulations are based on Definition B with an eight allele QTL, and only a single observation per strain. Cells are colored red to white with decreasing mean and blue to white with decreasing 95% quantile. Regularization of the means and 95% quantile was accomplished through averaging the observed results with pseudo-counts; see **Figure S3** for the raw measurements. Increasing the number of strains reduces the location error, both in terms of the mean and 95% quantile, more so than QTL effect size, also shown in **Figure S6**. The maximum possible location error was 5Mb due to the 10Mb window centered around the true QTL position used for detecting QTL.

**Figure 5.**
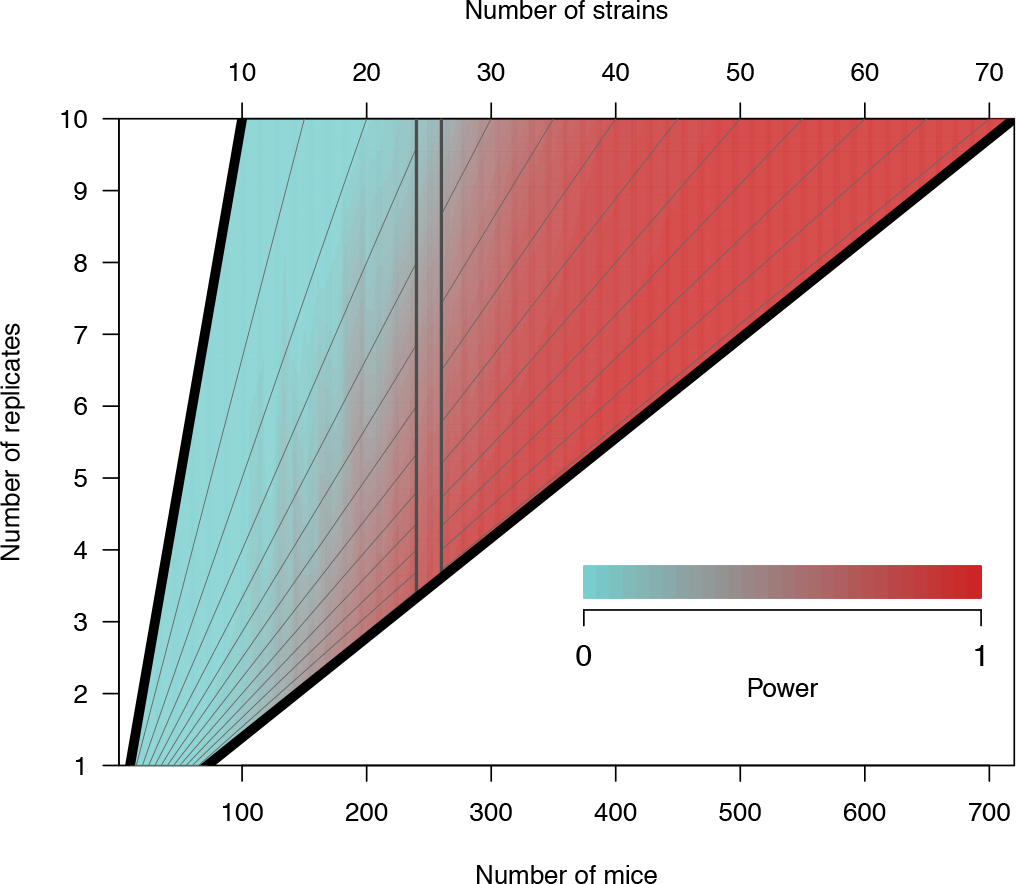
Heatmap of QTL mapping power by number of replicates and total number of mice in the experiment. Power is based on a QTL effect size of 20%, no background strain effect, and two functional alleles, though varying these parameters does not affect the dynamic between number of strains and replicates. The gray diagonal lines represent fixed values of the number of CC strains, ranging from 10 to 70 in intervals of five. Holding the total number of mice fixed, power is reduced as the percentage of the sample that are replicates is increased. This is illustrated with a cutout band centered on 250 mice, where power is lower at the top of the band when replicate mice are a relatively higher proportion of the total number of mice.

**Figure 6.**
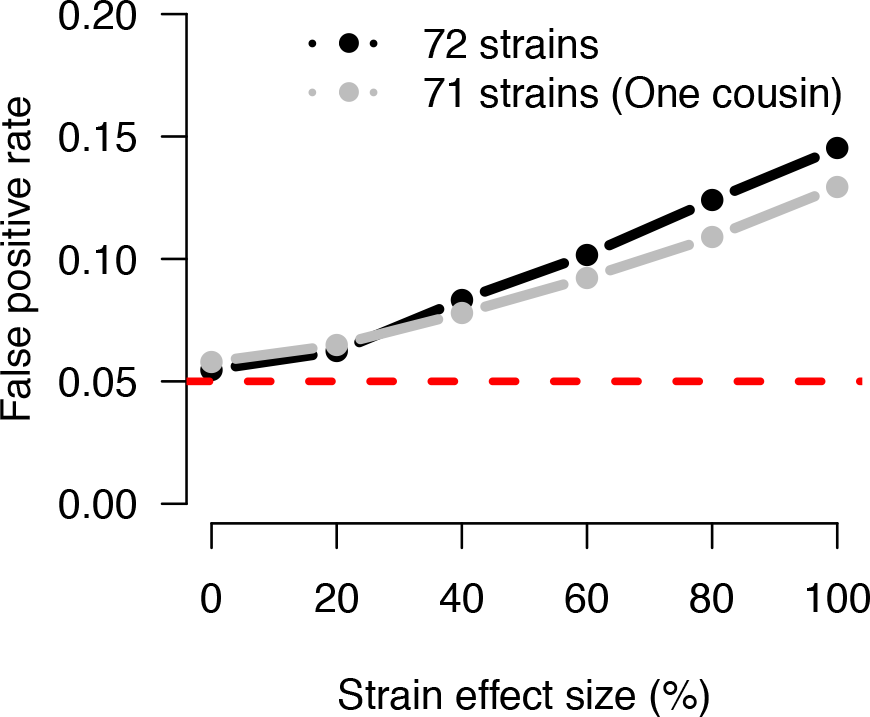
The FPR increases due to population structure among the realized genomes of the CC strains in the presence of a background strain effect and no QTL. Curves are based on 10,000 simulations for each setting of strain effect and strain sample, based on a single observation per strain. The inflation in FPR is greater for all 72 CC strains, which includes two closely related cousin strains (CC051 and CC059). Removing CC059 reduces the inflation in FPR (gray line). The dashed red line marks the specified type I error rate of 0.05, which is approximately met as expected when no strain effect is simulated, as in **Figure S2**. **Table S2** reports the specific FPR values.

**Figure 7.**
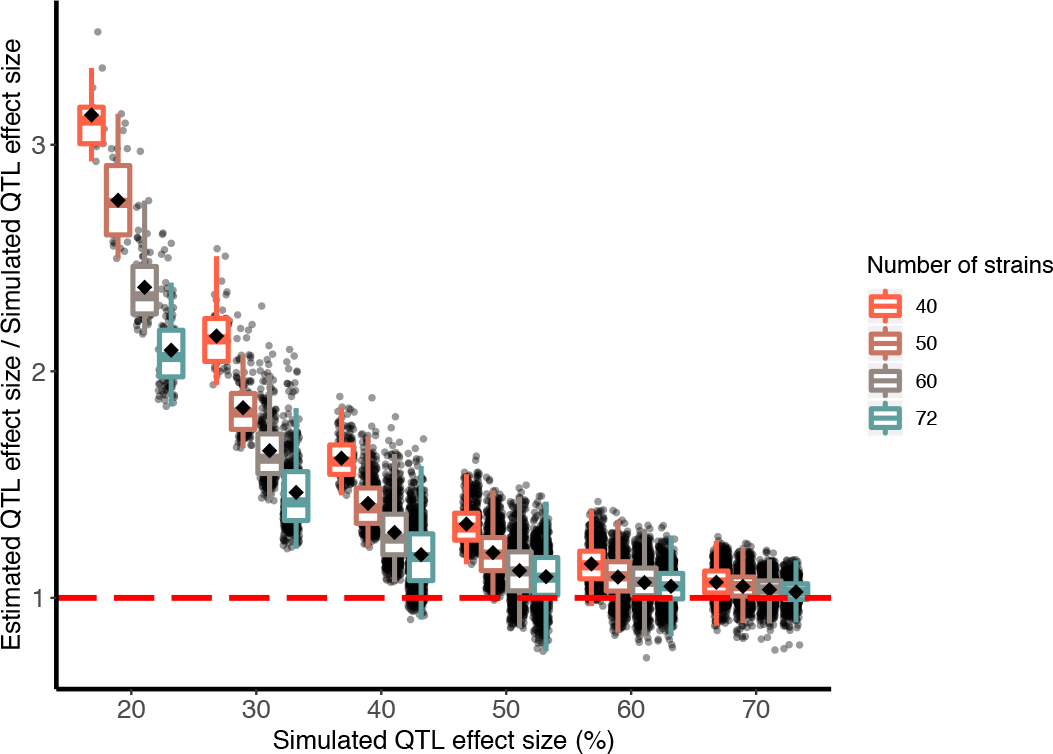
The Beavis effect (inflation of QTL effect size estimates) is more pronounced with smaller simulated QTL effect sizes and reduced numbers of strains. For different settings of numbers of strains (40, 50, 60, 72) and simulated QTL effect sizes (20%, 30%, 40%, 50%, 60%, 70%), black dots plot the ratio of the estimated effect size at a detected QTL peak to the effect size that was simulated at the true QTL locus. Out of 1,000 simulations under each setting, only successful detections are shown. Black diamonds represents the mean ratio for a category; horizontal red dashed line marks a ratio of 1, when QTL effect size estimates are unbiased (*i.e*., no Beavis effect).

**Figure 8.**
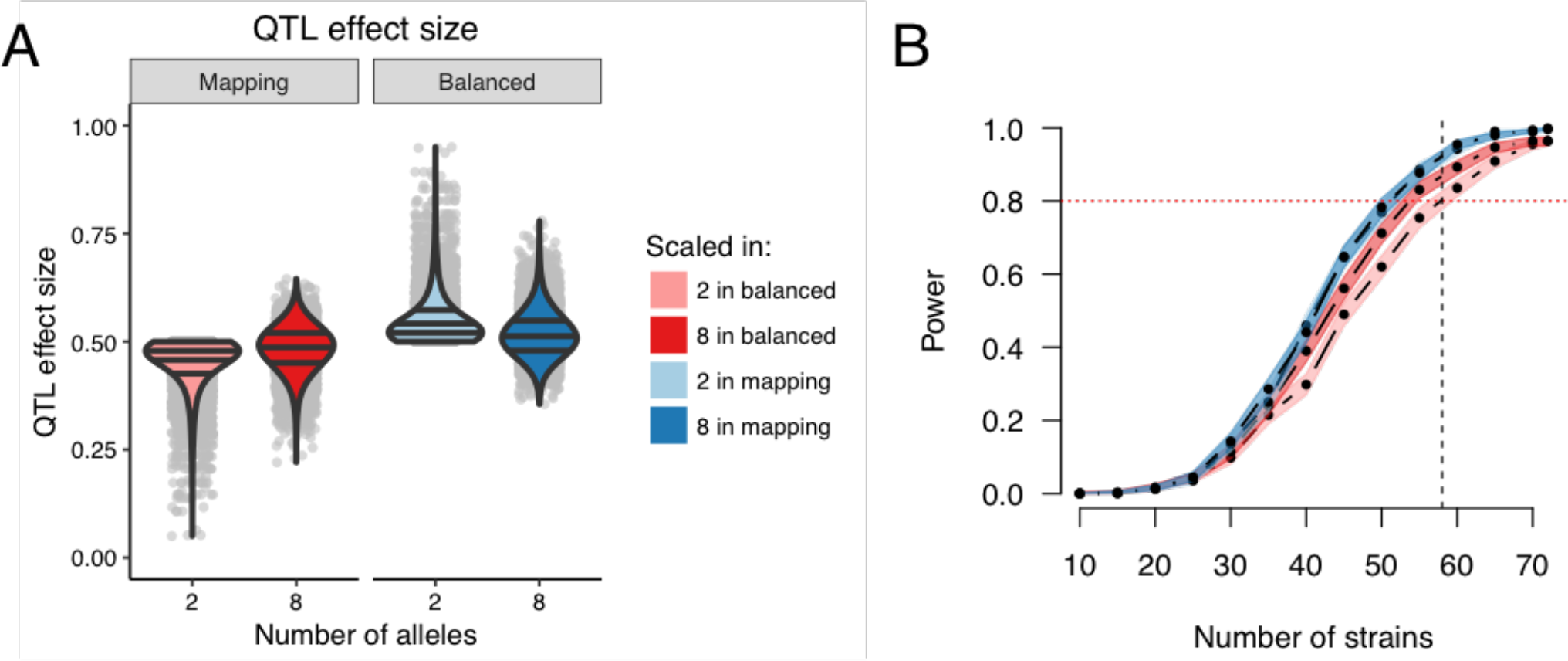
QTL effect sizes are in reference to a population, though effect size in the specific mapping population will determine the mapping power. Consider two populations as examples: the mapping population (definition DAMB) and a population balanced in the functional alleles (definition B). (A) QTL effect size distributions based on 10,000 simulations of the QTL for 72 strains. Using definition B, the effect sizes for the mapping population for two alleles is pink and eight alleles is red. Using definition DAMB, the effect sizes in the balanced population for two alleles is light blue and eight alleles is dark blue. Horizontal lines within the violin plots represent the 25^th^, 50^th^, and 75^th^ quantiles from the estimated densities. Gray dots represent actual data points. (B) Power curves corresponding to the previously described settings of alleles and QTL effect size definitions. Power curves are estimated from 1,000 simulations per number of strains for a 50% QTL, no background strain effect, and a single observation per strain. The horizontal red dotted line marks 80% power. The vertical black dashed line marks 58 strains, which is currently the number of unrelated strains available from UNC.

**Figure 9.**
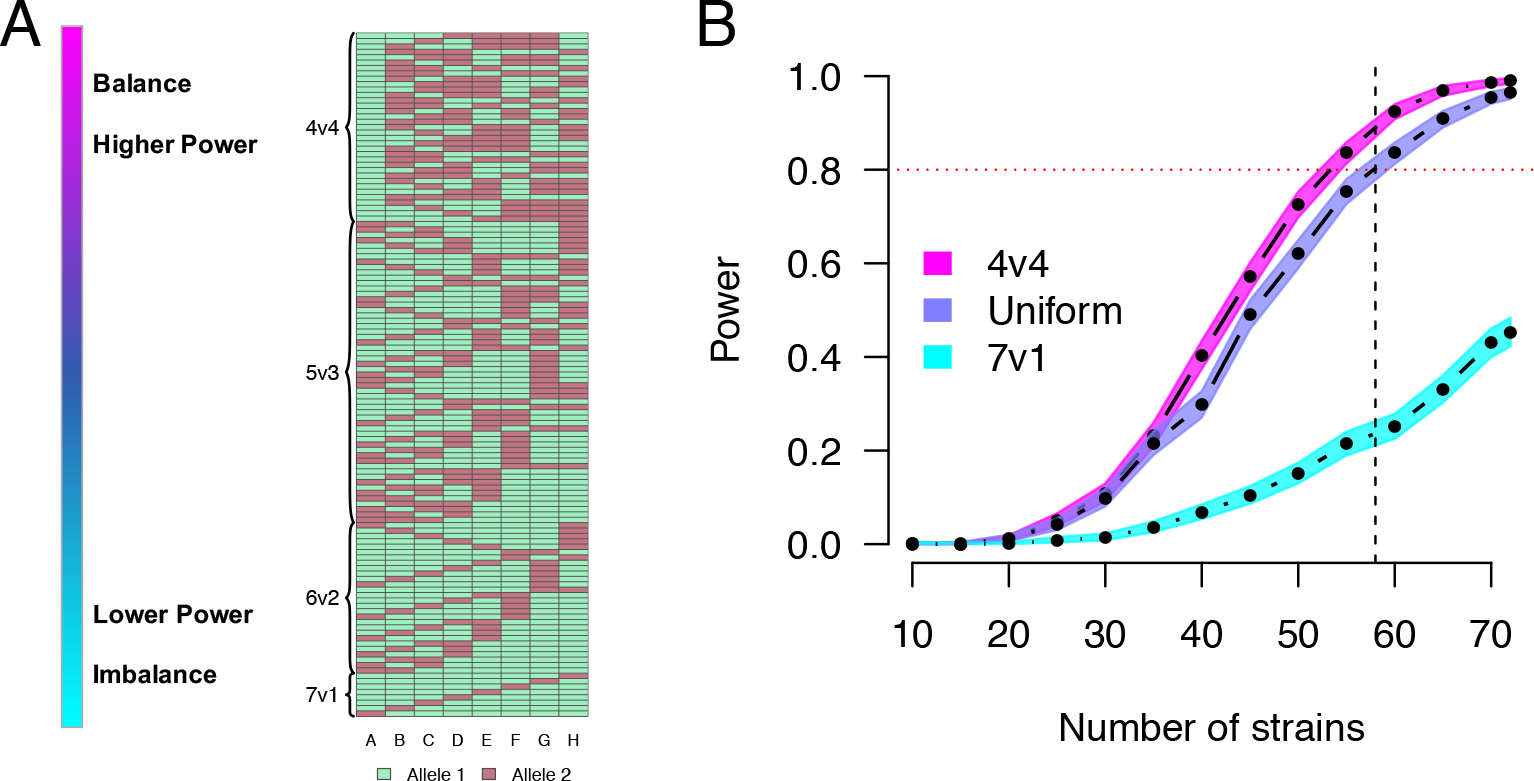
The balance of the allelic series for QTL with two functional alleles, and its effect on QTL mapping power. (A) The 127 possible allelic series for a bi-allelic QTL, categorized by the balance in the distribution of alleles among the CC founder strains, and ordered with balanced allelic series at the top and imbalanced at the bottom. (B) Power curves comparing three different sampling approaches for the allelic series with two functional alleles, for populations simulated to have a QTL effect size of 50% in a balanced theoretical population, with a single observation per CC strain. The horizontal red dotted line marks 80% power. The vertical black dashed line marks 58 strains, which is currently the number of unrelated strains available from UNC.

The power calculations described here take advantage of realized CC genomes, allowing the power estimates to be highly specific to the available strains but also necessarily restricting the number that can be used. This differs from the simulations of Valdar *et al.* (2006a), which primarily focused on comparing potential breeding designs with numbers of strains that far exceed (500-1,000) the realized population (50-70). As such, directly comparing these studies is challenging. The closest comparison case is for a 5% QTL with 45% background strain effect with 100 simulated strains with 10 replicates, for which Valdar *et al.* (2006a) estimates 4% power. Matching those settings with the exception of 72 strains instead of 100, and using the DAMB definition of QTL effect size, we find 0.4% power. The relatively lower power with the realized data likely reflects both reduction in the number of strains by 28% (72 to 100) and the deviations from an ideally-randomized population, such as the observed reduction in contributions from the CAST and PWK founders (Srivastava *et al.* 2017). This emphasizes the challenge in projecting the results from Valdar *et al.* (2006a) into the realized population for the purpose of designing an experiment.

We did not attempt power simulations with epistatic QTL or phenotypes with large background strain effect. From the results of Valdar *et al.* (2006a), it was clear that mapping studies in the realized CC, even with replicates, would not be well-powered in those contexts. Nonetheless, despite the reduced number of strains of realized population, we found that successful mapping experiments can be designed in the realized CC, particularly by harnessing the ability of genetic replicates to reduce random noise, as well as within the context of molecular phenotypes such as gene expression for which the genetic architecture is relatively simple.

### Interpreting QTL effect sizes

Our simulations suggest that QTL mapping experiments in the CC are well-powered for large-effect QTL, in the neighborhood of 20-40%, depending on the number of strains and replicates, and the presence of a background strain effect. As such, it is useful to provide some context for what traits might plausibly yield QTL of this size. That said, we note that comparisons of reported estimates of QTL effect size should be interpreted with caution since they vary across different traits and model systems, are calculated under different experimental protocols that may imply different levels of noise, such as different numbers of strains or replicates, and may be estimated by different analysis conditions (statistical methods, data transformations, etc.). And ultimately, these estimates are subject to overestimation due to both the aforementioned Beavis effect and reporting bias.

Multiple studies in the pre-CC, which had more strains than the realized CC population, have reported QTL effect sizes for a variety of traits. Philip *et al.* (2011) report effect sizes for 17 QTL for 102 morphological and behavioral traits in 235 incipient CC strains, ranging from 5.3% (tail-clip latency) to 26% (red cell distribution width). Durrant *et al.* (2011) mapped seven QTL for susceptibility to *Aspergillus fumigatus* infection in 371 mice from 66 strains, with effects ranging from 12.2-16.2%. Gralinski *et al.* (2015) identified four SARS susceptibility QTL in 140 strains with effect sizes between 21-26% (vascular cuffing, 21% and 26%; viral titer, 22%; eosinophilia, 26%).

More closely mirroring the number of strains considered here, Levy *et al.* (2015) detected six strong QTL for traits related to trabecular bone microstructure using 160 mice from 31 strains, which ranged from 61-86%. In an ongoing project involving the mapping of expression QTL (eQTL) from RNA-seq data collected from three tissues of single individuals from 47 strains, 478-739 eQTL were detected at genome-wide significance, ranging in effect size from 60-90%. These results reiterate that QTL mapping studies in the CC are best suited for detection of large effect QTL, as are more common in molecular traits.

In considering the above, it is useful to understand how this relates to effect sizes seen in humans, for which the CC is often used as a model system (Flint and Mackay 2009). In particular, human GWASs, which often use much larger sample sizes, routinely report QTL with estimated effect sizes far smaller than is detectable in the CC. Nonetheless, there are reasons to expect effect sizes in the CC to be larger than in humans. Human GWASs are observational, and as such include many additional sources of noise, reducing QTL effect sizes relative to what would be possible in more tightly-controlled experimental designs. Experimental populations will also have larger QTL effect sizes because: 1) they typically have more balanced allele frequencies; 2) in the case of panels of RILs such as the CC, because they are homozygous across the genome, which increases the contrast in additive allele effects and thus boosts additive QTL effect size; and 3), again for RILs, because they furnish biological replicates, which, as illustrated in Eq 4, can increase effect size by reducing individual error.

### Strains versus replicates

When holding the total number of mice fixed, we found that adding more strains improves power and reduces location error to a greater degree than does adding more replicates. Moreover, this inference was made in the absence of a background strain effect—given that replicates reduce individual-level variance but not strain-level variance, the presence of background effects would reduce the relative value of replicates yet further. These observations are consistent with the results of Valdar *et al.* (2006a) and established theoretical arguments (Soller and Beckmann 1990; Knapp and Bridges 1990).

Nonetheless, for many CC mapping experiments we predict that adding replicates will provide considerable value. First, for all but the most highly polygenic traits, mapping on the means of replicates, a strategy originally termed “replicated progeny” (Cowen 1988) or “progeny testing” (Lander and Botstein 1989), will always provide additional power. Indeed, with a limited number of strains available, and the possibility that all available strains are used, replication may sometimes be the only way power can be further increased (Belknap 1998).

Second, replicates provide not only an insurance policy against phenotyping errors, but also a way to average over batches and similar nuisance parameters (Cowen 1988), thus protecting against the negative consequences of gene by environment interactions while also providing the opportunity for such interactions to be detected [*e.g*., Kafkafi *et al.* (2005, 2018)].

Third, replicates enable deeper phenotypic characterization and in particular measurement of strain-level phenotypes that are necessarily a function of multiple individuals. For example, treatment response phenotypes (*e.g*., response to drug) are ideally defined in terms of counterfactual-like observations of drug-treated and vehicle-treated strain replicates [*e.g*., Festing (2010); Crowley *et al.* (2014)] and recombinant inbred lines such as the CC are uniquely able to combine such definitions with QTL mapping [*e.g*., Mosedale *et al.* (2017) and also, in flies, Kislukhin *et al.* (2013); Najarro *et al.* (2015)]. Similarly, strain-specific phenotypic variance ideally requires replicates (Rönnegård and Valdar 2011; Ayroles *et al.* 2015). We did not consider such elaborations here, but we expect the trade-off between number of strains vs replicates will be more nuanced in such cases.

### Population structure in the CC

Our simulations indicate that deviations from equal relatedness in the realized CC strains have introduced a degree of population structure that potentially increases the risk of false positives if not addressed, albeit to a far lesser extent than has been observed in traditional inbred strain association (Kang *et al.* 2008). In particular, null simulations that assumed correlated strain effects due to genetic relatedness increased FPR for our mapping approach when the strain effect was large relative to individual error, as would be the case for a highly heritable polygenic trait or when using many replicates. This elevated FPR supports the use of QTL mapping approaches that account for the effect of genetic similarity on phenotypes, such as a mixed effect model (Kang *et al.* 2008, 2010; Lippert *et al.* 2011; Zhou and Stephens 2012), especially in the context of marginally significant QTL, which may not remain significant given a higher threshold that controls FPR more appropriately. Software packages that can fit the LMM specifically with CC data include our miQTL package (available on GitHub at https://github.com/gkeele/miqtl) and R/qtl2 (Broman *et al.* 2019).

For the analyses reported here, a mixed effect model approach was not feasible owing to its increased computational burden (and in particular, its incompatibility with the computational shortcut in **Appendix A**). Instead, we simulated independent strain effects and employed a fixed effect mapping procedure due to its computational efficiency, especially when computing permutation-based significance thresholds. Nonetheless, the conclusions drawn in this study should be largely consistent with the use of a mixed effect model that correctly controls for correlated strain effects due to genetic relatedness.

### Allelic series, and use of an eight allele mapping model

We found that the allelic series can strongly affect power through its influence on observed allele frequencies. Specifically, imbalanced bi-allelic QTL have significantly reduced mapping power whereas highly multi-allelic QTL do not because the potential for imbalance is reduced.

Regardless of the true allelic series at a QTL, which is unknown in practice, our statistical procedure assumed an eight allele model. For QTL with fewer functional alleles than founder strains, this assumption could reduce power due to the estimation of redundant allele effect parameters. Indeed, QTL consistent with a bi-allelic series have been more powerfully detected in some MPP studies using SNP association (Baud *et al.* 2013; Keele *et al.* 2018).

Nonetheless, multi-allelic QTL (with more than two alleles) do occur. This has been seen, for example, in cis-regulation of gene expression that largely corresponds to the three subspecies lineages of *Mus musculus*, present in the CC (Crowley *et al.* 2015). Moreover, multi-allelic QTL will not be as powerfully detected through SNP association, as seen, for example, in Aylor *et al.* (2011). SNP (or more generally, variant) association also poses additional challenges, such as how to handle regions of the genome (and variants) that are difficult to genotype, as well as the requirement of extensive quality control filtering to remove markers with low minor allele frequencies. These challenges are implicitly reduced in haplotype analysis.

An ideal statistical procedure would formally model the unknown allelic series and their corresponding uncertainty. Though challenging, the development of alternative mapping strategies that specifically account for the allelic series is clearly an imperative methodological advance that would greatly benefit QTL analyses in MPPs with diverse founder alleles. That said, allelic series-aware approaches would likely be computationally expensive and poorly suited to simulation-based power analyses. Meanwhile, in the absence of more sophisticated approaches, the eight allele model, though potentially redundant, has several advantages over SNP association that suggest it will remain a useful (and maybe the default) tool for CC mapping, namely: it encompasses all possible simpler allelic series, implicitly models local epistasis, and, in reflecting the LD decay around detected QTL, more clearly delineates the limits of mapping resolution.

### Inclusion of extinct CC strains in simulations

Our simulations included genomes from CC strains that are now extinct, and also did not include all the CC strains that are currently available. This discrepancy reflects the inherent challenge of maintaining a stable genetic population resource. RI panels, such as the CC, are an approximation to an ideal: they attempt to provide reproducible genomes that can be observed multiple times as well as across multiple studies; yet, as a biological population, the genomes are mutable, and through time will accumulate mutations and drift, and even potentially go extinct.

Although the inclusion of genomes of extinct strains, or those that have drifted since the strains were genotyped, result in power calculations that do not perfectly correspond to the current CC population, they are preferable to simulated genomes, since they represent genomes that were viable at some point. We view the use of extinct genomes as realistic observations of possible genomes that reflect both the potential that more strains will become extinct or be gained from other breeding sites with time, and thus can be reasonably extended to the realized population, now and into the future.

### Future use and directions

Any analysis of power is subject to the assumptions underlying that analysis. One of the advantages of simulation is the ability to evaluate the impact of many of these assumptions, as well as the consideration of new scenarios by re-running the simulation under different settings, or by elaborating the simulation itself. We have attempted to make re-running the simulations under different settings straightforward for other researchers by developing a software package for this purpose. This package could be used to investigate highly-specialized questions, such as the power for specific combinations of CC strains or assessing how the power to detect QTL varies depending on genomic position. In future work, the simulation code itself could be expanded to investigate additional topics of interest, such as how variance heterogeneity or model misspecification influence power.

### Conclusion

We used a focused simulation approach that incorporates realized CC genomes to provide more accurate estimates of QTL mapping power than were previously possible. As such, the results of our simulations provide tailored power calculations to aide the design of future QTL mapping experiments using the CC. Additionally, we evaluate how the balance of alleles at the QTL can strongly influence power to map QTL in the CC. We make available the R package SPARCC that we developed for running these simulations and analyses. It leverages an efficient model fitting approach in order to explore power in a level of detail that has previously been impractical, it is replicable, and it can be extended to user-specified questions of interest.

## Supporting information

The supplementary tables and figures

Example shell script to coordinate separate calls to sparcc_powersim.R

Provides description of options in SPARCC, and provides simple tutorial for using the package

CC haplotype mosaics formatted for SPARCC

SPARCC R package used for all analyses

Overview of data and Supplement

Example R script to aggregate results from multiple calls to sparcc_powersim.R

Example R script to perform large-scale power analysis

R script to generate figures in manuscript and Supplement

## Acknowledgments

This work was primarily supported by the National Institute of General Medical Sciences under awards R01-GM104125 and R35-GM127000 (to W.V) and the National Institute of Environmental Health Sciences under award R01-ES024965 (to S.N.P.K). Computing resources were generously provided by the University of North Carolina Information Technology Services.

## Author contributions

GRK, WLC, SNPK, and WV wrote the manuscript. GRK and WLC performed the statistical analysis. The authors declare no conflicts of interest.

## Appendix A QR decomposition for fast regression

To maximize power to detect QTL while controlling the FPR, permutations to determine significance thresholds are needed, which is computationally expensive and thus the underlying regression functionality must be highly optimized. We accomplish this through the QR matrix decomposition, which we will describe briefly (Venables and Ripley 2002).

Let **X** = **PA** be the *n* × *m* design matrix included in Eq 3, with *m* = 8. The solution for ***β*** from the least squares normal equations 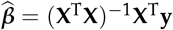. Through the QR decomposition, **X** = **QR**, for which **Q** is an *n* × *p* orthonormal matrix (**Q**^**T**^**Q** = **I**) and **R** is a *m* × *m* upper triangularmatrix. Withmatrix algebra, it is fairly straightforward to show that 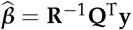, which is also more numerically stable than calculating 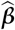 through (**X**^**T**^**X**)^−1^. After solving for 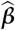, the residual sums of squares, and ultimately logP, can be rapidly calculated. Because our simulation approach involves regressing many permuted outcomes **U**_*p*_**y**^(*s*)^, where **U**_*p*_ is a permutation matrix that re-orders **y**^(*s*)^ randomly, on the same design matrices, computational efficiency can be vastly increased by pre-computing and saving the QR decompositions for all **X**.

Once the QR decomposition has been stored for a design matrix **X**_*j*_, *j* indexing locus, it is highly computationally efficient to conduct additional tests for any **y**, thus encompassing all permuted outcomes **U**_*p*_**y**. If **X**_*j*_ is the same across *S* simulations, the boost in computation can extend beyond permutations to samples of **y**^(*s*)^, as is the case when the set of CC strains is fixed. In effect, two cases result for our R package SPARCC: when the set of CC strains is fixed, and when the set varies.

- Fixed set of CC strains

1. Store QR decompositions of **X**_*j*_ for *j* = 1, 2,…, *J*
2. Run genome scans for **y**^(*s*)^ and **U**_*p*_**y**^(*s*)^ for *s* = 1,2,…, *S* × *p* = 1,2,…, *P*
- Varied set of CC strains

1. Store QR decompositions of **X**_*js*_ for *j* = 1,2,…, *J*
2. Run genome scans for **y**^(*s*)^ and **U**_*p*_**y**^(*s*)^ for *p* = 1,2,…, *P*
3. Repeat steps 1 and 2 for *s* = 1,2,…, *S*

Varying the sets of CC strains increases computation time linearly with respect to *S*. If the investigators do not have a predefined set of strains, it is appropriate that this source of variability be incorporated into the power calculation.

## Appendix B Computing environment and performance

We performed 1,000 simulations (in batches of 100) for each combination of the parameters, resulting in 8,400 individual jobs. These jobs were submitted in parallel to a distributed computing cluster (http://its.unc.edu/rc-services/killdevil-cluster/). Runtime varied depending on parameter settings and the hardware used, with the longest jobs taking approximately seven hours to complete.

## Appendix C CC strains

This study used haplotype mosiac data available from http://csbio.unc.edu/CCstatus/index.py?run=FounderProbs for the following 72 CC strains: CC001, CC002, CC003, CC004, CC005, CC006, CC007, CC008, CC009, CC010, CC011, CC012, CC013, CC014, CC015, CC016, CC017, CC018, CC019, CC020, CC021, CC022, CC023, CC024, CC025, CC026, CC027, CC028, CC029, CC030, CC031, CC032, CC033, CC034, CC035, CC036, CC037, CC038, CC039, CC040, CC041, CC042, CC043, CC044, CC045, CC046, CC047, CC048, CC049, CC050, CC051, CC052, CC053, CC054, CC055, CC056, CC057, CC058, CC059, CC060, CC061, CC062, CC063, CC065, CC068, CC070, CC071, CC072, CC073, CC074, CC075, CC076. This includes two strains CC051 and CC059 that are derived from the same breeding funnel and thus more closely related than typical pairs of CC strains.

Of the the 72 CC strains used here, 54 are among a larger set of 59 that are currently maintained and distributed by UNC (personal correspondence with Darla Miller, UNC). These 54/59 strains are CC001, CC002, CC003, CC004, CC005, CC006, CC007, CC008, CC009, CC010, CC011, CC012, CC013, CC015, CC016, CC017, CC019, CC021, CC023, CC024, CC025, CC026, CC027, CC029, CC030, CC031, CC032, CC033, CC035, CC036, CC037, CC038, CC039, CC040, CC041, CC042, CC043, CC044, CC045, CC046, CC049, CC051, CC053, CC055, CC057, CC058, CC059, CC060, CC061, CC062, CC065, CC068, CC071, CC072. The remaining 5/59 strains (CC078, CC079, CC080, CC081, CC083) lacked haplotype mosaic data at the time of simulation and so were not included (although note that their mosaics have since been added to the website).

## Appendix D Additive model and allelic series matrices

## Additive matrix

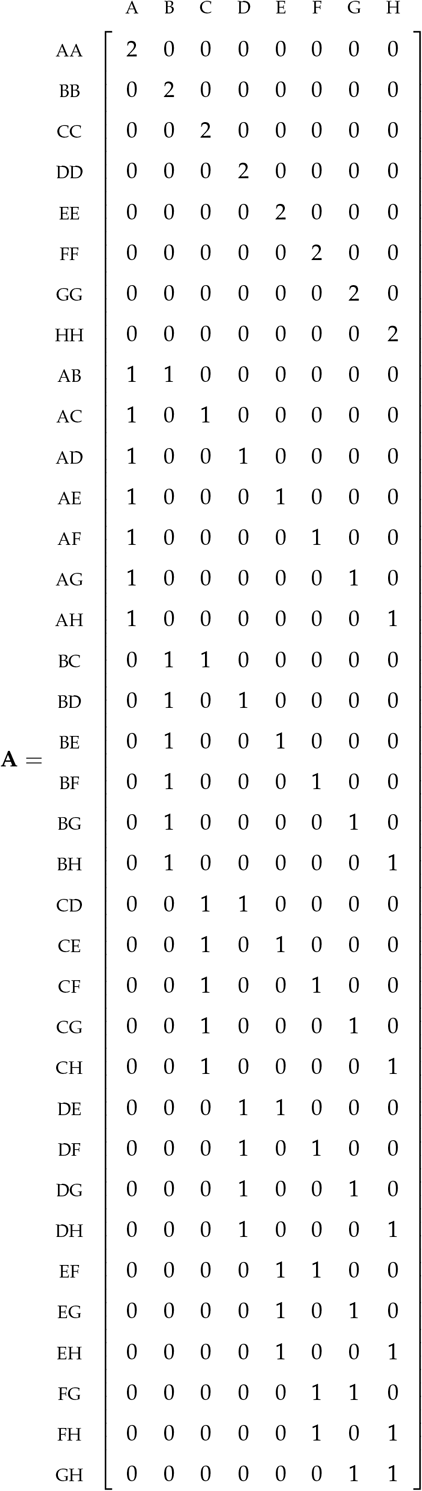

We can use matrices to specify simplifying linear combinations of the 36 diplotypes. The additive model matrix **A** is commonly used, and we use it here. Post-multiplication of the diplotype design matrix **D** with the **A** rotates the diplotypes at the locus to dosages of the founder haplotypes. If there is no uncertainty on the diplotype identities, **DA** will be the matrix of founder haplotype counts at the locus.

## Allelic series matrices

We explore the influence of the allelic series on QTL mapping power through the simulation procedure. The QTL mapping procedure estimates separate parameters for each founder, though in reality, there are likely fewer functional alleles. We denote the *q*^th^ functional allele as *k*_*q*_. The allelic series can be sampled and encoded in the `M.ID` argument within the `sim.CC.data()` function of SPARCC. Below are examples of balanced (4v4) and unbalanced (7v1) bi-allelic series, as well as tri-allelic series.

## Allelic series with eight alleles (maximum)

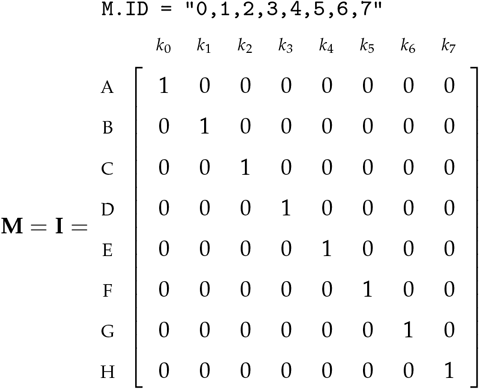

## Example balanced (4v4) bi-allelic series

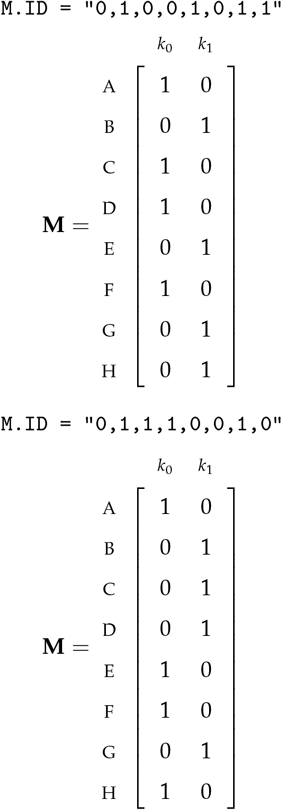

## Example unbalanced (7v1) bi-allelic series

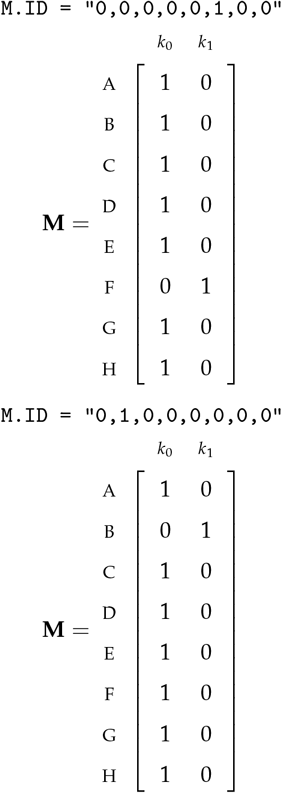

## Example tri-allelic series

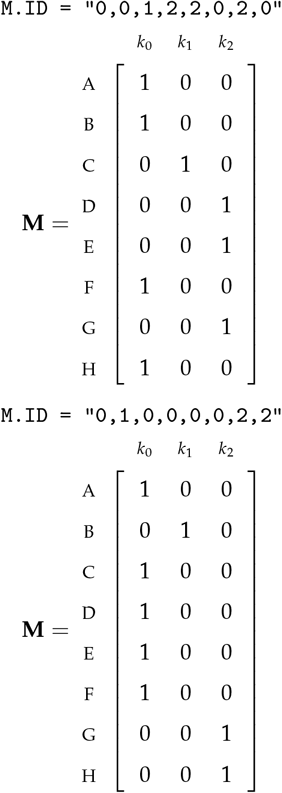

## Literature Cited

Aylor, D. L., W. Valdar, W. Foulds-mathes, R. J. Buus, R. A. Verdugo, et al., 2011 Genetic analysis of complex traits in the emerging Collaborative Cross. Genome research 21: 1213–22.

Ayroles, J. F., S. M. Buchanan, C. O’Leary, K. Skutt-Kakaria, J. K. Grenier, et al., 2015 Behavioral idiosyncrasy reveals genetic control of phenotypic variability. Proceedings of the National Academy of Sciences 112: 6706–6711.

Baud, A., R. Hermsen, V. Guryev, P. Stridh, D. Graham, et al., 2013 Combined sequence-based and genetic mapping analysis of complex traits in outbred rats. Nature genetics 45: 767–75.

Beavis, W., 1994 The power and deceit of qtl experiments: lessons from comparative qtl studies. In Proceedings of the forty-ninth annual corn and sorghum industry research conference, pp. 250–266, Washington, DC.

Belknap, J. K., 1998 Effect of within-strain sample size on GTL detection and mapping using recombinant inbred mouse strains. Behavior Genetics 28: 29–38.

Belknap, J. K., S. R. Mitchell, L. A. O’Toole, M. L. Helms, and J. C. Crabbe, 1996 Type I and type II error rates for quantitative trait loci (QTL) mapping studies using recombinant inbred mouse strains. Behavior genetics 26: 149–60.

Bouchet, S., M. O. Olatoye, S. R. Marla, R. Perumal, T. Tesso, et al., 2017 Increased Power To Dissect Adaptive Traits in Global Sorghum Diversity Using a Nested Association Mapping Population. Genetics 206: 573–585.

Broman, K. W., D. M. Gatti, P. Simecek, N. A. Furlotte, P. Prins, et al., 2019 R/qtl2: Software for Mapping Quantitative Trait Loci with High-Dimensional Data and Multiparent Populations. Genetics 211: 495–502.

Brown, L. D., T. T. Cai, and A. DasGupta, 2001 Interval Estimation for a Binomial Proportion. Statistical Science 16: 101–117.

Chesler, E. J., D. R. Miller, L. R. Branstetter, L. D. Galloway, B. L. Jackson, et al., 2008 The Collaborative Cross at Oak Ridge National Laboratory: developing a powerful resource for systems genetics. Mammalian Genome 19: 382–389.

Churchill, G. A., D. C. Airey, H. Allayee, J. M. Angel, A. D. Attie, et al., 2004 The Collaborative Cross, a community resource for the genetic analysis of complex traits. Nature Genetics 36: 1133–1137.

Collaborative Cross Consortium, 2012 The genome architecture of the Collaborative Cross mouse genetic reference population. Genetics 190: 389–401.

Cowen, N. M., 1988 The use of replicated progenies in marker-based mapping of QTL’s. Theoretical and Applied Genetics 75: 857–862.

Crowley, J. J., Y. Kim, A. B. Lenarcic, C. R. Quackenbush, C. J. Barrick, et al., 2014 Genetics of adverse reactions to haloperidol in a mouse diallel: a drug-placebo experiment and Bayesian causal analysis. Genetics 196: 321–47.

Crowley, J. J., V. Zhabotynsky, W. Sun, S. Huang, I. K. Pakatci, et al., 2015 Analyses of allele-specific gene expression in highly divergent mouse crosses identifies pervasive allelic imbalance. Nature genetics 47: 353–60.

Dell’Acqua, M., D. M. Gatti, G. Pea, F. Cattonaro, F. Coppens, et al., 2015 Genetic properties of the MAGIC maize population: a new platform for high definition QTL mapping in Zea mays. Genome biology 16: 167.

Doerge, R. and G. Churchill, 1996 Permutation tests for multiple loci affecting a quantitative character. Genetics 142: 285–94.

Donoghue, L. J., A. Livraghi-Butrico, K. M. McFadden, J. M. Thomas, G. Chen, et al., 2017 Identification of trans Protein QTL for Secreted Airway Mucins in Mice and a Causal Role for Bpifb1. Genetics 207: 801–812.

Dudbridge, F. and B. P. Koeleman, 2004 Efficient Computation of Significance Levels for Multiple Associations in Large Studies of Correlated Data, Including Genomewide Association Studies. The American Journal of Human Genetics 75: 424–435.

Durrant, C., H. Tayem, B. Yalcin, J. Cleak, L. Goodstadt, et al., 2011 Collaborative Cross mice and their power to map host susceptibility to Aspergillus fumigatus infection. Genome research 21: 1239–48.

Falke, K. C. and M. Frisch, 2011 Power and false-positive rate in QTL detection with near-isogenic line libraries. Heredity 106: 576–584.

Ferris, M. T., D. L. Aylor, D. Bottomly, A. C. Whitmore, L. D. Aicher, et al., 2013 Modeling Host Genetic Regulation of Influenza Pathogenesis in the Collaborative Cross. PLoS Pathogens 9: e1003196.

Festing, M. F. W., 2010 Inbred strains should replace outbred stocks in toxicology, safety testing, and drug development. Toxicologic Pathology 38: 681–690.

Flint, J. and T. F. Mackay, 2009 Genetic architecture of quantitative traits in mice, flies, and humans. Genome Research 19: 723–733.

Fu, C.-P., C. E. Welsh, F. P.-M. de Villena, and L. McMillan, 2012 Inferring ancestry in admixed populations using microarray probe intensities. In Proceedings of the ACM Conference on Bioinformatics, Computational Biology and Biomedicine -BCB ’12, pp. 105–112, New York, New York, USA, ACM Press.

Gatti, D. M., K. L. Svenson, A. Shabalin, L.-Y. Wu, W. Valdar, et al., 2014 Quantitative Trait Locus Mapping Methods for Diversity Outbred Mice. G3 (Bethesda, Md.) 4: 1623–1633.

Graham, J. B., J. L. Swarts, M. Mooney, G. Choonoo, S. Jeng, et al., 2017 Extensive Homeostatic T Cell Phenotypic Variation within the Collaborative Cross. Cell reports 21: 2313–2325.

Gralinski, L. E., M. T. Ferris, D. L. Aylor, A. C. Whitmore, R. Green, et al., 2015 Genome Wide Identification of SARS-CoV Susceptibility Loci Using the Collaborative Cross. PLoS genetics 11: e1005504.

Haley, C. S. and S. A. Knott, 1992 A simple regression method for mapping quantitative trait loci in line crosses using flankingmarkers. Heredity 69: 315–24.

Kaeppler, S. M., 1997 Quantitative trait locus mapping using sets of near-isogenic lines: Relative power comparisons and technical considerations. Theoretical and Applied Genetics 95: 384–392.

Kafkafi, N., J. Agassi, E. J. Chesler, J. C. Crabbe, W. E. Crusio, et al., 2018 Reproducibility and replicability of rodent phenotyping in preclinical studies. Neuroscience and Biobehavioral Reviews 87: 218–232.

Kafkafi, N., Y. Benjamini, A. Sakov, G. I. Elmer, and I. Golani, 2005 Genotype-environment interactions in mouse behavior: a way out of the problem. Proceedings of the National Academy of Sciences of the United States of America 102: 4619–24.

Kang, H. M., J. H. Sul, S. K. Service, N. A. Zaitlen, S.-Y. Kong, et al., 2010 Variance component model to account for sample structure in genome-wide association studies. Nature genetics 42: 348–354.

Kang, H. M., N. A. Zaitlen, C. M. Wade, A. Kirby, D. Heckerman, et al., 2008 Efficient control of population structure in model organism association mapping. Genetics 178: 1709–23.

Keele, G. R., J. W. Prokop, H. He, K. Holl, J. Littrell, et al., 2018 Genetic Fine-Mapping and Identification of Candidate Genes and Variants for Adiposity Traits in Outbred Rats. Obesity 26: 213–222.

Kelada, S. N. P., 2016 Plethysmography Phenotype QTL in Mice Before and After Allergen Sensitization and Challenge. G3 (Bethesda, Md.) 6: 2857–2865.

Kelada, S. N. P., D. L. Aylor, B. C. E. Peck, J. F. Ryan, U. Tavarez, et al., 2012 Genetic Analysis of Hematological Parameters in Incipient Lines of the Collaborative Cross. G3 (Bethesda, Md.) 2: 157–165.

King, E. G. and A. D. Long, 2017 The Beavis Effect in Next-Generation Mapping Panels in Drosophila melanogaster. G3 7: 1643 LP–1652.

King, E. G., S. J. Macdonald, and A. D. Long, 2012 Properties and power of the Drosophila synthetic population resource for the routine dissection of complex traits. Genetics 191: 935–949.

Kislukhin, G., E. G. King, K. N. Walters, S. J. Macdonald, and A. D. Long, 2013 The Genetic Architecture of Methotrexate Toxicity Is Similar in Drosophila melanogaster and Humans. G3: Genes, Genomes, Genetics 3: 1301–1310.

Klasen, J. R., H. P. Piepho, and B. Stich, 2012 QTL detection power of multi-parental RIL populations in Arabidopsis thaliana. Heredity 108: 626–632.

Knapp, S. J. and W. C. Bridges, 1990 Using molecular markers to estimate quantitative trait locus parameters: power and genetic variances for unreplicated and replicated progeny. Genetics 126: 769–77.

Kover, P. X., W. Valdar, J. Trakalo, N. Scarcelli, I. M. Ehrenreich, et al., 2009 A Multiparent Advanced Generation Inter-Cross to fine-map quantitative traits in Arabidopsis thaliana. PLoS genetics 5: e1000551.

Lander, E. S. and D. Botstein, 1989 Mapping mendelian factors underlying quantitative traits using RFLP linkage maps. Genetics 121: 185–99.

Levy, R., R. F. Mott, F. A. Iraqi, and Y. Gabet, 2015 Collaborative cross mice in a genetic association study reveal new candidate genes for bone microarchitecture. BMC Genomics 16: 1013.

Li, H., P. Bradbury, E. Ersoz, E. S. Buckler, and J. Wang, 2011 Joint QTL linkage mapping for multiple-cross mating design sharing one common parent. PloS one 6: e17573.

Lippert, C., J. Listgarten, Y. Liu, C. M. Kadie, R. I. Davidson, et al., 2011 FaST linear mixed models for genome-wide association studies. Nature Methods 8: 833–837.

Liu, E. Y., Q. Zhang, L. McMillan, F. P.-M. de Villena, and W. Wang, 2010 Efficient genome ancestry inference in complex pedigrees with inbreeding. Bioinformatics 26: i199–i207.

Lorè, N. I., F. A. Iraqi, and A. Bragonzi, 2015 Host genetic diversity influences the severity of Pseudomonas aeruginosa pneumonia in the Collaborative Cross mice. BMC genetics 16: 106.

Lynch, M. and B. Walsh, 1998 Genetics and Analysis of Quantitative Traits. Sinauer Associates, Sunderland, MA.

Mackay, T. F. C., S. Richards, E. A. Stone, A. Barbadilla, J. F. Ayroles, et al., 2012 The Drosophila melanogaster Genetic Reference Panel. Nature 482: 173–8.

Martínez, O. and R. N. Curnow, 1992 Estimating the locations and the sizes of the effects of quantitative trait loci using flanking markers. Theor. Appl. Genet. 85: 480–488.

Mathes, W. F., D. L. Aylor, D. R. Miller, G. A. Churchill, E. J. Chesler, et al., 2011 Architecture of energy balance traits in emerging lines of the Collaborative Cross. American Journal of Physiology-Endocrinology and Metabolism 300: E1124–E1134.

Molenhuis, R. T., H. Bruining, M. J. V. Brandt, P. E. van Soldt, H. J. Abu-Toamih Atamni, et al., 2018 Modeling the quantitative nature of neurodevelopmental disorders using Collaborative Cross mice. Molecular Autism 9: 63.

Morgan, A. P., C.-P. Fu, C.-Y. Kao, C. E. Welsh, J. P. Didion, et al., 2016 The Mouse Universal Genotyping Array: From Substrains to Subspecies. G3: Genes, Genomes, Genetics 6: 263–279.

Mosedale, M., Y. Kim, W. J. Brock, S. E. Roth, T. Wiltshire, et al., 2017 Candidate Risk Factors and Mechanisms for Tolvaptan-Induced Liver Injury Are Identified Using a Collaborative Cross Approach. Toxicological Sciences 156: kfw269.

Mott, R., C. J. Talbot, M. G. Turri, A. C. Collins, and J. Flint, 2000 A method for fine mapping quantitative trait loci in outbred animal stocks. PNAS 97: 12649–54.

Najarro, M. A., J. L. Hackett, B. R. Smith, C. A. Highfill, E. G. King, et al., 2015 Identifying Loci Contributing to Natural Variation in Xenobiotic Resistance in Drosophila. PLoS Genetics 11: 1–25.

Noble, L. M., I. Chelo, T. Guzella, B. Afonso, D. D. Riccardi, et al., 2017 Polygenicity and Epistasis Underlie Fitness-Proximal Traits in the Caenorhabditis elegans Multiparental Experimental Evolution (CeMEE) Panel. Genetics 207: genetics.300406.2017.

Orgel, K., J. M. Smeekens, P. Ye, L. Fotsch, R. Guo, et al., 2019 Genetic diversity between mouse strains allows identification of the CC027/GeniUnc strain as an orally reactive model of peanut allergy. The Journal of allergy and clinical immunology 143: 1027–1037.e7.

Peirce, J. L., L. Lu, J. Gu, L. M. Silver, and R. W. Williams, 2004 A new set of BXD recombinant inbred lines from advanced intercross populations in mice. BMC genetics 5: 7.

Pfaff, B. and A. McNeil, 2018 evir: Extreme Values in R. R package version 1.7-4.

Philip, V. M., G. Sokoloff, C. L. Ackert-Bicknell, M. Striz, L. Branstetter, et al., 2011 Genetic analysis in the Collaborative Cross breeding population. Genome Research 21: 1223–1238.

Phillippi, J., Y. Xie, D. R. Miller, T. A. Bell, Z. Zhang, et al., 2014 Using the emerging Collaborative Cross to probe the immune system. Genes & Immunity 15: 38–46.

R Core Team, 2018 R: A Language and Environment for Statistical Computing. R Foundation for Statistical Computing, Vienna, Austria.

Ram, R., M. Mehta, L. Balmer, D. M. Gatti, and G. Morahan, 2014 Rapid identification of major-effect genes using the collaborative cross. Genetics 198: 75–86.

Rasmussen, A. L., A. Okumura, M. T. Ferris, R. Green, F. Feldmann, et al., 2014 Host genetic diversity enables Ebola hemorrhagic fever pathogenesis and resistance. Science (New York, N.Y.) 346: 987–91.

Rogala, A. R., A. P. Morgan, A. M. Christensen, T. J. Gooch, T. A. Bell, et al., 2014 The Collaborative Cross as a resource for modeling human disease: CC011/Unc, a new mouse model for spontaneous colitis. Mammalian genome 25: 95–108.

Rönnegård, L. and W. Valdar, 2011 Detecting major genetic loci controlling phenotypic variability in experimental crosses. Genetics 188: 435–447.

Rutledge, H., D. L. Aylor, D. E. Carpenter, B. C. Peck, P. Chines, et al., 2014 Genetic regulation of Zfp30, CXCL1, and neutrophilic inflammation in murine lung. Genetics 198: 735–745.

Shorter, J. R., F. Odet, D. L. Aylor, W. Pan, C.-Y. Kao, et al., 2017 Male Infertility Is Responsible for Nearly Half of the Extinction Observed in the Mouse Collaborative Cross. Genetics 206: 557–572.

Shusterman, A., Y. Salyma, A. Nashef, M. Soller, A. Wilensky, et al., 2013 Genotype is an important determinant factor of host susceptibility to periodontitis in the Collaborative Cross and inbred mouse populations. BMC genetics 14: 68.

Soller, M. and J. S. Beckmann, 1990 Marker-based mapping of quantitative trait loci using replicated progenies. Theoretical and Applied Genetics 80: 205–208.

Srivastava, A., A. P. Morgan, M. L. Najarian, V. K. Sarsani, J. S. Sigmon, et al., 2017 Genomes of the mouse Collaborative Cross. Genetics 206: 537–556.

Svenson, K. L., D. M. Gatti, W. Valdar, C. E. Welsh, R. Cheng, et al., 2012 High-resolution genetic mapping using the Mouse Diversity outbred population. Genetics 190: 437–47.

Takuno, S., R. Terauchi, and H. Innan, 2012 The power of QTL mapping with RILs. PloS one 7: e46545.

Threadgill, D. W. and G. A. Churchill, 2012 Ten Years of the Collaborative Cross. Genetics 190: 291–294.

Threadgill, D. W., K. W. Hunter, and R. W. Williams, 2002 Genetic dissection of complex and quantitative traits: from fantasy to reality via a community effort. Mammalian genome: official journal of the International Mammalian Genome Society 13: 175–8.

Valdar, W., J. Flint, and R. Mott, 2006a Simulating the Collaborative Cross: power of quantitative trait loci detection and mapping resolution in large sets of recombinant inbred strains of mice. Genetics 172: 1783–97.

Valdar, W., L. C. Solberg, D. Gauguier, S. Burnett, P. Klenerman, et al., 2006b Genome-wide genetic association of complex traits in heterogeneous stock mice. Nature Genetics 38: 879–887.

Venables, W. N. and B. D. Ripley, 2002 Modern Applied Statistics with S. Springer, New York, fourth edition, ISBN 0-387-954570.

Venkatratnam, A., S. Furuya, O. Kosyk, A. Gold, W. Bodnar, et al., 2017 Collaborative Cross Mouse Population Enables Refinements to Characterization of the Variability in Toxicokinetics of Trichloroethylene and Provides Genetic Evidence for the Role of PPAR Pathway in Its Oxidative Metabolism. Toxicological Sciences 158: 48–62.

Vered, K., C. Durrant, R. Mott, and F. A. Iraqi, 2014 Susceptibility to Klebsiella pneumonaie infection in collaborative cross mice is a complex trait controlled by at least three loci acting at different time points. BMC genomics 15: 865.

Welsh, C. E., D. R. Miller, K. F. Manly, J. Wang, L. McMillan, et al., 2012 Status and access to the Collaborative Cross population. Mammalian genome: official journal of the International Mammalian Genome Society 23: 706–12.

Xu, S., 2003 Theoretical basis of the Beavis effect. Genetics 165: 2259–68.

Yalcin, B., J. Flint, and R. Mott, 2005 Using progenitor strain information to identify quantitative trait nucleotides in outbred mice. Genetics 171: 673–81.

Yamamoto, E., H. Iwata, T. Tanabata, R. Mizobuchi, J.-i. Yonemaru, et al., 2014 Effect of advanced intercrossing on genome structure and on the power to detect linked quantitative trait loci in a multi-parent population: a simulation study in rice. BMC genetics 15: 50.

Yang, H., J. R. Wang, J. P. Didion, R. J. Buus, T. A. Bell, et al., 2011 Subspecific origin and haplotype diversity in the laboratory mouse. Nature genetics 43: 648–55.

Yu, J., J. B. Holland, M. D. McMullen, and E. S. Buckler, 2008 Genetic design and statistical power of nested association mapping in maize. Genetics 178: 539–551.

Zhang, Z., W. Wang, and W. Valdar, 2014 Bayesian modeling of haplotype effects in multiparent populations. Genetics 198: 139–56.

Zheng, C., M. P. Boer, and F. A. van Eeuwijk, 2015 Reconstruction of Genome Ancestry Blocks in Multiparental Populations. Genetics 200: 1073–1087.

Zhou, X. and M. Stephens, 2012 Genome-wide efficient mixed-model analysis for association studies. Nature genetics 44: 821–4.

Zollner, S. and J. K. Pritchard, 2007 Overcoming the winner’s curse: estimating penetrance parameters from case-control data. American journal of human genetics 80: 605–15.

Zou, F., Z. Xu, and T. Vision, 2006 Assessing the significance of quantitative trait loci in replicable mapping populations. Genetics 174: 1063–8.

