## Supplementary material for "Determinants of QTL mapping power in the realized Collaborative Cross": The supplementary tables and figures

### Supplemental Tables and Figures

**Table S1 QTL mapping power in the Collaborative Cross based on QTL effect sizes in the mapping population (Definition DAMB)**

| QTL | | | Power | | | | | | | | |
|---|---|---|---|---|---|---|---|---|---|---|---|
| | | | 30 strains | | | 50 strains | | | 72 strains | | |
| 1 obs[a] | 3 rep[b] | 5 rep[b] | 2 alleles | 3 alleles | 8 alleles | 2 alleles | 3 alleles | 8 alleles | 2 alleles | 3 alleles | 8 alleles |
| 0.01 | 0.003 | 0.002 | 0.000 | 0.002 | 0.001 | 0.000 | 0.000 | 0.000 | 0.000 | 0.003 | 0.000 |
| 0.05 | 0.017 | 0.010 | 0.000 | 0.001 | 0.001 | 0.000 | 0.002 | 0.000 | 0.005 | 0.003 | 0.004 |
| 0.1 | 0.036 | 0.022 | 0.004 | 0.002 | 0.000 | 0.009 | 0.007 | 0.003 | 0.015 | 0.016 | 0.018 |
| 0.15 | 0.056 | 0.034 | 0.002 | 0.006 | 0.001 | 0.016 | 0.017 | 0.018 | 0.056 | 0.051 | 0.046 |
| 0.2 | 0.077 | 0.048 | 0.008 | 0.007 | 0.003 | 0.032 | 0.038 | 0.032 | 0.119 | 0.141 | 0.135 |
| 0.25 | 0.100 | 0.062 | 0.009 | 0.005 | 0.008 | 0.071 | 0.066 | 0.088 | 0.264 | 0.264 | 0.281 |
| 0.3 | 0.125 | 0.079 | 0.015 | 0.013 | 0.014 | 0.141 | 0.120 | 0.134 | 0.460 | 0.466 | 0.492 |
| 0.35 | 0.152 | 0.097 | 0.028 | 0.029 | 0.030 | 0.234 | 0.229 | 0.262 | 0.695 | 0.664 | 0.684 |
| 0.4 | 0.182 | 0.118 | 0.045 | 0.038 | 0.040 | 0.415 | 0.376 | 0.413 | 0.854 | 0.848 | 0.854 |
| 0.45 | 0.214 | 0.141 | 0.082 | 0.074 | 0.078 | 0.603 | 0.594 | 0.620 | 0.958 | 0.964 | 0.974 |
| 0.5 | 0.250 | 0.167 | 0.136 | 0.134 | 0.143 | 0.769 | 0.783 | 0.783 | 0.996 | 0.998 | 0.999 |
| 0.55 | 0.289 | 0.196 | 0.198 | 0.204 | 0.248 | 0.911 | 0.922 | 0.924 | 1.000 | 0.999 | 1.000 |
| 0.6 | 0.333 | 0.231 | 0.334 | 0.331 | 0.328 | 0.985 | 0.980 | 0.994 | 0.999 | 1.000 | 1.000 |
| 0.65 | 0.382 | 0.271 | 0.519 | 0.489 | 0.534 | 0.998 | 0.995 | 0.999 | 0.999 | 1.000 | 1.000 |
| 0.7 | 0.438 | 0.318 | 0.707 | 0.703 | 0.756 | 0.998 | 0.999 | 1.000 | 0.999 | 0.999 | 1.000 |
| 0.75 | 0.500 | 0.375 | 0.866 | 0.864 | 0.914 | 0.998 | 0.998 | 1.000 | 0.999 | 1.000 | 1.000 |
| 0.8 | 0.571 | 0.444 | 0.940 | 0.954 | 0.979 | 0.996 | 0.998 | 1.000 | 1.000 | 1.000 | 1.000 |
| 0.85 | 0.654 | 0.531 | 0.962 | 0.967 | 0.995 | 0.998 | 0.999 | 1.000 | 1.000 | 0.999 | 1.000 |
| 0.9 | 0.750 | 0.643 | 0.978 | 0.981 | 0.998 | 0.999 | 0.998 | 1.000 | 1.000 | 1.000 | 1.000 |
| 0.95 | 0.864 | 0.792 | 0.970 | 0.988 | 0.999 | 0.999 | 0.999 | 1.000 | 0.999 | 1.000 | 1.000 |

[a] Convert QTL effect sizes from experiments with replicates to mean scale with Eq 4.
[b] Based on no background strain effect.

**Table S2 False positive rate in the Collaborative Cross with no simulated QTL and the presence of population structure (also in Figure 6)**

| | False positive rate | |
|---|---|---|
| Background Strain[a] | 71[b] strains | 72 strains |
| 0 | 0.058 | 0.055 |
| 0.2 | 0.0648 | 0.062 |
| 0.4 | 0.078 | 0.083 |
| 0.6 | 0.092 | 0.102 |
| 0.8 | 0.109 | 0.124 |
| 1 | 0.129 | 0.145 |

[a] Correlated based on the realized genomic similarity of the strains.
[b] Excluding CC059, cousin strain of CC051.

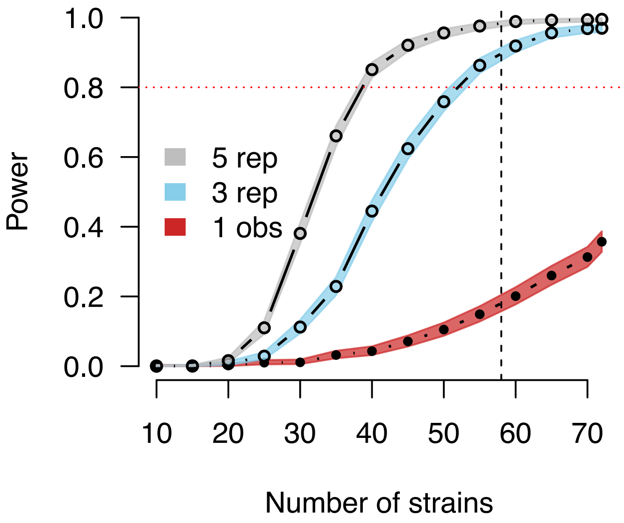


**Figure S1** Power estimates for experiments with three and five replicates interpolated from estimates from only a single observation per CC strain. Power curves correspond to a QTL with effect size of 30% and two functional alleles. QTL effect sizes for experiments with replicates are adjusted based on Eq 4, allowing for results from single observation simulations to be projected into experiments with replicates. Pre-computed power estimates for single observation simulations are stored in SPARCC and can conveniently be extrapolated into other settings, as is demonstrated here. The horizontal red dotted line marks 80% power. The vertical black dashed line marks 58 strains, which is currently the number of unrelated strains available from UNC. Closed circles represent power estimates that were directly evaluated. Open circles represent power estimates that were interpolated from single observation results.

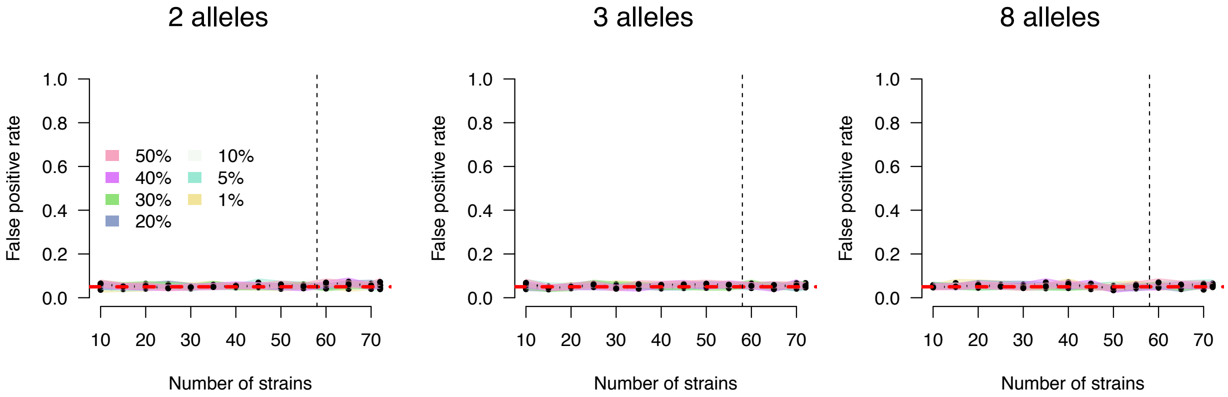


**Figure S2** False positive rate (FPR) based on 1,000 simulations per setting with respect to number of CC strains, stratified by the number of functional alleles. The horizontal red dashed line marks the 5% type I error (false positive) rate. CC strains and loci were varied in simulations, resulting in false positive rates that average over loci and strain combinations. Confidence intervals were calculated based on Jeffreys interval (Brown *et al.* 2001) for a binomial proportion. Plots, left to right, correspond to two, three, and eight functional alleles. The FPR represents the probability that any QTL is detected on chromosomes other than the chromosome on which the simulated QTL is located. The significance thresholds maintain the desired type I error rate of 0.05. As expected, the allelic series does not appear to influence FPR. The vertical black dashed line marks 58 strains, which is currently the number of unrelated strains available from UNC.

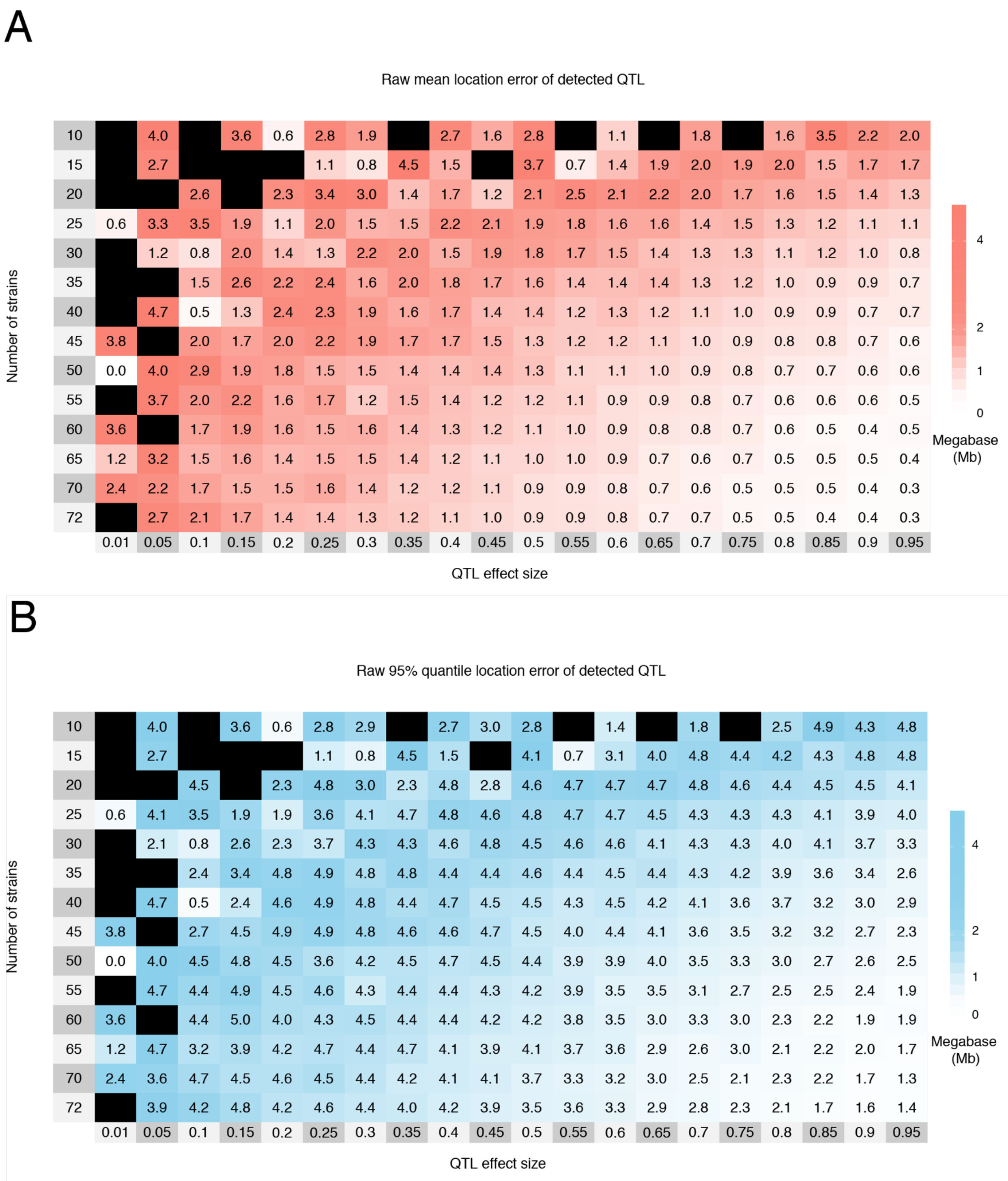


**Figure S3** The raw mean (A) and 95% quantile (B) of the location error, the distance in Mb between the detected and simulated QTL, by effect size and number of strains for 1,000 simulations of each setting. These simulations are based on Definition B with an eight allele QTL, and only a single observation per strain. Cells are colored red to white with decreasing mean and blue to white with decreasing 95% quantile. Black cells represent the case in which no simulated QTL were detected. Estimates from poorly-powered settings are more likely to be unobserved or unstable from low detection. Regularized measurements are provided in **Figure 4**. Increasing the number of strains reduces both the mean and 95% quantile location error more so than QTL effect size, also shown in **Figure S6**. The maximum possible location error was 5Mb due to the 10Mb window centered around the true QTL position used for detecting QTL.

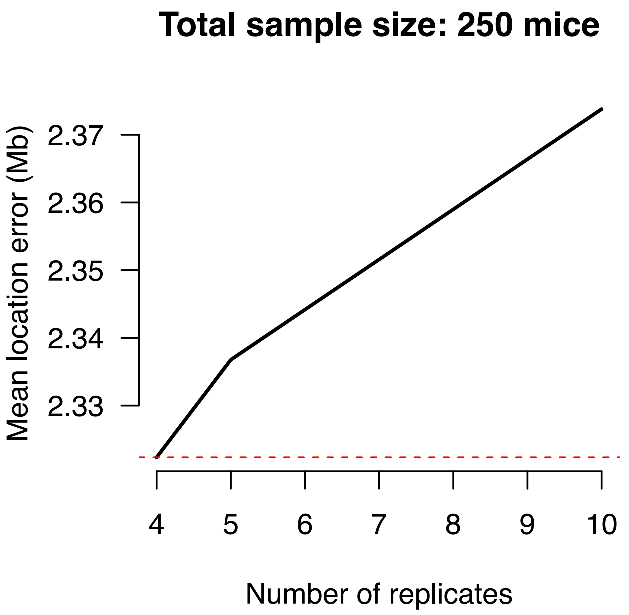


**Figure S4** Mean location error of detected QTL increases with the number of replicates while keeping total sample size fixed. Estimates are based on linear interpolation from dense simulations using Definition B with single observations per strains. The total number of mice and the QTL effect size are fixed at 250 and 50%, respectively. The red dotted line highlights that the lowest mean location error occurs at 4, the lowest number of replicates possible for a sample of 250 mice, given the 72 strains used in the simulations.

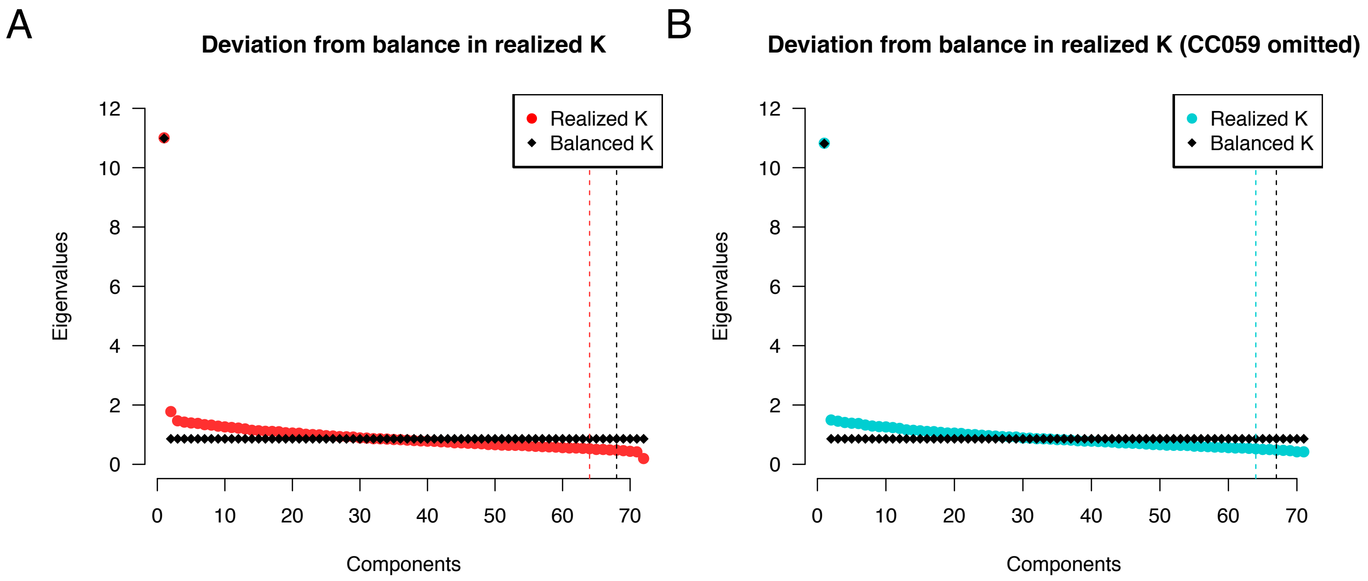


**Figure S5** The realized genetic relationship matrix $\mathbf{K}$ deviates from a perfectly balanced population. Red and blue circles represent the eigenvalues of the eigendecomposition of the realized $\mathbf{K}$, when including both cousin strains (A) and excluding one (B). Black diamonds represent the eigenvalues of a balanced $\mathbf{K}$, with the relationship fixed at the mean relationship observed in the realized $\mathbf{K}$. Vertical dashed lines represent the number of components necessary to explain 95% of the variation for the different $\mathbf{K}$. The first eigenvalue represents the variation accounted for by the overall mean of $\mathbf{K}$. In the balanced $\mathbf{K}$, after removing the effect of the mean, all components contribute equally to the variance. The eigenvalue of the second component for the 72 strains is slightly inflated, representing the cousin strains, a notable deviation from equal relatedness. This inflation disappears when one of the cousin strains is removed, however population structure still persists.

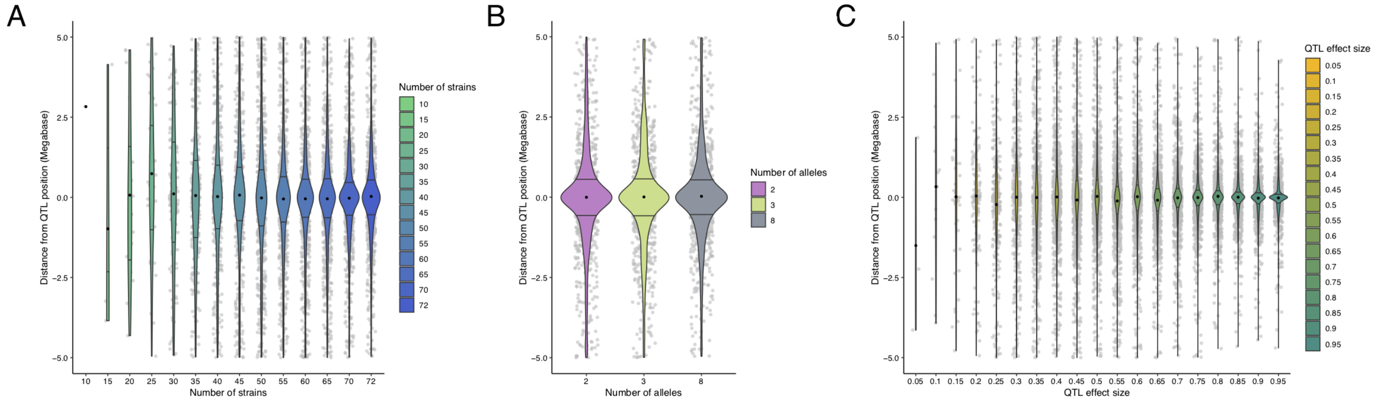


**Figure S6** Distributions of the un-regularized location error, by number of strains (A), number of alleles (B), and QTL effect size (C). Observed distances are between -5 and 5Mb due to the 10Mb window centered around the simulated QTL that was used for QTL detection in the large scale results. Gray dots represent the distances for a single simulations. The colored violin plots represent the distribution of distances across the simulations. The black dot marks the mean location error for each category. Horizontal lines represent the 25$^{th}$ and 75$^{th}$ quantiles. (A) With QTL effect size fixed at 50% and the number of alleles at 8, as the number of CC strains increases, the distribution of location error becomes more concentrated around zero, meaning the mapping resolution improves with increasing numbers of strains. (B) With the QTL effect size again fixed at 50% and the number of strains fixed at 72, the distribution of distances does not appear to differ based on the number of functional alleles. (C) With the number of strains fixed at 72 and the number of alleles fixed at 8, as the QTL effect size increase, the distribution of distances becomes more concentrated around zero. These simulations are based on Definition B and single observations per strain. See **Figures 4** and **S3** for specific estimates of location error over different settings of QTL effect size and numbers of strains.
