## Supplementary material for "Determinants of QTL mapping power in the realized Collaborative Cross": Example shell script to coordinate separate calls to sparcc_powersim.R: sparcc_powersim.sh

```
#!/bin/bash

#the full SPARCC parameter settings thatare commented out; in order to re-generate these we highly recommend parallelizing the underlying Rscript call (e.g. bsub)
#currently, this script is set up to generate a subset of the results, which can be run on a single machine but will still take a considerable amount of time

#output directory for each batch of jobs
mkdir -p results

#specify number of strains
#or i in `seq 10 5 70` 72; do
for i in 72; do
        #specify number of replicates
        #only need to simulate single replicate, can infer other values
        for j in 1; do
                #specify number of alleles
                #for k in 2 3 8; do
                for k in 8; do
                        #specify variance explained by qtl
                        #for l in 0.01 `seq -w 0.05 0.05 0.96`; do
                        for l in 0.5; do
                                #specify variance explained by strain
                                #only need to simulate 0; can infer other values
                                for m in 0; do
                                        #specify number of jobs per parameter setting
                                        #each job includes 100 simulations
                                        for n in `seq 1 10`; do
                                                job_id="powersim_n.strains_"$i"_n.reps_"$j"_n.alleles_"$k"_h.qtl_"$l"_h.strain_"$m"_job_"$n
                                                #run Rscript job only if the result file does not already exist
                                                if [ ! -f "./results/"$job_id".RData" ]; then
                                                        #run Rscript job OR submit to job management queue; optionally add a seed to replace NA for the final argument
                                                        Rscript sparcc_powersim.R "$job_id" ./CC_genomecache_l2norm0.1_build37/ "$i" "$j" 100 "$k" "$l" "$m" NA
                                                        #example job submission using bsub
                                                        #bsub "Rscript sparcc_powersim.R "$job_id" ./CC_genomecache_l2norm0.1_build37/ "$i" "$j" 100 "$k" "$l" "$m" NA"
                                                fi
                                        done
                                done
                        done
                done
        done
done


#after all of the previous result files have finished running, run the following Rscript which collates all of the results to "full_results.RData"
#make sure that the parameter settings (specified in the loops of this script) match the specifidations at the start of the following script
Rscript collapse_results.R
```
