## Supplementary material for "Determinants of QTL mapping power in the realized Collaborative Cross": Provides description of options in SPARCC, and provides simple tutorial for using the package

### SPARCC code supplement

#### SPARCC package options

##### *Phenotype simulation*

The output of the CC data simulation is a matrix of outcomes, where each column of $\mathbf{y}^{(s)}$ is the phenotype generated by the following equation:

$$\mathbf{y} = \mathbf{1}\mu + \underbrace{\mathbf{ZX}\boldsymbol{\beta}}_{\text{QTL effect}} + \underbrace{\mathbf{Zu}}_{\text{Strain effect}} + \underbrace{\boldsymbol{\varepsilon}}_{\text{Noise}},$$

where $\mathbf{1}$ is an $N$-vector of 1's, $\mu$ is an intercept, $\mathbf{Z}$ is an $N \times n$ incidence matrix mapping individuals to strains, $\mathbf{X}$ is an $n \times m$ allele dosage matrix mapping strains to their estimated dosage of each of the $m$ alleles, $\boldsymbol{\beta}$ is an $m$-vector of allele effects, $\mathbf{u}$ is an $n$-vector of strain effects (representing polygenic background variation), and $\boldsymbol{\varepsilon}$ is an $N$-vector of unstructured, residual error. The matrix of phenotypes is simulated using the `sim.CC.data()` function. The following options control various aspects of the simulation and the simulated QTL.

##### *QTL effect:*

- `qtl.effect.size`
  - $0 \leq$ `qtl.effect.size` $< 1 -$ `strain.effect.size`
  - This argument represents $h^2_{\text{QTL}}$, such that $\boldsymbol{\beta} \sim \text{N}(\mathbf{0}, \mathbf{I}h^2_{\text{QTL}})$.
  - A specific $\boldsymbol{\beta}$ can specified with the `beta` argument, though it will be scaled to match `qtl.effect.size`. If `beta=NULL`, then $\boldsymbol{\beta}$ is sampled accordingly.
- `num.alleles` (DEFAULT = 8)
  - $2 \leq$ `num.alleles` $\leq 8$
- `M.ID`
  - Rather than specifying `num.alleles` and then sampling $\mathbf{M}$, these can be fixed with the `M.ID` argument.
  - Expects strings of the form `"A,B,C,D,E,F,G,H"`, with each letter corresponding to a founder strain, taking an integer value 0-7, representing functional alleles.
  - Example: `M.ID="0,0,0,0,1,1,1,1"` represents a bi-allelic causal variant, in which the first four strains have one allele, and the last four having the other.
- `CC.lines` (DEFAULT = NULL)
  - This argument allows the user to provide a vector of CC strain IDs on which to base the power calculation. The CC genomes, along with `locus`, will determine $\mathbf{D}$ in the following equation: $\mathbf{X} = \mathbf{DAM}$.
  - If `CC.lines = NULL`, then SPARCC will sample `num.lines` from all available strains.
    - `vary.lines` (DEFAULT = TRUE)
      - If `vary.lines = TRUE`, the set of strains for each simulation will be sampled and vary.
      - If `vary.lines = FALSE`, the set of strains will be sampled once, and used for each simulated outcome.
- `locus` (DEFAULT = NULL)
  - This argument allows the user to specify a specific locus for the simulated QTL, in effect determining the haplotype dosage matrix $\mathbf{D}$.
  - If the argument is left empty, SPARCC will sample loci uniformly from the CC genomes, thus providing power estimates averaged over genomic positions.
- `impute` (DEFAULT = TRUE)
  - If `impute=TRUE`, then $\mathbf{D}$ is sampled from the probabilistically reconstructed diplotypes at the QTL

    $$\mathbf{D}_i \sim \text{Cat}(\mathbf{P}_i)$$

    where Cat(.) is a categorical distribution and $\mathbf{P}$ is a matrix of diplotype probabilities for the sampled CC genomes at the QTL.
  - If `impute=FALSE`, then $\mathbf{D} = \mathbf{P}$ in the simulation procedure.
- `scale.qtl.mode` (DEFAULT = "B")
  - If `scale.qtl.mode="B"`, $\text{V}(2\boldsymbol{\beta})$ is scaled to `qtl.effect.size`, where var is the maximum likelihood estimate of variance rather than sample variance, such that var $= \frac{n-1}{n}s$, where $s^2$ is the sample variance and $n$ is the number of individuals. This scaling sets the QTL effect size with respect to a theoretical population that is evenly balanced with respect to functional alleles.

- If `scale.qtl.mode="MB"`, $\mathrm{V}(2\mathbf{M}\boldsymbol{\beta})$ is scaled to `qtl.effect.size`, setting the QTL effect size to a theoretical population with a specific frequency of functional alleles among the founder strains. The effect size here is dependent on $\mathbf{M}$ but independent of $\mathbf{D}$, and the proportion of variance explained by the QTL in the mapping population will deviate from $h^2_{\mathrm{QTL}}$ due to imbalance $\mathbf{D}$ but not in $\mathbf{M}$.
- If `scale.qtl.mode="DAMB"`, $\mathrm{V}(\mathbf{DAM}\boldsymbol{\beta})$ is scaled to `qtl.effect.size`, setting the QTL effect size to a specific set of CC strains and allele series.
- If `scale.qtl.mode="none"`, $\boldsymbol{\beta}$ is not scaled, allowing the user to specify an effect vector and not have it modified.

***Strain effect:***

- `strain.effect.size`
  - $0 \leq$ `strain.effect.size` $\leq 1 -$ `qtl.effect.size`
  - This argument specifies $h^2_{\mathrm{strain}}$, such that $\mathbf{u} \sim \mathrm{N}(\mathbf{0}, \mathbf{I}h^2_{\mathrm{strain}})$.

***Additional options:***

- `num.sim`
  - This argument specifies SPARCC to simulate $s$ samples of $\mathbf{y}$.
- `num.replicates`
  - This argument allows the user to set the number $r$ of replicates of each CC strain. The reproducibility of CC genomes is an important feature, allowing noise variation to be reduced.
  - SPARCC currently requires all CC lines to have the same number of replicates.

***Genome scans***

Each phenotype from the simulation procedure is then evaluated via QTL mapping. The SPARCC function for running genome scans from the simulated data is `run.sim.scans()`, The primary argument is `sim.data`. There are additional arguments to restrict the scans to a subset of the chromosomes, to a subset of the simulated phenotypes, or to a subset of loci. Last, the user can provide the pre-computed QR decompositions and specify whether the output should return those decompositions for further use with additional simulations, although this can be expensive in terms of memory.

#### Simple tutorial for SPARCC simulation, scans, and power

This example is computationally efficient because the CC strains are not varied across simulations. Varying CC strains increases the computational expense, which was done for the reported results. Generates **Figure 2** from the manuscript.

```
###----------------------------------------------------
### Useful functions for parsing haplotype data
> h <- DiploprobReader$new("./sparcc_cache/")
> set.seed(100)

### Grabbing random sample of 65 CC strains
> these.cc.lines <- sample(h$getSubjects(), size=65)

> library(sparcc)
### Simulate 5 data sets:
#### Specified 65 CC strains
#### 5 replicates of each
#### 2 functional alleles, allelic series not specified
#### QTL effect size of 10
#### Background Strain effect of 0
> simple.data <- sim.CC.data(genomecache="./sparcc_cache/",
                             CC.lines=these.cc.lines,
                             num.replicates=5,
                             num.sim=5,
                             num.alleles=2,
                             qtl.effect.size=0.1,
                             strain.effect.size=0)

### Genome scans
> simple.scans <- run.sim.scans(sim.data=simple.data,
                                return.all.sim.qr=TRUE)

### Generating permutation index
```

```
> perm.index <- generate.perm.matrix(num.lines=65,
                                     num.perm=100)

### Permutation scans
> thresh.scans <- run.perm.scans(perm.matrix=perm.index,
                                 sim.CC.object=simple.data,
                                 sim.CC.scans=simple.scans)

### Calculating significance thresholds
> all.thresh <- get.thresholds(thresh.scans=thresh.scans)

### Power estimate
> pull.power(sim.scans=simple.scans,
 thresh=all.thresh)
[1] 0.8

### Plot a genome scan of a single simulated phenotype
> single.sim.plot(simple.scans,
                thresh=all.thresh,
                phenotype.index=1)
###-----------------------------------------------------
```

###### *Run-time performance of tutorial*

The simple example was run locally on an Early 2015 MacBook Pro with a 2.9 GHz Intel Core i5 processor and 8 GB of RAM. The data simulation and genome-wide scans for five phenotypes took 34.13 seconds. Computational time increases linearly with number of phenotypes simulated. Computational times will also decrease for lower numbers of CC strains. 100 permutations for 5 simulated phenotypes took 8.73 minutes. Although the time expense for these simulations is not trivial, the overall process is highly optimized; this simple example involves fitting 5 phenotypes $\times$ 17900 loci $\times$ 100 permutation alternative models. The process can be sped up using a parallel computing environment, as we do with the following large scale analysis. Highly specific power calculations for an experiment are feasible on a local computer using a single core.

##### Simple tutorial for SPARCC power projection

This example code demonstrates how to calculate power estimates from the dataset stored within SPARCC using projection and interpolation. This code produces **Figure S1** in the **Supplement**.

```
###-----------------------------------------------------
### r1.dat is included in SPARCC
### Project and interpolate power estimates for
### experiments with 3 replicates
> r3.interp.dat <- interpolate.table(r1.results=r1.dat,
                                     num.replicates=3,
                                     n.alleles=2)
### Project and interpolate power estimates for
### experiments with 5 replicates
> r5.interp.dat <- interpolate.table(r1.results=r1.dat,
                                      num.replicates=5,
                                      n.alleles=2)
### Plotting power curves
> power.plot(results=r1.dat,
           qtl.effect.size=0.3,
           n.alleles=2)
> add.curve.to.power.plot(results=r3.interp.dat,
                        qtl.effect.size=0.3,
                        n.alleles=2)
> add.curve.to.power.plot(results=r5.interp.dat,
                        qtl.effect.size=0.3,
                        n.alleles=2)
###-----------------------------------------------------
```
