## Supplementary material for "Determinants of QTL mapping power in the realized Collaborative Cross": SPARCC R package used for all analyses: DESCRIPTION

Package: sparcc
Title: Simulated Power Analysis in the Realized Collaborative Cross
Version: 1.1.1
	     person("Wesley", "Crouse", role = "aut"),
	     person("William", "Valdar", role = "aut"))
Description: This package provides functionality to simulate phenotypes in the actual Collaborative Cross (CC) lines, which can be used by other tools designed to analyze CC data. Additionally, it provides rapid QTL mapping for the estimation of power to map QTL, and other statistics of interest, such as false positive rates.
Depends: R (>= 3.2.2)
License: GPL (>= 2)
Encoding: UTF-8
LazyData: true
RoxygenNote: 6.1.0
Imports: lme4, akima
NeedsCompilation: no
Packaged: 2019-02-25 20:53:02 UTC; Greg
Author: Gregory Keele [aut, cre],
  Wesley Crouse [aut],
  William Valdar [aut]
Maintainer: Gregory Keele <>
