## Supplementary material for "Determinants of QTL mapping power in the realized Collaborative Cross": SPARCC R package used for all analyses: CC_extra.R

```
#' Calculate the QTL mapping power from SPARCC genome scans output from run.sim.scans()
#'
#' This function takes output genome scans from run.sim.scans() and calculates the power to map the simulated
#' QTL. A window around the QTL can be specified to allow peaks not immediately at the simulated locus but likely
#' tagging the QTL to also be included.
#'
#' @param sim.scans Output simulated genome scans from run.sim.scans().
#' @param thresh A list of threshold, calculated from get.gev.thresh(), that correspond to the scans in sim.scans. 
#' @param window.mb DEFAULT: 5. Loci upstream and downstream the specified window.mb in Mb will also be checked 
#' for statistically significant signals. Sometimes the statistical score will not pass at the simulated QTL, but
#' does at nearby loci.
#' @export pull.power
#' @examples pull.power()
pull.power <- function(sim.scans, thresh, window.mb=5){
  map <- rep(NA, nrow(sim.scans$p.value))
  for (i in 1:nrow(sim.scans$p.value)) {
    this.index <- which(colnames(sim.scans$p.value) == sim.scans$locus[i])
    this.chr <- sim.scans$chr[this.index]
    this.pos <- sim.scans$pos$Mb[this.index]
    p.value <- sim.scans$p.value[i, sim.scans$pos$Mb >= this.pos - window.mb & sim.scans$pos$Mb <= this.pos + window.mb & sim.scans$chr == this.chr]
    map[i] <- any(-log10(p.value) >= thresh[i])
  }
  return(mean(map))
}

#' Calculate the probability of detecting any false QTL from SPARCC genome scans output from run.sim.scans()
#'
#' This function takes output genome scans from run.sim.scans() and calculates the probability that any false QTL 
#' are detected. A window around the QTL can be specified to allow peaks not immediately at the simulated locus but likely
#' tagging the QTL to also be excluded
#'
#' @param sim.scans Output simulated genome scans from run.sim.scans().
#' @param thresh A list of threshold, calculated from get.gev.thresh(), that correspond to the scans in sim.scans. 
#' @param window.mb DEFAULT: "chromosome". Loci upstream and downstream the specified window.mb in Mb will also be checked 
#' for statistically significant signals. Sometimes the statistical score will not pass at the simulated QTL, but
#' does at nearby loci. "chromosome" results in only the consideration of signals from off-site chromosomes.
#' @export pull.false.positive.prob
#' @examples pull.false.positive.prob()
pull.false.positive.prob <- function(sim.scans, thresh, window.mb="chromosome"){
  map <- rep(NA, nrow(sim.scans$p.value))
  total <- length(sim.scans$p.value)
  for (i in 1:nrow(sim.scans$p.value)) {
    this.index <- which(colnames(sim.scans$p.value) == sim.scans$locus[i])
    this.chr <- sim.scans$chr[this.index]
    if (window.mb == "chromosome") {
      p.value <- sim.scans$p.value[i, !(sim.scans$chr == this.chr)]
    }
    else {
      this.pos <- sim.scans$pos$Mb[this.index]
      p.value <- sim.scans$p.value[i, !(sim.scans$pos$Mb >= this.pos - window.mb & sim.scans$pos$Mb <= this.pos + window.mb & sim.scans$chr == this.chr)]
    }
    map[i] <- any(-log10(p.value) >= thresh[i])
  }
  return(mean(map))
}

#' Calculate the distance between the simulated QTL and the minimum p-value locus within a window around the simulated QTL, given
#' a QTL was detected in the window from SPARCC genome scans output from run.sim.scans()
#'
#' This function takes output genome scans from run.sim.scans() and calculates the distance between the simulated QTL and the 
#' minimum p-value locus within a window around the simulated QTL if a QTL was detected. The intent is to quantify the distribution
#' of the position of max statistical signal from the simulated QTL position.
#'
#' @param sim.scans Output simulated genome scans from run.sim.scans().
#' @param thresh A list of threshold, calculated from get.gev.thresh(), that correspond to the scans in sim.scans. 
#' @param window.mb DEFAULT: 5. Loci upstream and downstream the specified window.mb in Mb will also be checked 
#' for statistically significant signals. Sometimes the statistical score will not pass at the simulated QTL, but
#' does at nearby loci. This is the region interrogated for signals greater than the simulated QTL.
#' @export pull.dist.from.locus
#' @examples pull.dist.from.locus()
pull.dist.from.locus <- function(sim.scans, thresh, window.mb=5) {
  map <- rep(NA, nrow(sim.scans$p.value))
  total <- length(sim.scans$p.value)
  for (i in 1:nrow(sim.scans$p.value)) {
    this.index <- which(colnames(sim.scans$p.value) == sim.scans$locus[i])
    this.chr <- sim.scans$chr[this.index]
    this.pos <- sim.scans$pos$Mb[this.index]
    p.value <- sim.scans$p.value[i, sim.scans$pos$Mb >= this.pos - window.mb & sim.scans$pos$Mb <= this.pos + window.mb & sim.scans$chr == this.chr]
    pos <- sim.scans$pos$Mb[sim.scans$pos$Mb >= this.pos - window.mb & sim.scans$pos$Mb <= this.pos + window.mb & sim.scans$chr == this.chr]
    if (any(-log10(p.value) >= thresh[i])) {
      map[i] <- pos[which.min(p.value)] - this.pos
    }
    else {
      map[i] <- NA
    }
  }
  return(map)
}

#' Calculates the QTL effect size in a population of means based on the reduction 
#' in residual error due to replicates
#'
#' This function calculates the QTL effect size in the mapping population on strain means based on the reduction
#' in random variation due to noise that results from averaging the phenotypes of replicates.
#'
#' @param qtl.effect.size The proportion of variance due to the QTL in the sample population with replicates.
#' Must be between 0 and 1.
#' @param strain.effect.size DEFAULT: 0. The proportion of varianced due to background strain effect. This source of 
#' noise, with respect to the QTL signal, cannot be reduced by replicate observation.
#' @param num.replicates The number of replicates per CC strain. Higher numbers will result in greater reduction of
#' random error, and larger increases in QTL effect size in the sample population of means.
#' @export convert.qtl.effect.to.means
#' @examples convert.qtl.effect.to.means()
convert.qtl.effect.to.means <- function(qtl.effect.size,
                                        strain.effect.size=0,
                                        num.replicates){
  mean.noise.effect.size <- (1 - qtl.effect.size - strain.effect.size)/num.replicates
  denominator <- sum(qtl.effect.size, strain.effect.size, mean.noise.effect.size)
  mean.qtl.effect.size <- qtl.effect.size/denominator
  mean.strain.effect.size <- strain.effect.size/denominator
  results <- c(mean.qtl.effect.size, mean.strain.effect.size)
  names(results) <- c("QTL", "strain")
  return(results)
}

non.sample.var <- function(x) {
  var.x <- var(x)*((length(x) - 1)/length(x))
  return(var.x)
}

#' Interpolates QTL mapping power for sample populations with replicates from dense results 
#' in single observation simulations
#'
#' This function interpolates the QTL mapping power for an experimental mapping population with replicates
#' based on dense simulated power results in simulations with only a single observation per strain. 
#'
#' @param r1.results Data frame of power estimates from simulations based on a single observation per strain.
#' r1.dat and r1.damb.dat are included in SPARCC for this purpose.
#' @param qtl.effect.sizes The desired QTL effect sizes for which power estimates will be interpolated from dense
#' single observation results.
#' @param strain.effect.sizes The desired proportions of variance due to background strain effect for hypothetical population in which
#' power is evaluated.
#' @param num.replicates The desired number of replicates for a hypothetical study in which to evaluate power.
#' @param n.alleles The number of functional alleles for the simulated QTL.
#' @param use.window DEFAULT: TRUE. Whether the interpolated powers are based on denser power estimates using a window
#' around a locus or just the actual locus.
#' @param n.strains The number of CC strains for the desired interpolated power estimate.
#' @export interpolate.qtl.power
#' @examples interpolate.qtl.power()
interpolate.qtl.power <- function(r1.results,
                                  qtl.effect.sizes,
                                  strain.effect.sizes=0,
                                  num.replicates,
                                  n.alleles,
                                  use.window=TRUE,
                                  n.strains) {
  
  if (length(strain.effect.sizes) == 1) { strain.effect.sizes <- rep(strain.effect.sizes, length(qtl.effect.sizes)) }
  if (length(num.replicates) == 1) { num.replicates <- rep(num.replicates, length(qtl.effect.sizes)) }
  r1.qtl.effect.sizes <- sapply(1:length(qtl.effect.sizes), function(x) convert.qtl.effect.to.means(qtl.effect.size=qtl.effect.sizes[x],
                                                                                                    strain.effect.size=strain.effect.sizes[x],
                                                                                                    num.replicates=num.replicates[x])["QTL"])
  ## Processing evaluated power
  power <- r1.results[r1.results$n.strains %in% n.strains & r1.results$n.alleles %in% n.alleles,]
  if (use.window) { y <- power$power }
  else { y <- power$power.window }
  x <- as.numeric(as.character(power$h.qtl))
  
  y <- c(0, y, 1)
  x <- c(0, x, 1)
  
  powers <- approx(x=x, y=y, xout=r1.qtl.effect.sizes)$y
  return(powers)
}

#' Interpolates QTL location error for sample populations with replicates from dense results 
#' in single observation simulations
#'
#' This function interpolates the QTL location error for an experimental mapping population with replicates
#' based on dense simulated power results in simulations with only a single observation per strain. 
#'
#' @param r1.results Data frame of power estimates from simulations based on a single observation per strain.
#' r1.dist.dat is included in SPARCC for this purpose.
#' @param qtl.effect.sizes The desired QTL effect sizes for which power estimates will be interpolated from dense
#' single observation results.
#' @param strain.effect.sizes DEFAULT: 0. The desired proportions of variance due to background strain effect for hypothetical population in which
#' power is evaluated.
#' @param num.replicates The desired number of replicates for a hypothetical study in which to evaluate power.
#' @param n.alleles The number of functional alleles for the simulated QTL.
#' @param n.strains The number of CC strains for the desired interpolated power estimate.
#' @param n.mean.psuedo DEFAULT: 10. The number of psuedo observations (of 2.5) to be used for regularization.
#' @export interpolate.qtl.distance
#' @examples interpolate.qtl.distance()
interpolate.qtl.distance <- function(r1.results,
                                     qtl.effect.sizes,
                                     strain.effect.sizes = 0,
                                     num.replicates,
                                     n.alleles,
                                     n.strains,
                                     n.mean.pseudo = 10) {
  
  if (length(strain.effect.sizes) == 1) { strain.effect.sizes <- rep(strain.effect.sizes, length(qtl.effect.sizes)) }
  if (length(num.replicates) == 1) { num.replicates <- rep(num.replicates, length(qtl.effect.sizes)) }
  r1.qtl.effect.sizes <- sapply(1:length(qtl.effect.sizes), function(x) convert.qtl.effect.to.means(qtl.effect.size=qtl.effect.sizes[x],
                                                                                                    strain.effect.size=strain.effect.sizes[x],
                                                                                                    num.replicates=num.replicates[x])["QTL"])
  ## Processing evaluated power
  dist.tab <- r1.results[r1.results$n.strains %in% n.strains & r1.results$n.alleles %in% n.alleles,]
  #y <- abs(dist.tab$dist)
  #browser()
  y <- sapply(1:length(dist.tab$dist),
              function(i) mean(c(abs(dist.tab$dist[i]), rep(2.5, n.mean.pseudo))))
  x <- as.numeric(as.character(dist.tab$h.qtl))
  
  y <- c(2.5, y, 0)
  x <- c(0, x, 1)

  distances <- approx(x = x, y = y, xout = r1.qtl.effect.sizes)$y
  return(distances)
}


#' Convenience function that interpolates QTL mapping power based on QTL effect sizes in a results data frame 
#' from single observation simulations
#'
#' This function interpolates the QTL mapping power through interpolate.qtl.power() based on a results data frame
#' of densely simulated power estimates in popuations with a single observation per strain. 
#'
#' @param r1.results Data frame of power estimates from simulations based on a single observation per strain.
#' r1.dat and r1.damb.dat are included in SPARCC for this purpose.
#' @param num.replicates The desired number of replicates for a hypothetical study in which to evaluate power.
#' @param strain.effect.size DEFAULT: NULL. The desired proportion of variance due to background strain effect for hypothetical population in which
#' power is evaluated.
#' @param n.alleles The number of functional alleles for the simulated QTL.
#' @param use.window DEFAULT: TRUE. Whether the interpolated powers are based on denser power estimates using a window
#' around a locus or just the actual locus.
#' @export interpolate.table
#' @examples interpolate.table()
interpolate.table <- function(r1.results,
                              num.replicates,
                              strain.effect.size=NULL,
                              n.alleles,
                              use.window=TRUE) {
  qtl.effect.size <- unique(r1.results$h.qtl)
  n.strains <- unique(r1.results$n.strains)
  if (is.null(strain.effect.size)) { strain.effect.size <- unique(r1.results$h.strain) }
  final.data <- NULL
  for(i in 1:length(n.strains)) {
    temp <- sapply(1:length(num.replicates), function(x) interpolate.qtl.power(r1.results=r1.results,
                                                                               qtl.effect.sizes=qtl.effect.size,
                                                                               strain.effect.size=strain.effect.size,
                                                                               num.replicates=num.replicates[x],
                                                                               n.alleles=n.alleles,
                                                                               use.window=use.window,
                                                                               n.strains=n.strains[i]))
    rownames(temp) <- qtl.effect.size
    colnames(temp) <- num.replicates
    temp.data <- reshape2::melt(temp)
    temp.data <- cbind(rep(n.strains[i], nrow(temp.data)), rep(n.alleles, nrow(temp.data)), temp.data)
    names(temp.data) <- c("n.strains", "n.alleles", "h.qtl", "n.replicates", ifelse(use.window, "power.window", "power"))
    final.data <- rbind(final.data, temp.data)
  } 
  return(final.data)
}

#function for curve point confidence interval using binomial with jeffery's prior
#' @export binomial.prop.ci
binomial.prop.ci <- function(p, n.sims=1000, alpha=0.05){
  ci.lower <- qbeta(0.5*alpha, p*n.sims+0.5, (1-p)*n.sims+0.5)
  ci <- c(ci.lower, qbeta(1-0.5*alpha, p*n.sims+0.5, (1-p)*n.sims+0.5))
  
  if (p==0){
    ci[1] <- 0
  } else if (p==1){
    ci[2] <- 1
  }
  return(ci)
}

# Originally from miqtl
get.f.stat.p.val <- function(qr.alt, 
                             qr.null, 
                             y){
  rss0 <- sum(qr.resid(qr.null, y)^2)
  rss1 <- sum(qr.resid(qr.alt, y)^2)
  df1 <- qr.alt$rank - qr.null$rank
  df2 <- length(y) - qr.alt$rank
  
  mst <- (rss0 - rss1)/df1
  mse <- rss1/df2
  f.stat <- mst/mse
  p.val <- pf(q=f.stat, df1=df1, df2=df2, lower.tail=FALSE)
  return(p.val)
}
```
