## Supplementary material for "Determinants of QTL mapping power in the realized Collaborative Cross": SPARCC R package used for all analyses: CC_perm.R

```
#' Extract GEV thresholds from permutation scans from SPARCC simulations
#' 
#' This function takes the output from run.perm.scans() and calculates GEV thresholds per simulated phenotype.
#' 
#' @param thresh.scans The output object from run.perm.scans(). Is simply a matrix of minimum p-values per
#' simulatione phenotype per permutation.
#' @param percentile DEFAULT: 0.95. Specifies that 1 - alpha level for significance.
#' @export
#' @examples get.thresholds()
get.thresholds <- function(thresh.scans, 
                           percentile=0.95){
  
  extreme.values <- -log10(thresh.scans)

  thresh.vec <- rep(NA, nrow(thresh.scans))
  
  for (i in 1:nrow(thresh.scans)) {
    evd.pars <- as.numeric(evir::gev(extreme.values[i,])$par.est)
    thresh.vec[i] <- evir::qgev(p=percentile, xi=evd.pars[1], sigma=evd.pars[2], mu=evd.pars[3])
  }
  return(thresh.vec)
}

#' Generates a matrix of permutation indeces. Can be used across simulations with the same number of CC lines
#'
#' This function takes the output from sim.CC.data() to produce a matrix of permutation indeces, allowing
#' the same permutations to be performed for different phenotypes but the same CC lines.
#'
#' @param sim.CC.object Simulated CC data output from sim.CC.data().
#' @param num.perm The number of permutations.
#' @export
#' @examples generate.perm.matrix()
generate.perm.matrix <- function(num.lines, 
                                 num.perm){
  perm.matrix <- replicate(n=num.perm, sample(1:num.lines, replace=FALSE))
  colnames(perm.matrix) <- paste0("perm.", 1:num.perm)
  return(perm.matrix)
}

#' Runs permutation scans from a permutation index matrix, simulated CC data, and simulated CC scans.
#'
#' This function takes the outputs from generate.perm.matrix(), sim.CC.data(), and run.sim.scans() to perform
#' permutation scans and determine a significance threshold. 
#'
#' @param perm.matrix Permutation index matrix that is output from generate.perm.matrix().
#' @param sim.CC.scans Genome scans from simulated CC data output from run.sim.scans().
#' @param sim.CC.object Simulated CC data output from sim.CC.data().
#' @param phenotype.index The phenotype index of simulated phenotype that correspond to those found in sim.CC.scans
#' and sim.CC.objects.
#' @param all.sim.qr DEFAULT: NULL. Allows qr decompositions to only be saved once. If NULL, it expects that they are
#' stored in sim.CC.scans$all.sim.qr, which can be specified in run.sim.scans().
#' @param scan.index DEFAULT: NULL. If NULL, it performs scans for all permuted data sets based on permutation index.
#' If given a vector of integers, representing the permutation index, it will only run scans for those permutations.
#' @param chr DEFAULT: "all". Specifies which chromosomes to scan.
#' @param just.these.loci DEFAULT: NULL. If NULL, all loci in genome cache are scanned.
#' @param use.progress.bar DEFAULT: FALSE. Specifies whether to use a progress bar for qr decompositions 
#' and genome scans. 
#' @param print.scans.progress DEFAULT: FALSE. Specifies whether to output a message that indicates the number of permutation
#' scans completed. 
#' @export
#' @examples run.perm.scans()
run.perm.scans <- function(perm.matrix, 
                           sim.CC.scans, 
                           sim.CC.object,
                           phenotype.index=NULL,
                           all.sim.qr=NULL,
                           scan.index=NULL,
                           chr="all", 
                           just.these.loci=NULL, 
                           use.progress.bar=FALSE,
                           print.scans.progress=FALSE,
                           ...){
  
  if (is.null(scan.index)) { scan.index <- 1:ncol(perm.matrix) }
  if (is.null(phenotype.index)) { phenotype.index <- 1:sum(grepl(colnames(x=sim.CC.object$data), pattern="sim.y")) }
  
  is.full <- ifelse(is.null(all.sim.qr), TRUE, FALSE)
  if (is.null(all.sim.qr)) {
    all.sim.qr <- sim.CC.scans$all.sim.qr
  }

  if (sim.CC.scans$properties$vary.lines) {
    loci <- names(all.sim.qr[[1]]$qr.list)
    rh.formula <- all.sim.qr[[1]]$formula
    loci.chr <- all.sim.qr[[1]]$chr
  }
  else {
    loci <- names(all.sim.qr$qr.list)
    rh.formula <- all.sim.qr$formula
    loci.chr <- all.sim.qr$chr
  }
  
  if (chr[1] != "all") {
    loci <- loci[loci.chr %in% chr]
  }
  if (!is.null(just.these.loci)) {
    loci <- loci[loci %in% just.these.loci]
    loci.chr <- loci.chr[loci %in% just.these.loci]
  }
  
  min.p <- matrix(NA, nrow=length(phenotype.index), ncol=length(scan.index))
  
  for (i in 1:length(phenotype.index)) {
    data <- sim.CC.object$data[,c(paste0("sim.y.", phenotype.index[i]), 
                                  paste0("SUBJECT.NAME.", ifelse(sim.CC.scans$properties$vary.lines, phenotype.index[i], 1)))]
    if(sim.CC.scans$properties$vary.lines){
      if(is.full){
        this.qr <- all.sim.qr[[phenotype.index[i]]]
      }
      else{
        this.qr <- all.sim.qr
      }
    }
    else{
      this.qr <- all.sim.qr
    }
    for(j in 1:length(scan.index)){
      perm.data <- data
      perm.data[,1] <- data[perm.matrix[,scan.index[j]], 1]
      names(perm.data) <- c("perm_y", "SUBJECT.NAME")
      
      this.scan <- scan.qr(qr.object=this.qr, data=perm.data, 
                           phenotype="perm_y", chr=chr,
                           return.allele.effects=FALSE, use.progress.bar=use.progress.bar,
                           ...)

      min.p[i,j] <-  min(this.scan$p.value)
      if(print.scans.progress){
        cat("\n", "Threshold scan: phenotype index =", phenotype.index[i], "----",
            "perm index =", scan.index[j], "---- out of", length(scan.index), "permutations ---- out of", length(phenotype.index), "phenotypes\n")
      }
    }
  }
  return(min.p)
}
```
