## Supplementary material for "Determinants of QTL mapping power in the realized Collaborative Cross": SPARCC R package used for all analyses: CC_plot.R

```
#' Plots a single simulated scan. Built on genome.plotter.whole(), originally from the R package miqtl
#'
#' This function takes the output from run.sim.scans(), and plots a single genome scane of a simulated phenotype.
#'
#' @param sim.scans Genome scans of simulated data, output by run.sim.scans().
#' @param phenotype.index DEFAULT: 1. The index of the simulated phenotype to plot. Cannot exceed the number of 
#' phenotypes contained in sim.scans.
#' @param scale DEFAULT: "Mb". The scale in which to plot genomic position. "cM" is the other option.
#' @param thresh DEFAULT: NULL. If NULL, no significance is threshold is plotted. Expects a numeric value, commonly
#' output from get.gev.thresholds() based on permutation scans.
#' @export
#' @examples single.sim.plot()
single.sim.plot <- function(sim.scans, 
                            phenotype.index=1, 
                            scale=c("Mb", "cM"), 
                            thresh=NULL, 
                            ...){
  
  scale <- scale[1]
  dummy.scan <- sim.scans
  if(is.matrix(dummy.scan$p.value)){
    dummy.scan$p.value <- sim.scans$p.value[phenotype.index,]
  }
  else{
    dummy.scan$p.value <- sim.scans$p.value
  }
  dummy.scan$p.value[dummy.scan$p.value == 0] <- .Machine$double.xmin
  
  if(length(sim.scans$locus) > 1){
    locus <- sim.scans$locus[phenotype.index]
  }
  else{
    locus <- sim.scans$locus
  }
  
  outcome.type <- NULL
  if (sim.scans$properties$return.value != "raw") { outcome.type <- sim.scans$properties$return.value }
  
  if (!is.null(sim.scans$properties$strain.reduction.fraction)) { 
    effect.title <- paste(paste0("QTL=",  sim.scans$properties$qtl.effect.size*100, "%"),
                          paste0("Strain=", sim.scans$properties$strain.effect.size*100, "%"),
                          paste0("Strain reduction=", round(sim.scans$properties$strain.reduction.fraction*100, 2), "%"),
                          sep=", ")
  } 
  else {
    effect.title <- paste(paste0("QTL=",  sim.scans$properties$qtl.effect.size*100, "%"),
                          paste0("Strain=", sim.scans$properties$strain.effect.size*100, "%"),
                          sep=", ")
  }
  
  this.title <- c(paste(outcome.type, 
                        ifelse(sim.scans$properties$return.means, "means:", "replicates:"),
                        effect.title),
                  paste(ifelse(sim.scans$properties$impute, "Impute", "ROP"), "sim,",
                        sim.scans$properties$num.alleles, "functional alleles,",
                        sim.scans$properties$num.lines, "lines,",
                        sim.scans$properties$num.replicates, ifelse(sim.scans$properties$num.replicates == 1, "replicate", "replicates")))
  if (sim.scans$properties$qtl.effect.size == 0) { locus <- NULL }
  genome.plotter.sparcc(scan.list=list(dummy.scan), override.title=this.title, distinguish.chr.type="color",
                        scale=scale, mark.locus=locus, hard.thresholds=thresh[phenotype.index], ...)
}

#' @export
sim.data.effect.plot <- function(sim.data,
                                 x.var=c("B.effect", "MB.effect", "DAMB.effect", "ZDAMB.effect"),
                                 y.var=c("MB.effect", "DAMB.effect", "ZDAMB.effect", "B.effect"),
                                 qtl.col="red", title="") {
  x.var <- x.var[1]
  y.var <- y.var[1]
  nominal.effect.size <- sim.data$properties$qtl.effect.size
  var.table <- sim.data$properties$var.table
  
  x.val <- var.table[,x.var]
  y.val <- var.table[,y.var]
  
  x.max <- max(x.val, nominal.effect.size)
  y.max <- max(y.val, nominal.effect.size)
  
  plot(x.val, y.val,
       xlab=x.var, ylab=y.var, xlim=c(0, x.max), ylim=c(0, y.max),
       frame.plot=FALSE, pch=20, las=1, main=title)
  abline(0, 1, lty=3)
  abline(v=nominal.effect.size, lty=2, col=qtl.col)
  abline(h=nominal.effect.size, lty=2, col=qtl.col)
}

#' @export
sim.data.ve.plot <- function(sim.data,
                             x.var=c("B.ve", "MB.ve", "DAMB.ve"),
                             y.var=c("MB.ve", "DAMB.ve", "B.ve"),
                             qtl.col="red", title="") {
  x.var <- x.var[1]
  y.var <- y.var[1]
  nominal.effect.size <- sim.data$properties$qtl.effect.size
  var.table <- sim.data$properties$var.table
  
  x.val <- var.table[,x.var]
  y.val <- var.table[,y.var]
  
  plot(x.val, y.val,
       xlab=x.var, ylab=y.var, xlim=c(0, 1), ylim=c(0, 1),
       frame.plot=FALSE, pch=20, las=1, main=title)
  abline(0, 1, lty=3)
  abline(v=nominal.effect.size, lty=2, col=qtl.col)
  abline(h=nominal.effect.size, lty=2, col=qtl.col)
}

#' @export
sim.data.all.ve.plot <- function(sim.data,
                                 x.var=c("B.ve", "MB.ve", "DAMB.ve"),
                                 include.types=c("B.ve", "MB.ve", "DAMB.ve"),
                                 ve.col=c("#A6CEE3", "#B2DF8A", "#FB9A99"), # alts #1F78B4 #33A02C #E31A1C
                                 ve.pch=c(15, 17, 18, 19),
                                 qtl.col="red",
                                 transparency=0.5,
                                 legend.pos="topleft",
                                 title="") {
  x.var <- x.var[1]
  nominal.effect.size <- sim.data$properties$qtl.effect.size
  original.types <- colnames(sim.data$properties$var.table)[grep(x=colnames(sim.data$properties$var.table), pattern="ve")]
  
  var.table <- sim.data$properties$var.table[,unique(c(x.var, include.types))]
  
  ve.col <- c(scales::alpha(ve.col[1], transparency),
              scales::alpha(ve.col[2], transparency),
              scales::alpha(ve.col[3], transparency),
              scales::alpha(ve.col[4], transparency))
  ve.col <- ve.col[original.types %in% include.types]
  ve.pch <- ve.pch[original.types %in% include.types]
  
  x.val <- var.table[,x.var]
  keep <- !(colnames(var.table) %in% x.var)
  y.tab <- var.table[,keep]
  ve.col <- ve.col[keep]
  ve.pch <- ve.pch[keep]
  
  plot(x.val, y.tab[,ncol(y.tab)],
       xlab=x.var, ylab="Proportion of variance", xlim=c(0, 1), ylim=c(0, 1),
       frame.plot=FALSE, pch=ve.pch[ncol(y.tab)], las=1, col=ve.col[ncol(y.tab)], cex=1.5, main=title)
  if (ncol(y.tab) > 1) {
    for (i in rev(2:ncol(y.tab)-1)) {
      points(x.val, y.tab[,i], col=ve.col[i], pch=ve.pch[i], cex=1.5)
    }
  }
  abline(0, 1, lty=3)
  abline(v=nominal.effect.size, lty=2, col=qtl.col)
  abline(h=nominal.effect.size, lty=2, col=qtl.col)
  
  legend(legend.pos, legend=colnames(y.tab), col=ve.col, pch=ve.pch, cex=1.5, bty="n")
}


#' @export
sim.data.all.effect.plot <- function(sim.data,
                                     x.var=c("B.effect", "MB.effect", "DAMB.effect"),
                                     include.types=c("B.effect", "MB.effect", "DAMB.effect"),
                                     ve.col=c("#A6CEE3", "#B2DF8A", "#FB9A99"), # alts #1F78B4 #33A02C #E31A1C
                                     ve.pch=c(15, 17, 18, 19),
                                     qtl.col="red",
                                     transparency=0.5,
                                     legend.pos="topleft",
                                     title="") {
  x.var <- x.var[1]
  nominal.effect.size <- sim.data$properties$qtl.effect.size
  original.types <- colnames(sim.data$properties$var.table)[grep(x=colnames(sim.data$properties$var.table), pattern="effect")]
  
  var.table <- sim.data$properties$var.table[,unique(c(x.var, include.types))]
  
  ve.col <- c(scales::alpha(ve.col[1], transparency),
              scales::alpha(ve.col[2], transparency),
              scales::alpha(ve.col[3], transparency),
              scales::alpha(ve.col[4], transparency))
  ve.col <- ve.col[original.types %in% include.types]
  ve.pch <- ve.pch[original.types %in% include.types]
  
  x.val <- var.table[,x.var]
  keep <- !(colnames(var.table) %in% x.var)
  y.tab <- var.table[,keep]
  ve.col <- ve.col[keep]
  ve.pch <- ve.pch[keep]
  
  x.max <- max(x.val, nominal.effect.size)
  y.max <- max(y.tab, nominal.effect.size)
  
  plot(x.val, y.tab[,ncol(y.tab)],
       xlab=x.var, ylab="Proportion of variance", xlim=c(0, x.max), ylim=c(0, y.max),
       frame.plot=FALSE, pch=ve.pch[ncol(y.tab)], las=1, col=ve.col[ncol(y.tab)], cex=1.5, main=title)
  if (ncol(y.tab) > 1) {
    for (i in rev(2:ncol(y.tab)-1)) {
      points(x.val, y.tab[,i], col=ve.col[i], pch=ve.pch[i], cex=1.5)
    }
  }
  abline(0, 1, lty=3)
  abline(v=nominal.effect.size, lty=2, col=qtl.col)
  abline(h=nominal.effect.size, lty=2, col=qtl.col)
  
  legend(legend.pos, legend=colnames(y.tab), col=ve.col, pch=ve.pch, cex=1.5, bty="n")
}

#' @export power.plot
power.plot <- function(results,
                       qtl.effect.size,
                       n.alleles,
                       n.replicates=NULL,
                       col="firebrick3",
                       pch=20,
                       use.window=TRUE,
                       alpha=0.7) {
  n.replicates <- ifelse(is.null(n.replicates), results$n.replicates[1], n.replicates)
  results[,c("lower", "upper")] <- t(sapply(results[,ifelse(use.window, "power.window", "power")], binomial.prop.ci))
  plot(c(), c(), ylim=c(0,1), xlim=c(10,72), las=1, xlab="Number of strains", ylab = "Power", frame.plot=FALSE)
  data.subset <- results[results$n.replicates %in% n.replicates & results$n.alleles %in% n.alleles & results$h.qtl %in% qtl.effect.size,]
  polygon(c(data.subset$n.strains, rev(data.subset$n.strains)), 
          c(data.subset$lower,rev(data.subset$upper)), 
          col=scales::alpha(col, alpha=alpha), density=NA)
  lines(data.subset$n.strains, data.subset[,ifelse(use.window, "power.window", "power")], col="black", lwd=1.5, lend=1, type="b", cex=0.8, pch=pch)
}


#' @export add.curve.to.power.plot
add.curve.to.power.plot <- function(results,
                                    qtl.effect.size,
                                    n.alleles,
                                    n.replicates=NULL,
                                    col="skyblue",
                                    pch=1,
                                    use.window=TRUE,
                                    alpha=0.7) {
  n.replicates <- ifelse(is.null(n.replicates), results$n.replicates[1], n.replicates)
  results[,c("lower", "upper")] <- t(sapply(results[,ifelse(use.window, "power.window", "power")], binomial.prop.ci))
  data.subset <- results[results$n.replicates %in% n.replicates & results$n.alleles %in% n.alleles & results$h.qtl %in% qtl.effect.size,]
  polygon(c(data.subset$n.strains, rev(data.subset$n.strains)), 
          c(data.subset$lower,rev(data.subset$upper)), 
          col=scales::alpha(col, alpha=alpha), density=NA)
  lines(data.subset$n.strains, data.subset[,ifelse(use.window, "power.window", "power")], col="black", lwd=1.5, lend=1, type="b", cex=0.8, pch=pch)
}

#' @export fpr.plot
fpr.plot <- function(results,
                     qtl.effect.size,
                     n.alleles,
                     n.replicates=NULL,
                     col="firebrick3",
                     pch=20,
                     alpha=0.7) {
  n.replicates <- ifelse(is.null(n.replicates), results$n.replicates[1], n.replicates)
  results[,c("lower", "upper")] <- t(sapply(results$false, binomial.prop.ci))
  plot(c(), c(), ylim=c(0,1), xlim=c(10,72), las=1, xlab="Number of strains", ylab = "False positive rate", frame.plot=FALSE)
  data.subset <- results[results$n.replicates %in% n.replicates & results$n.alleles %in% n.alleles & results$h.qtl %in% qtl.effect.size,]
  polygon(c(data.subset$n.strains, rev(data.subset$n.strains)), 
          c(data.subset$lower,rev(data.subset$upper)), 
          col=scales::alpha(col, alpha=alpha), density=NA)
  lines(data.subset$n.strains, data.subset$false, col="black", lwd=1.5, lend=1, type="b", cex=0.8, pch=pch)
}


#' @export add.curve.to.fpr.plot
add.curve.to.fpr.plot <- function(results,
                                  qtl.effect.size,
                                  n.alleles,
                                  n.replicates=NULL,
                                  col="skyblue",
                                  pch=1,
                                  alpha=0.7) {
  n.replicates <- ifelse(is.null(n.replicates), results$n.replicates[1], n.replicates)
  results[,c("lower", "upper")] <- t(sapply(results$false, binomial.prop.ci))
  data.subset <- results[results$n.replicates %in% n.replicates & results$n.alleles %in% n.alleles & results$h.qtl %in% qtl.effect.size,]
  polygon(c(data.subset$n.strains, rev(data.subset$n.strains)), 
          c(data.subset$lower,rev(data.subset$upper)), 
          col=scales::alpha(col, alpha=alpha), density=NA)
  lines(data.subset$n.strains, data.subset$false, col="black", lwd=1.5, lend=1, type="b", cex=0.8, pch=pch)
}

### From miqtl originally
genome.plotter.sparcc <- function(scan.list, use.lod=FALSE, just.these.chr=NULL,
                                 scale="Mb", main.colors=c("black", "cyan", "darkgreen"),
                                 distinguish.chr.type=c("color", "box"), distinguish.box.col="gray88", 
                                 distinguish.chr.col=c("gray60", "#008080", "greenyellow"),
                                 override.col=NULL,
                                 use.legend=TRUE, 
                                 main="",
                                 my.legend.cex=0.6, my.legend.lwd=NULL, my.legend.lty=1, my.legend.pos="topright", my.legend.bty="n",
                                 y.max.manual=NULL, my.y.line=2, my.y.axis.cex=1, my.y.lab.cex=0.7,
                                 my.x.lab.cex=0.7, my.x.labels=TRUE,
                                 no.title=FALSE, override.title=NULL, my.title.line=NA, title.cex=1,
                                 hard.thresholds=NULL, thresholds.col="red", thresholds.legend=NULL,
                                 thresholds.lwd=NULL, thresholds.legend.pos="topleft", thresholds.lty=NULL,
                                 add.chr.to.label=FALSE, axis.cram=TRUE, include.x.axis.line=TRUE,
                                 mark.locus=NULL, mark.locus.col="red", which.mark=1,
                                 mark.manual=list(chr=NULL,
                                                  pos=NULL),
                                 add.polygon=FALSE, which.polygon=1,
                                 my.type="l", my.pch=20, my.cex=0.5){
  # If list has no names, use.legend is set to FALSE
  if (is.null(names(scan.list))){ use.legend=FALSE }
  if (is.null(my.legend.lwd)){ my.legend.lwd <- rep(1.5, length(scan.list)) }
  if (is.null(thresholds.lwd)){ thresholds.lwd <- rep(1, ifelse(is.null(hard.thresholds), 0, length(hard.thresholds))) }
  if (is.null(thresholds.lty)){ thresholds.lty <- rep(2, ifelse(is.null(hard.thresholds), 0, length(hard.thresholds))) }
  if (length(my.legend.lty) == 1) { my.legend.lty <- rep(my.legend.lty, length(scan.list)) }
  
  if(length(thresholds.col) < length(hard.thresholds)){ thresholds.col <- rep(thresholds.col, length(hard.thresholds)) }
  if(length(my.type) < length(scan.list)) { my.type <- rep(my.type, length(scan.list)) } 
  if(length(my.pch) < length(scan.list)) { my.pch <- rep(my.pch, length(scan.list)) } 
  
  distinguish.chr.type <- distinguish.chr.type[1]
  main.object <- scan.list[[1]]
  if(use.lod){
    outcome <- main.object$LOD
    plot.this <- this.ylab <- "LOD"
  }
  if(!use.lod){
    outcome <- -log10(main.object$p.value)
    plot.this <- "p.value"
    this.ylab <- expression("-log"[10]*"P")
  }
  if (is.null(names(outcome))) { names(outcome) <- main.object$loci } # Option for SNP scan
  
  chr <- main.object$chr
  pos <- ifelse(rep(scale=="Mb", length(outcome)), main.object$pos$Mb, main.object$pos$cM)
  
  if(!is.null(just.these.chr)){
    keep.chr <- chr %in% just.these.chr
    chr <- chr[keep.chr]
    outcome <- outcome[keep.chr]
    pos <- pos[keep.chr]
  }
  
  has.X <- FALSE
  if(any(chr=="X")){
    has.X <- TRUE
    chr[chr=="X"] <- max(as.numeric(unique(chr[chr != "X"]))) + 1
  }
  
  pre.chr <- as.factor(as.numeric(chr))
  order.i <- order(pre.chr, pos)
  
  outcome <- outcome[order.i]
  pre.chr <- pre.chr[order.i]
  pos <- pos[order.i]
  
  chr.types <- levels(pre.chr)
  
  ##################################################################
  ######################### Build scaffold ######################### 
  ##################################################################
  shift.vector <- build.position.scaffold(scan.list=scan.list, scale=scale)
  if (!is.null(just.these.chr)) { shift.vector <- shift.vector[just.these.chr]}
  
  updated.pos <- rep(NA, length(outcome))
  names(updated.pos) <- names(outcome)
  updated.pos[pre.chr==chr.types[1]] <- pos[pre.chr==chr.types[1]]
  
  x.max <- sum(shift.vector)
  ##################################################################
  ##################################################################
  
  # Finding max y of plot window
  y.max <- ceiling(max(outcome, hard.thresholds)) 
  if(length(scan.list) > 1){
    for(i in 2:length(scan.list)){
      if(use.lod){
        y.max <- ceiling(max(y.max, unlist(scan.list[[i]][plot.this])))
      }
      if(!use.lod){
        y.max <- ceiling(max(y.max, -log10(unlist(scan.list[[i]][plot.this]))))
      }
    }
  }
  if(!is.null(y.max.manual)){
    y.max <- y.max.manual
  }
  
  ### Fixef or ranef
  if(length(scan.list) == 1 & !is.null(scan.list[[1]]$locus.effect.type)){
    locus.effect.type <- ifelse(scan.list[[1]]$locus.effect.type == "fixed", "fixef", "ranef")
    locus.term <- paste("locus", locus.effect.type, sep=".")
  }
  else{
    locus.term <- "locus"
  }
  
  if(no.title){ this.title <- NULL }
  else if(!is.null(override.title)){ this.title <- override.title }
  else{
    ### Handling the annoying differences between lmer and lm objects
    if(is.null(scan.list[[1]]$fit0)){
      this.title <- c(main, 
                      paste0(scan.list[[1]]$formula, " + ", locus.term, " (", scan.list[[1]]$model.type, ")"),
                      paste("n =", round(scan.list[[1]]$n, 2)))
    }
    else{
      if(class(scan.list[[1]]$fit0) != "lmerMod"){
        this.title <- c(main, 
                        paste0(scan.list[[1]]$formula, " + ", locus.term, " (", scan.list[[1]]$model.type, ")"),
                        paste("n =", round(ifelse(is.null(scan.list[[1]]$fit0$weights), 
                                                  length(scan.list[[1]]$fit0$y),
                                                  sum(scan.list[[1]]$fit0$weights)), 2)))
      }
      else{
        this.title <- c(main, 
                        paste0(scan.list[[1]]$formula, " + ", locus.term, " (", scan.list[[1]]$model.type, ")"),
                        paste("n =", round(sum(scan.list[[1]]$fit0@resp$weights), 2)))
      }
    }
  }
  
  if (distinguish.chr.type == "box") { 
    this.col <- rep(main.colors[1], length(outcome)) 
  }
  else if (distinguish.chr.type == "color") { 
    this.col <- c(main.colors[1], distinguish.chr.col[1])[(as.numeric(as.character(pre.chr)) %% 2 == 0) + 1]
  }
  if (!is.null(override.col)) {
    this.col <- override.col[(as.numeric(as.character(pre.chr)))]
  }
  
  plot(pos[pre.chr==chr.types[1]], outcome[pre.chr==chr.types[1]], 
       xlim=c(0, x.max), 
       ylim=c(-0.1, y.max), 
       xaxt="n", yaxt="n", ylab="", xlab="", main=NA,
       frame.plot=FALSE, type=my.type[1], pch=my.pch[1], cex=my.cex, lwd=my.legend.lwd[1], lty=my.legend.lty[1], col=this.col[1])
  title(main=this.title, line=my.title.line, cex.main=title.cex)
  axis(side=2, at=0:y.max, las=2, cex.axis=my.y.axis.cex)
  mtext(text=this.ylab, side=2, line=my.y.line, cex=my.y.lab.cex)
  
  if (1 %in% which.polygon & add.polygon) {
    polygon.x.and.y <- expand.for.polygon(x=pos[pre.chr==chr.types[1]], y=outcome[pre.chr==chr.types[1]])
    polygon(polygon.x.and.y$x, 
            polygon.x.and.y$y, 
            col=main.colors[1],
            border=NA)
  }
  
  label.spots <- shift.vector[1]/2
  x.tick.spots <- c(0, shift.vector[1])
  shift <- shift.vector[1]
  
  if(length(chr.types) > 1){
    for(i in 2:length(chr.types)){
      this.pos <- pos[pre.chr==chr.types[i]] + shift
      updated.pos[pre.chr==chr.types[i]] <- pos[pre.chr==chr.types[i]] + shift
      if(distinguish.chr.type == "box"){
        if(i %% 2 == 0){
          polygon(x=c(shift, max(this.pos, na.rm=TRUE), max(this.pos, na.rm=TRUE), shift), 
                  y=c(y.max, y.max, 0, 0), border=NA, col=distinguish.box.col)
          
        }
      }
      label.spots <- c(label.spots, shift + shift.vector[i]/2)
      x.tick.spots <- c(x.tick.spots, shift + shift.vector[i])
      points(this.pos, outcome[pre.chr==chr.types[i]], type=my.type[1], pch=my.pch[1],
             lty=my.legend.lty[1], cex=my.cex, lwd=my.legend.lwd[1], col=this.col[pre.chr==chr.types[i]])
      if (1 %in% which.polygon & add.polygon) {
        polygon.x.and.y <- expand.for.polygon(x=this.pos, y=outcome[pre.chr==chr.types[i]])
        polygon(polygon.x.and.y$x, 
                polygon.x.and.y$y, 
                col=this.col[pre.chr==chr.types[i]][1],
                border=NA)
      }
      shift <- shift + shift.vector[i]
    }
  }
  
  # Plot other method's statistics
  if(length(scan.list) > 1){
    for(i in 2:length(scan.list)){
      this.scan <- scan.list[[i]]
      if(use.lod){
        compar.outcome <- this.scan$LOD
      }
      if(!use.lod){
        compare.outcome <- -log10(this.scan$p.value)
      }
      pos <- ifelse(rep(scale=="Mb", length(compare.outcome)), this.scan$pos$Mb, this.scan$pos$cM)
      
      ## Resetting for new scan objects
      chr <- this.scan$chr
      if(!is.null(just.these.chr)){
        keep.chr <- chr %in% just.these.chr
        chr <- chr[keep.chr]
        compare.outcome <- compare.outcome[keep.chr]
        pos <- pos[keep.chr]
      }
      
      has.X <- FALSE
      if(any(chr=="X")){
        has.X <- TRUE
        chr[chr=="X"] <- max(as.numeric(unique(chr[chr != "X"]))) + 1
      }
      
      pre.chr <- as.factor(as.numeric(chr))
      order.i <- order(pre.chr, pos)
      
      compare.outcome <- compare.outcome[order.i]
      pre.chr <- pre.chr[order.i]
      pos <- pos[order.i]
      
      chr.types <- levels(pre.chr)
      
      ## Setting up colors for additional scans
      if(distinguish.chr.type == "box"){ 
        this.col <- rep(main.colors[i], length(compare.outcome)) 
      }
      else if(distinguish.chr.type == "color"){ 
        this.col <- c(main.colors[i], distinguish.chr.col[i])[(as.numeric(as.character(pre.chr)) %% 2 == 0) + 1]
      }
      if (!is.null(override.col)) {
        this.col <- override.col[(as.numeric(as.character(pre.chr)))]
      }
      
      compare.shift <- shift.vector[1]
      points(pos[pre.chr==chr.types[1]], compare.outcome[pre.chr==chr.types[1]], type=my.type[i], cex=my.cex, pch=my.pch[i],
             col=this.col[pre.chr==chr.types[1]], lwd=my.legend.lwd[i], lty=my.legend.lty[i])
      
      if (i %in% which.polygon & add.polygon) {
        polygon.x.and.y <- expand.for.polygon(x=pos[pre.chr==chr.types[1]], y=compare.outcome[pre.chr==chr.types[1]])
        polygon(polygon.x.and.y$x, 
                polygon.x.and.y$y, 
                col=this.col[pre.chr==chr.types[1]][1],
                border=NA)
      }
      
      ## For later plotting, like mark.locus
      if (i == which.mark) {
        updated.pos <- rep(NA, length(compare.outcome))
        names(updated.pos) <- names(compare.outcome)
        updated.pos[pre.chr==chr.types[1]] <- pos[pre.chr==chr.types[1]]
      }
      
      if (length(chr.types) > 1) {
        for (j in 2:length(chr.types)) {
          points(pos[pre.chr==chr.types[j]] + compare.shift, compare.outcome[pre.chr==chr.types[j]], 
                 type=my.type[i], cex=my.cex, pch=my.pch[i],
                 col=this.col[pre.chr==chr.types[j]], 
                 lwd=my.legend.lwd[i],
                 lty=my.legend.lty[i])
          if (i %in% which.polygon & add.polygon) {
            polygon.x.and.y <- expand.for.polygon(x=pos[pre.chr==chr.types[j]] + compare.shift, y=compare.outcome[pre.chr==chr.types[j]])
            polygon(polygon.x.and.y$x, 
                    polygon.x.and.y$y, 
                    col=this.col[pre.chr==chr.types[j]][1],
                    border=NA)
          }
          if (i == which.mark) {
            updated.pos[pre.chr==chr.types[j]] <- pos[pre.chr==chr.types[j]] + compare.shift
          }
          compare.shift <- compare.shift + shift.vector[j]
        }
      }
    }
  }
  if (has.X) {
    axis.label <- c(chr.types[-length(chr.types)], "X")
  }
  else {
    axis.label <- chr.types
  }
  
  if (include.x.axis.line) {
    axis(side=1, tick=TRUE, line=NA, at=x.tick.spots, 
         labels=NA, xpd=TRUE)
  }
  
  if (add.chr.to.label) {
    axis.label <- paste("Chr", axis.label)
  }
  else {
    axis.label <- c("Chr", axis.label)
    label.spots <- c(-0.04*x.max, label.spots)
  }
  
  if (axis.cram) {
    odd.axis.label <- axis.label[(1:length(axis.label) %% 2) == 1]
    odd.label.spots <- label.spots[(1:length(label.spots) %% 2) == 1]
    
    even.axis.label <- axis.label[(1:length(axis.label) %% 2) == 0]
    even.label.spots <- label.spots[(1:length(label.spots) %% 2) == 0]
    
    if (!my.x.labels) { even.axis.label <- FALSE; odd.axis.label <- FALSE }
    
    axis(side=1, tick=FALSE, line=NA, at=odd.label.spots, labels=odd.axis.label, cex.axis=my.x.lab.cex, padj=-1.5, xpd=TRUE)
    axis(side=1, tick=FALSE, line=NA, at=even.label.spots, labels=even.axis.label, cex.axis=my.x.lab.cex, padj=-1.5, xpd=TRUE)
  }
  else {
    if (!my.x.labels) { axis.label <- FALSE }
    axis(side=1, tick=FALSE, line=NA, at=label.spots, labels=axis.label, cex.axis=my.x.lab.cex, padj=-1.5, xpd=TRUE)
  }
  if (!is.null(mark.locus)) {
    rug(x=updated.pos[which(names(updated.pos) %in% mark.locus)], lwd=10, col=mark.locus.col, ticksize=0.05)
  }
  if (!is.null(mark.manual$chr)) {
    rug(x=calc.manual.mark.locus(shift.vector=shift.vector, mark.manual=mark.manual), lwd=10, col=mark.locus.col, ticksize=0.05)
  }
  if (use.legend) {
    if (add.polygon) {
      these.lty <- my.legend.lty
      these.lty[my.type != "l"] <- 0
      these.pch <- my.pch
      these.pch[my.type != "p"] <- NA
      these.lwd <- my.legend.lwd
      these.fill.col <- main.colors
      these.fill.border <- main.colors
      these.lty[which.polygon] <- these.lwd[which.polygon] <- NA
      these.fill.col[!(1:length(scan.list) %in% which.polygon)] <- these.fill.border[!(1:length(scan.list) %in% which.polygon)] <-  NA
      this.x.intersp=sapply(1:length(these.fill.col), function(x) ifelse(is.na(these.fill.col[x]) & my.type[x] == "l", 2, 0.5))
      legend(my.legend.pos, 
             legend=names(scan.list), 
             lty=these.lty, 
             pch=these.pch,
             lwd=these.lwd,
             fill=these.fill.col,
             border=these.fill.border,
             x.intersp=this.x.intersp,
             col=main.colors[1:length(scan.list)], bty=my.legend.bty, cex=my.legend.cex)
    }
    else {
      these.lty <- my.legend.lty
      these.lty[my.type != "l"] <- 0
      these.pch <- my.pch
      these.pch[my.type != "p"] <- NA
      legend(my.legend.pos, 
             legend=names(scan.list), 
             lty=these.lty, 
             pch=these.pch,
             lwd=my.legend.lwd, 
             col=main.colors[1:length(scan.list)], bty=my.legend.bty, cex=my.legend.cex)
    }
  }
  if (!is.null(hard.thresholds)) {
    for (i in 1:length(hard.thresholds)) {
      abline(h=hard.thresholds[i], col=thresholds.col[i], lty=thresholds.lty[i], lwd=thresholds.lwd[i])
    }
  }
  if (!is.null(thresholds.legend)) {
    legend(thresholds.legend.pos, legend=thresholds.legend, col=thresholds.col, lty=thresholds.lty,
           lwd=thresholds.lwd, bty=my.legend.bty, cex=my.legend.cex)
  }
}

build.position.scaffold <- function(scan.list, scale) {
  for (i in 1:length(scan.list)) {
    chr <- scan.list[[i]]$chr
    pos <- scan.list[[i]]$pos[[scale]]
    ## Handling X
    has.X <- FALSE
    if(any(chr=="X")){
      has.X <- TRUE
      chr[chr=="X"] <- max(as.numeric(unique(chr[chr != "X"]))) + 1
    }
    
    pre.chr <- as.factor(as.numeric(chr))
    max.pos <- tapply(pos, pre.chr, function(x) max(x, na.rm=TRUE))
    if (i == 1) {
      total.max.pos <- rep(0, length(max.pos))
    }
    total.max.pos <- sapply(1:length(max.pos), function(x) max(max.pos[x], total.max.pos[x]))
  }
  names(total.max.pos) <- names(max.pos)
  if (has.X) { names(total.max.pos)[length(total.max.pos)] <- "X" }
  return(total.max.pos)
}

expand.for.polygon <- function(x, y){
  if(any(is.na(x)) | any(is.na(y))){
    remove.na.x <- which(is.na(x))
    remove.na.y <- which(is.na(y))
    remove.na <- sort(c(remove.na.x, remove.na.y))
    x <- x[-remove.na]
    y <- y[-remove.na]
  }
  return(list(x=c(x[1], x, x[length(x)]),
              y=c(0, y, 0)))
}

calc.manual.mark.locus <- function(shift.vector, mark.manual) {
  here <- which(names(shift.vector) == mark.manual$chr)
  if (here == 1) {
    new.pos <- mark.manual$pos
  }
  else {
    new.pos <- sum(shift.vector[1:(here - 1)]) + mark.manual$pos
  }
  return(new.pos)
}
```
