## Supplementary material for "Determinants of QTL mapping power in the realized Collaborative Cross": SPARCC R package used for all analyses: CC_sim.R

```
#' Runs the genome scans of the simulated data output by sim.CC.data()
#'
#' This function takes the output from sim.CC.data() and performs the genome scans. Internally it runs
#' the require QR decompositions, which it can save for later if specified.
#'
#' @param sim.data Output simulated data from sim.CC.data().
#' @param scan.index DEFAULT: NULL. If NULL, it performs scans for all simulated data sets stored in sim.data.
#' If given a vector of integers, representing the simulation index, it will only run scans for those phenotypes.
#' @param chr DEFAULT: "all". Specifies which chromosomes to scan.
#' @param just.these.loci DEFAULT: NULL. If NULL, all loci in genome cache are scanned.
#' @param use.progress.bar DEFAULT: FALSE. Specifies whether to use a progress bar for qr decompositions 
#' and genome scans. 
#' @param print.scans.progress DEFAULT: FALSE. Specifies whether to output a message that indicates the number of
#' scans completed.
#' @param all.sim.qr DEFAULT: NULL. If NULL, necessary qr decompositions are performed, which will slow down
#' the function significantly. If non-NULL, it expects either output from extract.qr for a single sample
#' of CC lines, or a list of output for multiple samples.
#' @param return.all.sim.qr DEFAULT: TRUE. Saves the qr decompositions for later use, though will make output
#' substantially larger. Could then be used for later scans with the same set of CC lines.
#' @export
#' @examples run.sim.scans()
run.sim.scans <- function(sim.data,
                          scan.index=NULL, 
                          chr="all", 
                          just.these.loci=NULL,
                          use.progress.bar=FALSE,
                          print.scans.progress=FALSE,
                          all.sim.qr=NULL,
                          return.all.sim.qr=TRUE,
                          ...){
  
  if(is.null(scan.index)){ 
    scan.index <- 1:sum(grepl("sim.y", colnames(sim.data$data)))
  }
  h <- DiploprobReader$new(sim.data$genomecache)
  loci <- h$getLoci()
  loci.chr <- h$getChromOfLocus(loci)
  if(chr[1] != "all"){
    loci <- loci[loci.chr %in% chr]
  }
  if(!is.null(just.these.loci)){
    loci <- loci[loci %in% just.these.loci]
    loci.chr <- loci.chr[loci %in% just.these.loci]
  }
  
  full.p <- matrix(NA, nrow=length(scan.index), ncol=length(loci))
  colnames(full.p) <- loci
  these.pos <- list(Mb=h$getLocusStart(loci, scale="Mb"),
                    cM=h$getLocusStart(loci, scale="cM"))
  
  if(!sim.data$properties$vary.lines){
    if(is.null(all.sim.qr)){
      this.qr <- extract.qr(genomecache=sim.data$genomecache, model="additive",
                            formula=~1, data=sim.data$data, 
                            id="SUBJECT.NAME.1",
                            just.these.loci=just.these.loci, chr=chr,
                            use.progress.bar=use.progress.bar)
    }
    else{
      this.qr <- all.sim.qr
    }
    this.id <- "SUBJECT.NAME.1"
  }
  compute.qr.boolean <- is.null(all.sim.qr)
  if (return.all.sim.qr & is.null(all.sim.qr)){ all.sim.qr <- list() }
  for (i in 1:length(scan.index)) {
    if (sim.data$properties$vary.lines) {
      if (compute.qr.boolean) {
        this.qr <- extract.qr(genomecache=sim.data$genomecache, model="additive",
                              formula=~1, data=sim.data$data,
                              id=paste0("SUBJECT.NAME.", i),
                              just.these.loci=just.these.loci, chr=chr,
                              use.progress.bar=use.progress.bar)
      }
      else{
        this.qr <- pull.qr.from.compact(compact.qr.list=all.sim.qr, 
                                        qr.index=scan.index[i])
      }
      this.id <- paste0("SUBJECT.NAME.", i)
    }
    this.scan <- scan.qr(qr.object=this.qr, data=sim.data$data, 
                         phenotype=paste0("sim.y.", i), id=this.id, chr=chr, 
                         return.allele.effects=FALSE, use.progress.bar=use.progress.bar,
                         ...)
    full.p[i,] <- this.scan$p.value
    if (return.all.sim.qr & sim.data$properties$vary.lines) { all.sim.qr[[i]] <- this.qr }
    else if (return.all.sim.qr & !sim.data$properties$vary.lines) { all.sim.qr <- this.qr }
    if(print.scans.progress){
      cat("\n", "Simulation scan: index", scan.index[i], "complete ---------- final index of this run:", scan.index[length(scan.index)], "\n")
    }
  }
  output <- list(LOD=NULL,
                 p.value=full.p,
                 chr=loci.chr, 
                 pos=these.pos,
                 locus=sim.data$locus,
                 locs.pos=sim.data$locus.pos,
                 properties=sim.data$properties)
  if(return.all.sim.qr){
    output$all.sim.qr <- all.sim.qr
  }
  return(output)
}

#' Simulate Collaborative Cross (CC) phenotype data from the realized CC lines founder haplotype probabilities stored in a genome cache directory
#'
#' This function takes various input parameters to simulate CC data to be used in power calculations or as input data for 
#' other tools that analyze CC data.
#'
#' @param genomecache The path to the genome cache directory. The genome cache is a particularly structured
#' directory that stores the haplotype probabilities/dosages at each locus. It has an additive model
#' subdirectory and a full model subdirectory. Each contains subdirectories for each chromosome, which then
#' store .RData files for the probabilities/dosages of each locus.
#' @param CC.lines DEFAULT: NULL. If NULL is specified, sim.CC.data() will randomly draw samples of the 
#' available CC lines of the size specified in num.lines.
#' @param num.lines DEFAULT: NULL. If NULL, sim.CC.data() expects that CC.lines is non-NULL. If CC.lines
#' is NULL, num.lines determines the number of CC lines that are sampled.
#' @param vary.lines DEFAULT: TRUE. If CC.lines is NULL and vary.lines is TRUE, then sim.CC.data() will
#' sample multiple sets of CC lines of the size specified in num.lines. If CC.lines is NULL and vary.lines is
#' FALSE, then just one set of CC lines will be sampled.
#' @param locus DEFAULT: NULL. If NULL is specified, sim.CC.data() will randomly draw a locus stored in
#' the genome cache. 
#' @param vary.locus DEFAULT: TRUE. If locus is NULL and vary.locus is TRUE, then sim.CC.data() will sample
#' as many loci as specified in num.sim. If locus is NULL and vary.locus is FALSE, then sim.CC.data() will
#' only sample one locus.
#' @param num.replicates The number of replicates of each CC line that will be simulated. Mapping for power calculations
#' will use strain means. Currently requires that all lines have the same number of replicates.
#' @param num.sim The number of phenotypes to be simulated for a given parameter setting.
#' @param M.ID DEFAULT: NULL. M is a matrix that maps from counts of founder haplotypes to counts of functional 
#' alleles. Mapping will be based on haplotype association, but potentially there are two to the number of
#' founders alleles at the QTL. M.ID is a string that codifies this mapping. One potential balanced two allele
#' M.ID would be "c(0,0,0,0,1,1,1,1)". With 8 functional alleles, on per founder, the only M.ID is "c(0,1,2,3,4,5,6,7)".
#' If M.ID is NULL, M.ID will be sampled.
#' @param sample.as.method DEFAULT: "uniform". The procedure used for sampling the allelic series. If
#' every strain has its own allele, this option does not matter. Alternatively, a Chinese restaurant process ("crp") can be used,
#' which is possibly more biologically accurate, and will favor allelic series that are less balanced (1 vs 7).
#' @param num.alleles DEFAULT: 8. The number of functional alleles. Must be less than or equal to the number of 
#' founders.
#' @param num.founders DEFAULT: 8. The number of founders, which must correspond to the genome cache. The CC has
#' eight.
#' @param qtl.effect.size The size of the simulated QTL effect. The scale of the input is in proportion of
#' the phenotypic variance due to the QTL, thus should be greater than or equal to zero, and less than one.
#' @param beta DEFAULT: NULL. Allows for the manual specification of QTL effect. Is expected to be a vector
#' the length of the number of alleles. It will be scaled based on qtl.effect.size.
#' @param strain.effect.size DEFAULT: 0. The size of the simulated strain effect, which represents something akin to a polygenic 
#' effect. Other variants specific to CC lines will result in overall strain-specific effects. The scale of the input 
#' is in proportion of the phenotypic variance due to the strain, thus should be greater than or equal to zero, and less than one.
#' @param impute DEFAULT: TRUE. If TRUE, the QTL portion of the design matrix in the simulation is a realized sampling
#' of haplotypes from the probabilities. If FALSE, the simulations are based on the probabilities, which is flawed in
#' terms of biological reality.
#' @param scale.qtl.mode DEFAULT: "B". Specifies how the QTL effect is scaled. If "B", then the variance of the qtl effect
#' vector beta is scaled to the effect size specified in qtl.effect.size, which would be the variance explained in a population
#' perfectly balanced in terms of the functional alleles. If "MB", the the variance of M %*% beta is scaled, which would be
#' the variance explained in a population balanced in terms of founder strains with the allelic series that is specified in M. 
#' If "DAMB", the variance of D %*% A %*% M %*% beta is scaled, which would be the variance explained in a population of a 
#' specific set of CC strains (specified in D).
#' @param return.value DEFAULT: "raw". If "raw", residuals are not taken. If "fixef.resid", then the data
#' are residuals after regressing phenotype on strain. If "ranef.resid", then the data have had the strain BLUP
#' effect subtracted.
#' @param return.means DEFAULT: TRUE. If TRUE, strain means are returned. If FALSE, the full data
#' with replicate observations of strains are returned.
#' @export
#' @examples sim.CC.data()
sim.CC.data <- function(genomecache, 
                        CC.lines=NULL, 
                        num.lines=NULL, 
                        vary.lines=TRUE,
                        locus=NULL, 
                        vary.locus=TRUE,
                        num.replicates, 
                        num.sim,
                        M.ID=NULL,
                        sample.as.method=c("uniform", "crp"),
                        num.alleles=8, 
                        num.founders=8,
                        qtl.effect.size,
                        beta=NULL,
                        strain.effect.size=0,
                        impute=TRUE,
                        scale.qtl.mode=c("B", "MB", "DAMB", "none"),
                        return.value=c("raw", "fixef.resid", "ranef.resid"),
                        return.means=TRUE){
  
  h <- DiploprobReader$new(genomecache)
  scale.qtl.mode <- scale.qtl.mode[1]
  return.value <- return.value[1]
  sample.as.method <- sample.as.method[1]
  
  original.effects <- list(qtl.effect.size=qtl.effect.size,
                           strain.effect.size=strain.effect.size)
  
  ############ Scaling effects
  noise.effect.size <- (1 - qtl.effect.size - strain.effect.size)
  reduced.noise.effect.size <- noise.effect.size/num.replicates
  s <- 1/min(c(qtl.effect.size, strain.effect.size, reduced.noise.effect.size)[c(qtl.effect.size, strain.effect.size, reduced.noise.effect.size) != 0])
  qtl.effect.size <- s*qtl.effect.size
  strain.effect.size <- s*strain.effect.size
  noise.effect.size <- s*noise.effect.size
  
  ## Sampling lines
  if (is.null(CC.lines)) {
    CC.lines <- sapply(1:ifelse(vary.lines, num.sim, 1), function(i) sample(x=h$getSubjects(), size=num.lines, replace=FALSE))
    if (vary.lines) {
      cc.index <- 1:num.sim
    }
    else {
      cc.index <- rep(1, num.sim)
    }
  }
  else{
    if(is.vector(CC.lines)){
      CC.lines <- matrix(CC.lines, ncol=1)
      vary.lines <- FALSE
      num.lines <- length(CC.lines)
      cc.index <- rep(1, num.sim)
    }
    else{
      vary.lines <- TRUE
      num.lines <- nrow(CC.lines)
      num.sim <- ncol(CC.lines)
      cc.index <- 1:num.sim
    }
  }
  num.ind <- ifelse(return.means, num.lines, num.lines*num.replicates)
  
  ## Setting number of alleles to match pre-specified QTL effect - convenience
  if(!is.null(beta)){
    num.alleles <- length(beta)
  }
  
  ## Sampling loci
  if(is.null(locus)){
    locus <- sample(h$getLoci(), size=ifelse(vary.locus, num.sim, 1), replace=TRUE)
    if(vary.locus){
      locus.index <- 1:num.sim
    }
    else{
      locus.index <- rep(1, num.sim)
    }
  }
  else{
    vary.locus <- FALSE
    locus.index <- rep(1, num.sim)
  }
  
  M <- NULL
  if(!is.null(M.ID)){
    M <- model.matrix.from.ID(M.ID)
    num.alleles <- length(unique(unlist(strsplit(x=M.ID, split=","))))
  }
  B.var.table <- MB.var.table <- DAMB.var.table <- matrix(NA, nrow=num.sim, ncol=6)
  sim.matrix <- matrix(NA, nrow=num.ind, ncol=num.sim)
  id.matrix <- matrix(NA, nrow=num.ind, ncol=ifelse(vary.lines, num.sim, 1))
  for(i in 1:num.sim){
    this.sim <- simulate.CC.qtl(CC.lines=CC.lines[,cc.index[i]], 
                                num.replicates=num.replicates,
                                M=M,
                                sample.as.method=sample.as.method,
                                num.alleles=num.alleles, 
                                num.founders=num.founders,
                                qtl.effect.size=qtl.effect.size, 
                                strain.effect.size=strain.effect.size,
                                noise.effect.size=noise.effect.size,
                                impute=impute,
                                scale.qtl.mode=scale.qtl.mode,
                                locus.matrix=h$getLocusMatrix(locus=locus[locus.index[i]], model="full"),
                                return.value=return.value,
                                return.means=return.means,
                                num.sim=1)
    these.var.tables <- this.sim$properties$var.tables
    this.sim <- this.sim$data
    sim.matrix[,i] <- this.sim[,1]
    B.var.table[i,] <- these.var.tables$B[1,]
    MB.var.table[i,] <- these.var.tables$MB[1,]
    DAMB.var.table[i,] <- these.var.tables$DAMB[1,]
    if(vary.lines | i == 1){
      id.matrix[,i] <- as.character(this.sim[,2])
    }
  }
  colnames(sim.matrix) <- paste0("sim.y.", 1:num.sim)
  colnames(id.matrix) <- paste0("SUBJECT.NAME.", 1:ncol(id.matrix))
  colnames(B.var.table) <- colnames(MB.var.table) <- colnames(DAMB.var.table) <- colnames(these.var.tables$B)
  
  outcome <- data.frame(sim.matrix, id.matrix)
  return(list(data=outcome,
              locus=locus,
              locus.pos=list(Mb=h$getLocusStart(locus, scale="Mb"),
                             cM=h$getLocusStart(locus, scale="cM")),
              genomecache=genomecache,
              properties=list(num.alleles=num.alleles,
                              sample.as.method=sample.as.method,
                              num.replicates=num.replicates,
                              num.founders=num.founders,
                              qtl.effect.size=original.effects$qtl.effect.size, 
                              strain.effect.size=original.effects$strain.effect.size,
                              num.lines=num.lines,
                              impute=impute,
                              scale.qtl.mode=scale.qtl.mode,
                              M.ID=M.ID,
                              vary.lines=vary.lines,
                              vary.locus=vary.locus,
                              return.value=return.value,
                              return.means=return.means,
                              var.tables=list(B=B.var.table,
                                              MB=MB.var.table,
                                              DAMB=DAMB.var.table))))
}

simulate.CC.qtl <- function(CC.lines, 
                            num.replicates,
                            M=NULL,
                            sample.as.method=c("uniform", "crp"),
                            qtl.effect.size, 
                            beta=NULL,
                            strain.effect.size,
                            noise.effect.size,
                            num.alleles=8, 
                            num.founders=8,
                            locus.matrix, 
                            num.sim,
                            impute=TRUE, 
                            scale.qtl.mode=c("B", "MB", "DAMB", "none"),
                            return.value=c("raw", "fixef.resid", "ranef.resid"),
                            return.means=TRUE,
                            ...){
  scale.qtl.mode <- scale.qtl.mode[1]
  return.value <- return.value[1]
  sample.as.method <- sample.as.method[1]

  this.locus.matrix <- locus.matrix[CC.lines,]
  ## Imputation for simulation
  if(impute){
    this.locus.matrix <- t(apply(this.locus.matrix, 1, function(x) rmultinom(1, 1, x)))
    rownames(this.locus.matrix) <- CC.lines
  }
  D <- this.locus.matrix ## Saved for variance calculations
  ZD <- D[rep(1:length(CC.lines), each=num.replicates),]
  
  full.to.add.matrix <- t(straineff.mapping.matrix(M=num.founders))
  
  ## Strain
  Z <- incidence.matrix(factor(CC.lines, levels=CC.lines))
  Z <- Z[rep(1:nrow(Z), each=num.replicates),]
  if (strain.effect.size != 0) {
    strain.effect <- rnorm(n=length(CC.lines))
    strain.effect <- (strain.effect - mean(strain.effect))/sqrt(non.sample.var(strain.effect))
    strain.effect <- strain.effect * sqrt(strain.effect.size)

    strain.predictor <- Z %*% matrix(strain.effect, ncol=1)
  }
  else{ strain.predictor <- rep(0, nrow(Z)) }
  
  ## QTL
  ZDA <- tcrossprod(ZD, full.to.add.matrix)
  if (qtl.effect.size != 0) {
    QTL.effect <- simulate.QTL.model.and.effects(num.alleles=num.alleles, 
                                                 num.founders=num.founders, 
                                                 M=M,
                                                 sample.as.method=sample.as.method,
                                                 beta=beta,
                                                 ...)
  }
  else {
    QTL.effect <- list(M=diag(8),
                       beta=rep(0, 8))
  }
  M <- QTL.effect$M
  raw.beta <- QTL.effect$beta 
  if (scale.qtl.mode != "none" & qtl.effect.size != 0) {
    beta <- (raw.beta - mean(raw.beta))/sqrt(non.sample.var(raw.beta))
  }
  else { beta <- raw.beta }
  
  ### B
  B.var.effects <- calc.qtl.effect(beta = 0.5*beta*sqrt(qtl.effect.size),
                                   M = M,
                                   A = t(full.to.add.matrix),
                                   D = D,
                                   Z = Z,
                                   strain.effect.size = strain.effect.size,
                                   noise.effect.size = noise.effect.size)
  if (scale.qtl.mode == "B") {
    QTL.predictor <- B.var.effects$QTL.predictor
  }
  
  ### MB
  var.ratio <- ifelse(qtl.effect.size != 0,
                      c(non.sample.var(2*M %*% beta)/non.sample.var(2*beta)), 
                      1)
  MB.var.effects <- calc.qtl.effect(beta = 0.5*beta*sqrt(qtl.effect.size)*sqrt(1/var.ratio),
                                    M = M,
                                    A = t(full.to.add.matrix),
                                    D = D,
                                    Z = Z,
                                    strain.effect.size = strain.effect.size,
                                    noise.effect.size = noise.effect.size)
  if (scale.qtl.mode == "MB") {
    QTL.predictor <- MB.var.effects$QTL.predictor
  }
  
  ### DAMB
  var.ratio <- ifelse(qtl.effect.size != 0,
                      c(non.sample.var(D %*% t(full.to.add.matrix) %*% M %*% beta)/non.sample.var(2*beta)),
                      1)
  if (var.ratio != 0) { # Case when more than one allele is observed
    DAMB.beta <- 0.5*beta*sqrt(qtl.effect.size)*sqrt(1/var.ratio)
  }
  else { # Case when only one allele is observed, should result in a null scan
    DAMB.beta <- 0.5*beta*sqrt(qtl.effect.size)
  }
  DAMB.var.effects <- calc.qtl.effect(beta = DAMB.beta,
                                      M = M,
                                      A = t(full.to.add.matrix),
                                      D = D,
                                      Z = Z,
                                      strain.effect.size = strain.effect.size,
                                      noise.effect.size = noise.effect.size)
  if (scale.qtl.mode == "DAMB") {
    QTL.predictor <- DAMB.var.effects$QTL.predictor
  }

  B.var.table <- MB.var.table <- DAMB.var.table <- matrix(NA, nrow=num.sim, ncol=6)
  sim.data <- matrix(NA, nrow=nrow(Z), ncol=num.sim)
  for (i in 1:num.sim) {
    scaled.resid <- calc.scaled.residual(noise.effect.size=noise.effect.size,
                                         n=nrow(Z))
    sim.data[,i] <- QTL.predictor + strain.predictor + scaled.resid
    B.var.table[i,] <- B.var.effects$summaries
    MB.var.table[i,] <- MB.var.effects$summaries
    DAMB.var.table[i,] <- DAMB.var.effects$summaries
  }
  
  colnames(sim.data) <- paste0("sim.y.", 1:ncol(sim.data))
  sim.data <- data.frame(sim.data, "SUBJECT.NAME"=rep(CC.lines, each=num.replicates))
  colnames(B.var.table) <- colnames(MB.var.table) <- colnames(DAMB.var.table) <- c("B.effect", "MB.effect", "DAMB.effect",
                                                                                   "B.ve", "MB.ve", "DAMB.ve")
  
  ## Taking residuals
  if (return.value == "fixef.resid") {
    sim.data$sim.y.1 <- take.fixef.residuals(data=sim.data, 
                                             locus.matrix=this.locus.matrix, 
                                             strain.matrix=this.strain.matrix)
  }
  else if (return.value == "ranef.resid") {
    sim.data$sim.y.1 <- take.ranef.residuals(data=sim.data,
                                             locus.matrix=this.locus.matrix)
  }
  if (return.means) {
    sim.data <- apply(sim.data[,-ncol(sim.data), drop=FALSE], 2, function(x) tapply(x, INDEX=as.factor(rep(CC.lines, each=num.replicates)), FUN=mean))
    colnames(sim.data) <- paste0("sim.y.", 1:ncol(sim.data))
    sim.data <- data.frame(sim.data, "SUBJECT.NAME"=rownames(sim.data))
  }
  return(list(data=sim.data,
              properties=list(qtl.effect.size=qtl.effect.size,
                              strain.effect.size=strain.effect.size,
                              num.alleles=num.alleles,
                              sample.as.method=sample.as.method,
                              num.replicates=num.replicates,
                              num.lines=length(CC.lines),
                              impute=impute,
                              return.value=return.value,
                              return.means=return.means,
                              var.tables=list(B=B.var.table,
                                              MB=MB.var.table,
                                              DAMB=DAMB.var.table))))
}

## Draws and scales residuals in single function
calc.scaled.residual <- function(noise.effect.size, n){
  residual <- rnorm(n=n)
  residual <- (residual - mean(residual))/sqrt(non.sample.var(residual))
  residual <- residual*sqrt(noise.effect.size)
  return(residual)
}

calc.qtl.effect <- function(beta,
                            M,
                            A,
                            D,
                            Z,
                            strain.effect.size,
                            noise.effect.size) {
  MB <- M %*% beta
  DAMB <- D %*% A %*% MB
  ZDAMB <- Z %*% DAMB
  
  B.effect <- non.sample.var(2*beta)
  MB.effect <- non.sample.var(2*MB)
  DAMB.effect <- non.sample.var(DAMB)
  
  summaries <- c(B.effect, MB.effect, DAMB.effect,
                 B.effect/(B.effect + strain.effect.size + noise.effect.size),
                 MB.effect/(MB.effect + strain.effect.size + noise.effect.size),
                 DAMB.effect/(DAMB.effect + strain.effect.size + noise.effect.size))
  names(summaries) <- c("B.effect", "MB.effect", "DAMB.effect",
                        "B.ve", "MB.ve", "DAMB.ve")
  return(list(QTL.predictor = ZDAMB,
              summaries = summaries))
}


take.fixef.residuals <- function(data, 
                                 locus.matrix, 
                                 strain.matrix){
  locus.matrix <- locus.matrix[,-which.max(colSums(locus.matrix))]
  rownames(locus.matrix) <- NULL
  colnames(locus.matrix) <- paste("allele", 1:ncol(locus.matrix), sep=".")
  
  strain.matrix <- strain.matrix[,-1]
  
  temp.data <- data.frame(data, locus.matrix, strain.matrix)
  this.formula <- formula(paste("sim.y.1~1", 
                               paste(paste("allele", 1:ncol(locus.matrix), sep="."), collapse="+"),
                               paste(colnames(strain.matrix), collapse="+"),
                               sep="+"))
  this.fit <- lm(this.formula, data=temp.data)
  
  drop.na <- is.na(this.fit$coefficients[colnames(strain.matrix)])
  
  these.residuals <- data$sim.y.1 - strain.matrix[,!drop.na] %*% this.fit$coefficients[colnames(strain.matrix)][!drop.na]
  return(these.residuals)
}

take.ranef.residuals <- function(data, locus.matrix){
  locus.matrix <- locus.matrix[,-which.max(colSums(locus.matrix))]
  rownames(locus.matrix) <- NULL
  colnames(locus.matrix) <- paste("allele", 1:ncol(locus.matrix), sep=".")
  
  temp.data <- data.frame(data, locus.matrix)
  this.formula <- formula(paste0("sim.y.1~", 
                                paste(paste("allele", 1:ncol(locus.matrix), sep="."), collapse="+"),
                                "+(1|SUBJECT.NAME)"))

  this.fit <- lme4::lmer(this.formula, data=temp.data)
  these.residuals <- data$sim.y.1 - lme4::ranef(this.fit)$SUBJECT.NAME[data$SUBJECT.NAME,]
  return(these.residuals)
}

## From Wes. Returns SDP matrix and QTL effects
simulate.QTL.model.and.effects <- function(num.alleles=8, 
                                           num.founders=8, 
                                           M=NULL,
                                           sample.as.method=c("uniform", "crp"), 
                                           beta=NULL,
                                           ...){
  
  sample.as.method <- sample.as.method[1]
  if(is.null(M)){
    M <- matrix(0, num.founders, num.alleles)
    M[cbind(sample(1:num.founders, num.alleles), 1:num.alleles)] <- 1
    
    if(sample.as.method == "uniform"){
      M[which(rowSums(M)==0),] <- t(rmultinom(num.founders - num.alleles, 1, rep(1/num.alleles, num.alleles)))
    }
    else if(sample.as.method == "crp"){
      for(i in which(rowSums(M)==0)){
        M[i,] <- rmultinom(1, 1, colSums(M)/sum(M))
      }
    }
  }
  
  if(is.null(beta)){
    beta <- rnorm(num.alleles)
  }

  effect <- list(M=M, 
                 beta=beta)
  return(effect)
}

## Produce SDP mapping matrix from SDP string
model.matrix.from.ID <- function(M.ID){
  m <- as.numeric(unlist(strsplit(M.ID, ","))) + 1
  J <- length(m)
  M <- matrix(0, J, max(m))
  M[cbind(1:J, m)] <- 1
  return(M)
}

## Produces an incidence matrix from a factor
incidence.matrix <- function(fact){
  m <- diag(nlevels(fact))[fact,]
  colnames(m) <- levels(fact)
  return(m)
}

#' @export extract.compact.qr
extract.compact.qr <- function(genomecache, 
                               CC.lines.matrix,
                               use.progress.bar=TRUE){
  
  results <- list(qr.alt.list=list(), CC.lines.matrix=CC.lines.matrix)
  for(i in 1:ncol(CC.lines.matrix)){
    this.qr <- extract.qr(genomecache=genomecache, 
                          model="additive",
                          formula=~1, 
                          id="SUBJECT.NAME.1", 
                          data=data.frame(SUBJECT.NAME.1=CC.lines.matrix[,i]),
                          use.progress.bar=use.progress.bar)
    results$qr.alt.list[[i]] <- this.qr$qr.list
    if(i == 1){
      results$shared <- list(intercept.allele=this.qr$intercept.allele,
                             condition.loci=this.qr$condition.loci,
                             qr.0=this.qr$qr.0,
                             chr=this.qr$chr,
                             pos=this.qr$pos,
                             model=this.qr$model,
                             founders=this.qr$founders,
                             formula=this.qr$formula)
    }
  }
  return(results)
}

#' @export pull.qr.from.compact
pull.qr.from.compact <- function(compact.qr.list, 
                                 qr.index){
  
  output.qr <- list(qr.list=compact.qr.list$qr.alt.list[[qr.index]],
                    intercept.allele=compact.qr.list$shared$intercept.allele,
                    condition.loci=compact.qr.list$shared$condition.loci,
                    qr.0=compact.qr.list$shared$qr.0,
                    chr=compact.qr.list$shared$chr,
                    pos=compact.qr.list$shared$pos,
                    model=compact.qr.list$shared$model,
                    founders=compact.qr.list$shared$founders,
                    subjects=compact.qr.list$CC.lines.matrix[,qr.index],
                    formula=compact.qr.list$shared$formula)
  return(output.qr)
}

### From miqtl originally
extract.qr <- function(genomecache, 
                       id="SUBJECT.NAME",
                       data, 
                       formula, 
                       model=c("additive", "full"), 
                       condition.loci=NULL,
                       chr="all", 
                       just.these.loci=NULL, 
                       use.progress.bar=TRUE){
  K <- NULL
  model <- model[1]
  
  h <- DiploprobReader$new(genomecache)
  founders <- h$getFounders()
  num.founders <- length(founders)
  loci <- h$getLoci()
  
  loci.chr <- h$getChromOfLocus(loci)
  if(chr[1] != "all"){
    loci.chr <- h$getChromOfLocus(loci)
    loci <- loci[loci.chr %in% chr]
  }
  if(!is.null(just.these.loci)){
    loci <- loci[loci %in% just.these.loci]
    loci.chr <- loci.chr[loci %in% just.these.loci]
  }
  subjects <- as.character(data[,id])
  X.0 <- model.matrix(formula, data=data)
  if(!is.null(condition.loci)){
    for(i in 1:length(condition.loci)){
      X.condition <- h$getLocusMatrix(condition.loci[i], model=model, subjects=subjects)
      keep.col <- 1:ncol(X.condition)
      max.column <- which.max(colSums(X.condition, na.rm=TRUE))[1]
      keep.col <- keep.col[keep.col != max.column]
      X.0 <- cbind(X.0, X.condition[,keep.col])
    }
  }
  qr.0 <- qr(X.0)
  
  if(use.progress.bar){ pb <- txtProgressBar(min=0, max=length(loci), style=3) }
  qr.list <- list()
  intercept.allele <- rep(NA, length(loci)) # For allele effects
  for(i in 1:length(loci)){
    X <- h$getLocusMatrix(loci[i], model=model, subjects=subjects)
    keep.col <- 1:ncol(X)
    max.column <- which.max(colSums(X, na.rm=TRUE))[1]
    intercept.allele[i] <- founders[max.column]
    keep.col <- keep.col[keep.col != max.column]
    X <- cbind(X.0, X[,keep.col])
    qr.list[[i]] <- qr(X)
    if(use.progress.bar){ setTxtProgressBar(pb, i) }
  }
  names(qr.list) <- loci
  
  qr.object <- list(qr.list=qr.list,
                    intercept.allele=intercept.allele,
                    condition.loci=condition.loci,
                    qr.0=qr.0,
                    chr=loci.chr,
                    pos=list(cM=h$getLocusStart(loci, scale="cM"),
                             Mb=h$getLocusStart(loci, scale="Mb")),
                    model=model,
                    founders=h$getFounders(),
                    subjects=subjects,
                    formula=Reduce(paste, deparse(formula)))
}

### From miqtl originally
scan.qr <- function(qr.object, 
                    data, 
                    phenotype,
                    return.allele.effects=FALSE,
                    chr="all", 
                    id="SUBJECT.NAME",
                    just.these.loci=NULL,
                    debug.single.fit=FALSE, 
                    use.progress.bar=TRUE,
                    ...){
  model <- qr.object$model
  subjects <- qr.object$subjects
  founders <- qr.object$founders
  num.founders <- length(founders)
  loci <- names(qr.object$qr.list)
  loci.chr <- qr.object$chr
  rh.formula <- qr.object$formula
  
  if(model == "full" & return.allele.effects){
    return.allele.effects <- FALSE
    cat("Allele effects from regression models currently only available with additive model\n",
        "Setting return.allele.effects to FALSE\n")
  }
  
  ## Matching the subject order in the data with the qr object
  reorder <- match(subjects, data[,id])
  data <- data[reorder,]
  n <- nrow(data)
  
  if(chr[1] != "all"){
    loci <- loci[loci.chr %in% chr]
  }
  if(!is.null(just.these.loci)){
    loci <- loci[loci %in% just.these.loci]
    loci.chr <- loci.chr[loci %in% just.these.loci]
  }
  
  formula.string <- paste(phenotype, rh.formula)
  formula <- formula(formula.string)
  
  allele.effects <- NULL
  p.vec <- rep(NA, length(loci))
  
  if(return.allele.effects){ 
    allele.effects <- matrix(NA, nrow=length(founders), ncol=length(loci),
                             dimnames=list(founders, loci))
  }
  y <- as.vector(model.frame(formula, data=data)[,1])
  names(y) <- subjects
  # Progress bar
  if(use.progress.bar){ pb <- txtProgressBar(min=0, max=length(loci), style=3) }
  for(i in 1:length(loci)){
    p.vec[i] <- get.f.stat.p.val(qr.alt=qr.object$qr.list[[i]], 
                                 qr.null=qr.object$qr.0, 
                                 y=y)
    if(is.nan(p.vec[i])){ p.vec[i] <- 1 }
    if(return.allele.effects){
      allele.effects[,i] <- get.allele.effects.from.fixef.eQTL(qr.alt=qr.object$qr.list[[i]], 
                                                               y=y, 
                                                               founders=founders,
                                                               intercept.allele=qr.object$intercept.allele[i])
    }
    if(debug.single.fit){ browser() }
    # Update progress bar
    if(use.progress.bar){ setTxtProgressBar(pb, i) }
  }
  names(p.vec) <- loci
  if(!is.null(qr.object$condition.loci)){
    formula.string <- paste(Reduce(paste, deparse(formula)), paste(qr.object$condition.loci, collapse="+"), sep="+")
  }
  output <- list(LOD=NULL,
                 p.value=p.vec,
                 df=NULL,
                 pos=list(Mb=qr.object$pos$Mb, 
                          cM=qr.object$pos$cM),
                 loci=loci, 
                 chr=loci.chr,
                 allele.effects=allele.effects,
                 y=y,
                 formula=formula.string,
                 model.type=model,
                 p.value.method="ANOVA",
                 locus.effect.type="fixed",
                 n=length(y))
  return(output)
}

### From miqtl originally
straineff.mapping.matrix <- function(M=8){
  T <- M*(M+1)/2
  mapping <- matrix(rep(0, T*M), M, T)
  idx <- 1;
  for (i in 1:M){
    mapping[i, idx] <- mapping[i, idx] + 2
    idx <- idx + 1;
  }
  for (i in 2:M){
    for (j in 1:(i-1)){
      mapping[i, idx] <- mapping[i, idx] + 1;
      mapping[j, idx] <- mapping[j, idx] + 1;
      idx <- idx + 1;
    }
  }
  return(t(mapping))
}

#' Simulate Collaborative Cross (CC) phenotype data from the realized CC lines founder haplotype probabilities 
#' stored in a genome cache directory with a correlated strain effect
#'
#' This function takes various input parameters to simulate CC data with a correlated strain effect based on 
#' genetic relatedness to be used in power calculations or as input data for other tools that analyze CC data.
#'
#' @param genomecache The path to the genome cache directory. The genome cache is a particularly structured
#' directory that stores the haplotype probabilities/dosages at each locus. It has an additive model
#' subdirectory and a full model subdirectory. Each contains subdirectories for each chromosome, which then
#' store .RData files for the probabilities/dosages of each locus.
#' @param CC.lines DEFAULT: NULL. If NULL is specified, sim.CC.data.K() will randomly draw samples of the 
#' available CC lines of the size specified in num.lines.
#' @param num.lines DEFAULT: NULL. If NULL, sim.CC.data.K() expects that CC.lines is non-NULL. If CC.lines
#' is NULL, num.lines determines the number of CC lines that are sampled.
#' @param vary.lines DEFAULT: TRUE. If CC.lines is NULL and vary.lines is TRUE, then sim.CC.data.K() will
#' sample multiple sets of CC lines of the size specified in num.lines. If CC.lines is NULL and vary.lines is
#' FALSE, then just one set of CC lines will be sampled.
#' @param locus DEFAULT: NULL. If NULL is specified, sim.CC.data.K() will randomly draw a locus stored in
#' the genome cache. 
#' @param vary.locus DEFAULT: TRUE. If locus is NULL and vary.locus is TRUE, then sim.CC.data.K() will sample
#' as many loci as specified in num.sim. If locus is NULL and vary.locus is FALSE, then sim.CC.data.K() will
#' only sample one locus.
#' @param K The kinship matrix. A realized genetic relationship matrix is commonly used.
#' @param num.replicates The number of replicates of each CC line that will be simulated. Mapping for power calculations
#' will use strain means. Currently requires that all lines have the same number of replicates.
#' @param num.sim The number of phenotypes to be simulated for a given parameter setting.
#' @param M.ID DEFAULT: NULL. M is a matrix that maps from counts of founder haplotypes to counts of functional 
#' alleles. Mapping will be based on haplotype association, but potentially there are two to the number of
#' founders alleles at the QTL. M.ID is a string that codifies this mapping. One potential balanced two allele
#' M.ID would be "c(0,0,0,0,1,1,1,1)". With 8 functional alleles, on per founder, the only M.ID is "c(0,1,2,3,4,5,6,7)".
#' If M.ID is NULL, M.ID will be sampled.
#' @param sample.as.method DEFAULT: "uniform". The procedure used for sampling the allelic series. If
#' every strain has its own allele, this option does not matter. Alternatively, a Chinese restaurant process ("crp") can be used,
#' which is possibly more biologically accurate, and will favor allelic series that are less balanced (1 vs 7).
#' @param num.alleles DEFAULT: 8. The number of functional alleles. Must be less than or equal to the number of 
#' founders.
#' @param num.founders DEFAULT: 8. The number of founders, which must correspond to the genome cache. The CC has
#' eight.
#' @param qtl.effect.size The size of the simulated QTL effect. The scale of the input is in proportion of
#' the phenotypic variance due to the QTL, thus should be greater than or equal to zero, and less than one.
#' @param beta DEFAULT: NULL. Allows for the manual specification of QTL effect. Is expected to be a vector
#' the length of the number of alleles. It will be scaled based on qtl.effect.size.
#' @param strain.effect.size DEFAULT: 0. The size of the simulated strain effect, which represents something akin to a polygenic 
#' effect. Other variants specific to CC lines will result in overall strain-specific effects. The scale of the input 
#' is in proportion of the phenotypic variance due to the strain, thus should be greater than or equal to zero, and less than one.
#' @param impute DEFAULT: TRUE. If TRUE, the QTL portion of the design matrix in the simulation is a realized sampling
#' of haplotypes from the probabilities. If FALSE, the simulations are based on the probabilities, which is flawed in
#' terms of biological reality.
#' @param scale.qtl.mode DEFAULT: "B". Specifies how the QTL effect is scaled. If "B", then the variance of the qtl effect
#' vector beta is scaled to the effect size specified in qtl.effect.size, which would be the variance explained in a population
#' perfectly balanced in terms of the functional alleles. If "MB", the the variance of M %*% beta is scaled, which would be
#' the variance explained in a population balanced in terms of founder strains with the allelic series that is specified in M. 
#' If "DAMB", the variance of D %*% A %*% M %*% beta is scaled, which would be the variance explained in a population of a 
#' specific set of CC strains (specified in D).
#' @param return.value DEFAULT: "raw". If "raw", residuals are not taken. If "fixef.resid", then the data
#' are residuals after regressing phenotype on strain. If "ranef.resid", then the data have had the strain BLUP
#' effect subtracted.
#' @param return.means DEFAULT: TRUE. If TRUE, strain means are returned. If FALSE, the full data
#' with replicate observations of strains are returned.
#' @export sim.CC.data.K
#' @examples sim.CC.data.K()
sim.CC.data.K <- function(genomecache,
                          num.lines,
                          num.alleles,
                          num.sim,
                          num.replicates,
                          strain.effect.size,
                          K,
                          CC.lines = NULL,
                          ...) {
  
  vary.lines <- ifelse(num.lines == 72 | !is.null(CC.lines), FALSE, TRUE)
  
  sparcc_sims <- sim.CC.data(genomecache = genomecache, 
                             num.replicates = num.replicates,
                             num.lines = num.lines, 
                             num.alleles = num.alleles, 
                             qtl.effect.size = 0, 
                             strain.effect.size = 0, 
                             num.sim = num.sim,
                             vary.lines = vary.lines,
                             CC.lines = CC.lines)
  if (!vary.lines) { this_K <- K[CC.lines, CC.lines] }
  for (i in 1:num.sim) {
    if (vary.lines) {
      this_K <- K[sparcc_sims$data[,paste0("SUBJECT.NAME.", i)],
                  sparcc_sims$data[,paste0("SUBJECT.NAME.", i)]]
    }
    
    strain <- mvtnorm::rmvnorm(1, mean = rep(0, nrow(this_K)), sigma = this_K)[1,]
    
    strain <- (strain - mean(strain))/sqrt(sparcc:::non.sample.var(strain))
    strain <- strain * sqrt(strain.effect.size)
    
    sparcc_sims$data[, paste0("sim.y.", i)] <- strain
  }
  sparcc_sims$properties$qtl.effect.size <- 0
  sparcc_sims$properties$strain.effect.size <- strain.effect.size
  
  sparcc_sims
}
```
