## Supplementary material for "Determinants of QTL mapping power in the realized Collaborative Cross": SPARCC R package used for all analyses: DiploprobReaderClass.R

```
#### Class for a DiploprobReader object
#### Written by Will Valdar - from bagpipe.backend package
#' @export DiploprobReader
#' @exportClass DiploprobReader
DiploprobReader <- setRefClass("DiploprobReader",
  fields = c(
    "mHappy"
  ), # = "happy.genome"
  methods = list(
    #
    #
    initialize = function(dataDir, format=c("happy"), ...){
      if (inherits(dataDir, "happy.genome")){
        .self$mHappy <- dataDir
        warning("DiploprobReader initization from happy objects will be phased out in future versions.\n")
      } else if ("happy"==format[1]){
        .self$mHappy <- happy.load.genome(dataDir) 
      } else {
        stop("Unknown format ", format, "\n")
      }
      callSuper(...)
    },
#    getAllowedModels = function(){
#    },
    getChromEnd = function(chrom, scale){
      .self$getLocusEnd(tail(.self$getLoci(chrom), 1), scale=scale)
    },
    getChromStart = function(chrom, scale){
      .self$getLocusStart(.self$getLoci(chrom)[1], scale=scale)
    },
    getChromLength = function(chrom, scale){
      'Returns the length of the specified chromosome in units of the specified scale
      '
      happy.get.chromosome.length(.self$mHappy, chrom=chrom, scale=scale)
    },
    getChromList = function(){
      'Returns a character vector of the chromosome names
      '
      happy.list.chromosomes(.self$mHappy)
    },
    getChromOfLocus = function(loci){
      'Returns a character vector containing the name(s) of the chromosome(s) to which the specified loci belong
      '
      happy.get.chromosome(.self$mHappy, loci)
    },
    getGenotype = function(marker, ...){
      'Returns a vector containing the genotypes observed for the specified marker, with NA for
      missing genotypes'
      as.character(happy.get.genotype(.self$mHappy, marker=marker, genotype.model="factor"))
    },
    getLocusMatrix = function(locus, model, subjects=NULL, as.data.frame=FALSE, sdp=NULL){
      'Get matrix representing average probabilities or expectations of haplotypes
      over the interval
      '
      happy.get.design(.self$mHappy, marker=locus, model=model, as.data.frame=as.data.frame, sdp=sdp, subjects=subjects)
    },
    getLocusProbTensor = function(locus, model, subjects=NULL, simplify=FALSE, memoize.last=TRUE){
      '
      For n subjects descended from J strains, this function returns a tensor
      of n JxJ matrices. Each matrix gives the probability that a randomly
      chosen point within the locus interval is descended from a particular diplotype.
      
      model: When model = full.asymmetric, the diplotype probability matrix distinguishes
      between diplotype AB and BA for founders A and B. When model = full, these
      probabilities are set to be equal. Depending on how the HMM probabilities were
      estimated, full.asymmetric may not be available.
      
      simplify: When simplify=TRUE and only one subject is specified, the return value is
      simplified from a 1xJxJ tensor to a JxJ matrix.
      
      memoize.last: An optimization for when it is expected that the same tensor will be
      requested repeatedly. Setting memoize.last=TRUE causes the method to make
      an internal cache of the last return value. If the exact same tensor is requested,
      then the method returns the cached value, thereby avoiding the cost of repeating 
      various operations that may include file I/O.
      '
      if (missing(subjects)){
        subjects <- .self$getSubjects()
      }
      happy.get.diplotype.tensor(.self$mHappy, marker=locus, model=model, subjects=subjects, memoize=memoize.last, simplify=simplify)
    },
    getFirstLocus = function(chrom){
      'Returns the name of the first locus on the specified chromosome
      '
      happy.get.first.marker(.self$mHappy, chrom=chrom)
    },
    getFounders = function(){
      'Returns a character vector of the founder names
      '
      happy.get.strains(.self$mHappy)
    },
    getLastLocus = function(chrom){
      'Returns the name of the last locus on the specified chromosome
      '
      happy.get.last.marker(.self$mHappy, chrom=chrom, as.intervals=TRUE)
    },
    getLocusWidth = function(loci, scale){
      'Returns a numeric vector containing, for each specified locus, the left-to-right width in the
      units of the specified scale
      '
      happy.get.interval.length(.self$mHappy, loci, scale=scale)
    },
    getLoci = function(chrom=NULL, before=NULL, after=NULL, from=NULL, to=NULL, scale="interval", over=NULL){
      'Returns a character vector of the locus names
      '
      if (missing(before) & missing(after) & missing(from) & missing(to) & missing(scale)){
        return (happy.get.markers(.self$mHappy, chrom=chrom))
      } else {
        warning("Incompletely debugged code for getLoci\n")
      }
      if (!is.null(from)) {
        return (happy.get.intervals.in.range(h, chromosome=chrom, markers=loci, from=from, to=to))
      } else if (!is.null(over)) {
        return (happy.get.interval.over(.self$mHappy, chromosome=chrom, scale=scale))
      } else {
        return (happy.get.markers.between(.self$mHappy, before=before, after=after, as.intervals=TRUE))
      }
    },
    getMarkers = function(){
      '
      '
      happy.get.markers(.self$mHappy, model="genotype")
    },
    getLociInRange = function(from=NULL, to=NULL, scale="locus", chr=NULL){
      happy.get.intervals.in.range(.self$mHappy, from=from, to=to, chr=chr, 
        scale=ifelse("locus"==scale, "interval", scale))
    },
    getLocusOver = function(x, scale, chr){
      happy.get.interval.over(.self$mHappy, x, scale=scale, chr=chr)
    },
    getLocusEnd = function(loci, scale){
      .self$getLocusRange(loci, scale)[, 2]
    },
    getLocusStart = function(loci, scale){
      .self$getLocusRange(loci, scale)[, 1]
    },
    getLocusRange = function(loci, scale){
      'Returns the left and right boundaries of the specified loci, in units of the specified scale
      '
      happy.get.interval.range(.self$mHappy, markers=loci, scale=scale)
    },
    getMarkerLocation = function(markers, scale){
      'Returns the location of the specified marker(s) in units of the specified scale
      '
      happy.get.location(.self$mHappy, markers, scale=scale)
    },
		getNextMarker = function(markers){
			'Returns the next marker along, or NA if at the end of the chromosome
			'
			happy.get.next.marker(.self$mHappy, markers)
		},
    getNumFounders = function(){
      'Returns the number of founders
      '
      length(.self$getFounders())
    },
    getNumSubjects = function(){
      'Returns the number of subjects
      '
      length(.self$getSubjects())
    },
    getSubjects = function(){
      'Returns a character vector of the subject names
      '
      happy.get.subjects(.self$mHappy)
    },
    hasChrom = function(chrom){
      'Returns logical vector indicating whether the specified chromosomes exist
      '
      happy.has.chromosome(.self$mHappy, chrom)
    },
    hasLoci = function(loci){
      'Returns logical vector indicating whether loci with the specified names exist
      '
      happy.has.markers(.self$mHappy, loci)
    },
    hasMarkers = function(markers){
      'Returns logical vector indicating whether markers with the specified names exist. Compare $hasLoci()$.
      '
      markers %in% happy.get.markers(.self$mHappy, model = "genotype", as.intervals = FALSE)
    },
    hasSubjects = function(subjects){
      'Returns logical vector indicating whether subjects with the specified names exist
      '
      happy.has.subjects(.self$mHappy, subjects)
    },
    help = function(...){
      DiploprobReader$help(...)
    },
    methods = function(...){
      DiploprobReader$methods(...)
    },
    updateBp = function(markerBpTable, allowPartial=FALSE){
      'Update base pair positions of the markers. Handy when bp positions were unavailable or misspecified at the time of generating the genome cache. $markerBpTable$ is a data.frame with two columns: the first should have the locus name, the second should have the bp position. $allowPartial$ permits updating of a subset of the markers; by default this is set to FALSE.
      '
      # ideally this should be modifying one global lookup, but the current happy object is not structured that way.
      # TODO: store bp information in *one* central location, not duplicated in every model!!!
      models <- setdiff(names(.self$mHappy), c("subjects", "strains", "markers"))
      for (model in models){
        tab <- data.frame(markers=I(as.character(markerBpTable[, 1])), bp=as.integer(markerBpTable[, 2]))
        drMarkers <- .self$mHappy[[model]]$genome$marker
        tab <- tab[tab$markers %in% drMarkers, ]
        if (!allowPartial){
          missingMarkers <- setdiff(tab$markers, drMarkers)
          if (0<length(missingMarkers)){
            stop("Cannot update base pairs in method $updateBp()$: new marker data is short by ", length(missingMarkers), " markers. Either give complete set or set allowPartial flag to true")
          }
        }
        #
        i.replace <- match(tab$markers, drMarkers)
        .self$mHappy[[model]]$genome$bp[i.replace] <- tab$bp
      }
    }
  ) # end method defs
) # end ref class def
    
  
  
```
