## Supplementary material for "Determinants of QTL mapping power in the realized Collaborative Cross": SPARCC R package used for all analyses: read.happy.core.R

```
#### Functions to interact with happy genomecache
#### Written by Will Valdar - from bagpipe.backend package

assert.happy <- function(h)
{
	if (!inherits(h, "happy.genome"))
	{
		stop("Object must be of class happy.genome\n")
	}
}

happy.get.allowed.models <- function()
{
	c("genotype", "additive", "full", "full.asymmetric")
}	

happy.get.bp <- function(ha, markers)
{
  ha$genotype$genome$bp[
      match(markers, ha$genotype$genome$marker)
      ]
}

happy.get.chromosome <- function(h, markers)
{
	assert.happy(h)
  h$genotype$genome$chromosome[match(markers, h$genotype$genome$marker)]
}

happy.get.interval.length <- function(h, markers, scale="bp", fudge.bp=FALSE)
# return the length of the interval
{
  d <- rep(NA, length(markers))

  i <- match(markers, h$genotype$genome$marker)
  if ("bp"==scale | "Mb"==scale)
  {
    start <- h$genotype$genome$bp[i]
    end   <- h$genotype$genome$bp[i+1]
    d <- end - start
    ok=!is.na(d)
    if (any(d[ok] < 0))
    {
      if (fudge.bp)
      {
        stop("Fudging is currently deprecated\n")
        # find next bp on same chrom that is higher than current bp
        fudges <- which(d < 0)
        for (f in fudges)
        {
          i.end <- which( h$genotype$genome$bp > h$genotype$genome$bp[i[f]]
              & h$genotype$genome$chromosome == h$genotype$genome$chromosome[i[f]]
                  )[1]
          d[f] <- h$genotype$genome$bp[i.end] - start[f]
        }
      }
      else
      {
        warning("Cannot calculate lengths for the following intervals",
            " because their right-flank markers have a lower bp than their",
            " left-flank markers:",
            paste( markers[which(d < 0)], collapse=", "),
            "\n")
        d[ 0 > d ] <- NA
      }
    }
    if ("Mb"==scale) d <- d/1e6
  }
  else
  {
    start <- h$genotype$genome$map[i]
    end   <- h$genotype$genome$map[i+1]
    d     <- end - start
    d[ 0 > d ] <- NA # deals with intervals of negative length
  }
  return (d)
}


happy.get.markers <- function(h,
		chromosome = NULL,
		model      = "genotype",
		as.intervals = TRUE)
{
	assert.happy(h)
	if (!all(happy.has.model(h, unique(c(model, "genotype")))))
	{
		i <- happy.has.model(h, unique(c(model, "genotype")))
		stop("Model ", model[!i], " not loaded\n")
	}
	
	# optimized case
	if (is.null(chromosome))
	{
		if (!as.intervals)
		{
			return ( h$genotype$markers )
		}
		if ("genotype"!=model && as.intervals)
		{
			return ( h[[model]]$markers )
		}
	}
	
	# inefficient but general
	markers <- h$genotype$markers
	terminii <- tapply(1:length(h$genotype$marker), h$genotype$chromosome, tail, 1)
	is.terminus <- rep(FALSE,length(markers))
	is.terminus[terminii] <- TRUE
	if (is.null(chromosome)) chromosome <- unique(h$genotype$chromosome)
	
	if ("genotype"==model)
	{
		if (as.intervals)
		{
			return (markers[
			    !is.terminus
			    & h$genotype$chromosome %in% chromosome
			    ])
		}
		return (markers[ h$genotype$chromosome %in% chromosome ])
	}
	if (as.intervals)
	{
		return (h[[model]]$markers[ h[[model]]$chromosome %in% chromosome ])
	}
	return (
	  happy.get.markers(h,
	    chromosome=chromosome,
	    model="genotype",
	    as.intervals=FALSE)
	  )
}	

happy.get.models <- function(h)
{
	assert.happy(h)
	intersect(names(h), happy.get.allowed.models())
}

happy.get.position <- function(h, markers)
{
	assert.happy(h)
  h$genotype$genome$map[
            match(markers, h$genotype$genome$marker)
            ]
}

happy.get.subjects <- function(h)
{
	h$subjects
}

happy.get.strains <- function(h)
{
  h$strains
}

happy.has.model <- function(h, model)
{
	assert.happy(h)
  model %in% happy.get.models(h)
}

happy.load.genome <- function (dir, use.X = TRUE, chr = NULL, models = NULL)
{
  if (!is.null(chr) & 0==length(grep("chr", chr)))
  {
    chr <- paste("chr",sep="",chr)
  }
  if (is.null(models))
  {
    models <- intersect(happy.get.allowed.models(), list.subdirs(dir))
	  if (0==length(models))
	  {
	    stop("No genome cache models present in ", dir, "\n")
	  }
	  if (!"genotype" %in% models)
	  {
	    stop("Required genotype model is absent from cache dir ", dir, "\n") 
	  }
  }
	else
	{
		models <- unique(c(models, "genotype"))
	}

  g <- list()
  old.subjects <- NULL
  old.strains <- NULL
  for (model in models)
	{
    pkgs <- c()
    if (is.null(chr))
    {
      found.chr <- list.subdirs( paste(dir, "/", model, sep=""), pattern="^chr" )
  	  if (0==length(found.chr))
  		{
  			stop("Found no chromosomes for model ", model, " in dir ", dir, "\n")
  		}
      pkgs <- paste(dir, model, found.chr, sep = "/")
    }
    else
    {
      pkgs <- paste(dir, model, chr, sep = "/")
    }
    markers <- c()
    chromosome <- c()
    map <- c()
    pkgname <- c()
    bp <- c()
    for (p in pkgs)
		{
      chromosome <- c(chromosome, happy.load.data("chromosome", p))
      m <- happy.load.data("markers", p)
      markers <- c(markers, m)
      map <- c(map, happy.load.data("map", p))
      bp <- c(bp, happy.load.data("bp", p))
      pkgname <- c(pkgname, rep(p, length(m)))
      subjects <- happy.load.data("subjects", p)
      strains <- happy.load.data("strains", p)
      if (is.null(old.subjects))
      {
          old.subjects <- subjects
      }
      if (any(subjects != old.subjects))
      {
          cat("ERROR - subject names are inconsistent for chromosome ",
            tail(chromosome,1), "\n")
          stop("FATAL HAPPY ERROR")
      }
      if (is.null(old.strains))
      {
          old.strains <- strains
      }
      if (any(strains != old.strains))
      {
          cat("ERROR - strain names are inconsistent for chromosome ",
            chromosome, "\n")
          stop("FATAL HAPPY ERROR")
      }
    }
    genome <- data.frame(
		marker = I(as.character(markers)),
        	map = as.numeric(map),
		bp = as.numeric(bp),
		ddp = I(as.character(pkgname)),
        	chromosome = I(as.character(chromosome)))
    g[[model]] <- list(
		genome = genome,
		subjects = subjects,
        	strains = strains,
		markers = as.character(genome$marker),
        	chromosome = as.character(genome$chromosome),
		map = genome$map,
		design.matrix.colnames = happy.make.colnames(strains, model=model))
  }
  g$subjects <- g$genotype$subjects
  g$strains <- g$additive$strains
  g$markers <- g$genotype$markers
  g$haploid <- g$genotype$haploid
  
  class(g) <- "happy.genome"
  return(g)
}

happy.load.marker <- function(h, marker, model)
# internal function to load data for a single marker from genome cache
{
  # Check for bad arguments
	assert.happy(h)
	if (1!=length(marker))
  {
      stop("Must specify only one marker in happy.load.marker()\n")
  }
  if (1!=length(model))
  {
      stop("Must specify only one model in happy.load.marker()\n")
  }
  if (!happy.has.model(h, model))
  {
      stop("No such model ", model, " in happy object\n")
  }
	marker <- as.character(marker)
	model  <- as.character(model)

	# check whether marker is in DATA memory
	retval <- NULL
	if ( happy.has.reserved.marker(h, marker=marker, model=model) )
	{
		retval <- happy.get.reserved.marker(h, marker=marker, model=model)
	}
	else 
	{
	    i <- which(h[[model]]$genome$marker == marker )
	    if (1!=length(i))
	    {
	        string <- paste("marker", marker, "for ", model, " model.")
	        if (0==length(i))
	        {
	            stop("Could not find ", string, "\n")
	        }
	        if (1 < length(i))
	        {
	            stop("Found multiple markers matching", string, "\n")
	        }
	    }
	    pkg <- h[[model]]$genome$ddp[i]

	    max.tries  <- 10
	    num.tries  <- 0
	    sleep.time <- 1
	    has.loaded <- FALSE
	    retval     <- NULL
	    while (!has.loaded & num.tries < max.tries)
	    {
	        ## read marker data from the genome cache
	        num.tries <- num.tries + 1
	        retval <- try( happy.load.data(marker, pkg) )
	        if (!caught.error(retval))
	        {
	            if (!is.null(retval))
	            {
	                break
	            }
	        }
	        warning("Failed to load data for ", model, " ", marker,
	                ". Retrying after ", sleep.time, "s sleep...\n")
	        system(paste("sleep", sleep.time))
	    }
	    if (caught.error(retval))
	    {
	        stop("Error retrieving information via g.data.get() for marker ", marker, "\n")
	    }
	    if (is.null(retval))
	    {
	        stop("Error: null data retrieved via g.data.get() for marker ", marker, "\n")
	    }
	    if (!is.array(retval)) retval <- as.array(retval)    # ensure return value has a dim component
		colnames(retval) <- happy.make.colnames(happy.get.strains(h), model)
		
		if ("full"==model)
		# bug fix for when cache contains incorrectly doubled values
		{
			if (2==round(sum(retval[1,]),1))
			{
				retval <- retval/2
			}
		}
	}
	if (happy.is.auto.reserve(h))
	{
		if (happy.get.reserve.limit(h) > happy.reserve.memory.usage(h))
		{
			happy.reserve.marker(h, marker=marker, model=model, marker.data=retval)
		}
	}
  retval
}

happy.load.data <- function (item, dir) # replaces calls to g.data.get, to make things backwards compatible.
{
    env <- new.env()

    # determine which version of g.data was used to save the data
	# assume happy pre 2009 and g.data pre 2009
    filename.pre2009  <- file.path(dir, "data", paste(item, "RData", sep = "."))
    if (file.exists(filename.pre2009))
    {
        load(filename.pre2009, env)
        return ( get(item, envir = env) )
    }
    
    # assume happy 2009 and g.data 2009
	item.safe <- make.names(item)
    filename.post2009 <- file.path(dir,
            paste(gsub("([[:upper:]])", "@\\1", item.safe), "RData", sep = ".")
            )
    if (file.exists(filename.post2009))
    {       
        load(filename.post2009, env)
        return ( get(item.safe, envir = env ) )
    }

	# assume happy 2009 and g.data pre 2009
	filename.hybrid <- file.path(dir, "data", paste(item.safe, "RData", sep = "."))
    if (file.exists(filename.hybrid))
    {
        load(filename.hybrid, env)
        return ( get(item.safe, envir = env) )
    }

    stop("Could not find object file containing data for ", item, " in package ", dir,
 			". Tried ", filename.pre2009, ", ", filename.post2009, " and ", filename.hybrid, "\n")
}

happy.reserve.marker <- function(h, marker, model, marker.data=NULL)
{
	if (!happy.reserve.has(h, category=model))
	{
		stop("Cannot reserve marker ", marker, " for model ", model, " because memory cache has not been initialized\n")
	}
	if (is.null(marker.data))
	{
		marker.data <- happy.load.marker(h, marker=marker, model=model)
	}
	happy.reserve.put(h, category=model, object.name=marker, object=marker.data)
}

happy.reserve.markers <- function(h, markers, models, verbose=TRUE)
{
	assert.happy(h)
	if (length(markers)!=length(models))
	{
		stop("Number of markers must match number of models\n")
	}
	
	markers <- as.character(markers)
	models  <- as.character(models)
	
	cat("Reserving data for ", length(markers), " marker-model combinations in memory\n") 

	mem.size <- 0
	memory.limit.Mb <- happy.get.reserve.limit(h)
	if (0<length(h$DATA))
	{
		mem.size <- happy.reserve.memory.usage(h)/2^20
	}
	for (i in 1:length(markers))
	{
		if (happy.has.reserved.marker(h, marker=markers[i], model=models[i]))
		{
			next
		}

		marker.data <- happy.load.marker(h, markers[i], models[i])
		mem.size <- mem.size + object.size(marker.data)/2^20
		if (memory.limit.Mb <= mem.size)
		{
			warning(paste("Reached memory limit",round(mem.size,3),"Mb / ",
					memory.limit.Mb,"Mb for marker reserve with",
					i, "/",length(markers),"markers. The remaining", length(markers)-i,
					"markers will be accessed through disk I/O\n"))
			break
		}
		happy.reserve.marker(h, marker=markers[i], model=models[i], marker.data=marker.data)
		if (verbose)
		{
			cat("[",i,"]",sep="")
		}
	}
	
	if (verbose) cat("\n")
	cat("Marker data consumes", round(mem.size,3), "Mb\n")
}

happy.has.reserved.marker <- function(h, marker, model)
{
  happy.reserve.has(h, category=model, object.name=marker)
}

happy.get.reserved.marker <- function(h, marker, model)
{
  happy.reserve.get(h, category=model, object.name=marker)
}
```
