## Supplementary material for "Determinants of QTL mapping power in the realized Collaborative Cross": SPARCC R package used for all analyses: read.happy.derived.R

```
#### Written by Will Valdar - from bagpipe.backend package

happy.check.bp <- function(h, stop.on.fail=TRUE)
{
    WARN <- stop
    if (!stop.on.fail) WARN <- warning

    assert.happy(h)

    ok <- TRUE

    # check markers are in map order
    for (chr in happy.list.chromosomes(h))
    {
        pos <- happy.get.position(h,
                happy.get.markers(h, chromosome=chr))
        if (any(order(pos)!=1:length(pos)))
        {
            ok <- FALSE
            WARN("Disorder in internal representation: ",
                    "markers are not in cM order\n")
        }
    }

    # check markers have non-NA basepairs
    all.markers <- happy.get.markers(h)
    if (any(is.na(happy.get.bp(h, all.markers))))
    {
        ok <- FALSE
        WARN("Some markers with NA bp\n")
    }
    ok
}


happy.get.allele.freq <- function(h, markers, subjects=NULL)
{
    freqs <- rep(NA, length(markers))
    names(freqs) <- markers
    for (m in markers)
    {
        f <- mean(na.omit(
            happy.get.genotype(h, m, model="additive", subjects=subjects)
            ))/2
        freqs[m] <- min(f, 1-f)
    }
    return (freqs)
}

happy.get.chromosome.length <- function(h, chrom, scale="bp", subtract.offset=FALSE)
{
    out <- rep(NA, length(chrom))
    for (ic in 1:length(chrom))
    {
        i <- h$genotype$genome$chr == chrom[ic]

        rng <- NULL
        if ("bp"==scale | "Mb"==scale)
        {
            rng <- range(h$genotype$genome$bp[i])
            if ("Mb"==scale) rng <- rng/1e6
        }
        else if ("cM"==scale)
        {
            rng <- range(h$genotype$genome$map[i])
        }
        else
        {
            stop("Unknown scale type ", scale, "\n")
        }

        if (subtract.offset)
        {
            return (rng[2] - rng[1])
        }
        out[ic] <- rng[2]
   }
   out
}


happy.get.design.old <- function(h, marker,
        model="additive",
        subjects=NULL,
        as.data.frame=TRUE)
{
  assert.happy(h)
  
	# Specify different genotype models
  if ("genotype"==model)
	{
		model <- "genotype.full"
	}
	hmodel   <- model
	submodel <- NULL
	if (igrep("genotype", model))
	{   
		submodel <- sub("genotype.", "", model)
		hmodel   <- "genotype"
	}

	# prepare dominance model
	if ("dominance"==model)
	{
		hmodel <- "full"
	}
    
	# Load the marker data from file
  mat <- happy.load.marker(h, marker=marker, model=hmodel)

	# Reformat different genotype submodels, if applicable
	if ("genotype"==hmodel)
	{
    if (submodel %in% c("additive", "dominance", "hier") )
    {
      mat <- as.matrix(genotype.to.hier(as.vector(mat)))
    	if ("additive"==submodel) { mat <- mat[,1] }
    	else if ("dominance"==submodel) { mat <- mat[,2] }
    }
    else if ("full"==submodel)
    {
        mat <- as.factor(mat)
    }
    else if ("ped"==submodel)
    {
        g <- genotype.to.count(as.vector(mat))
        mat <- rep("00", length(g))
        mat[ g==0 ] <- 11
        mat[ g==1 ] <- 12
        mat[ g==2 ] <- 22
    }
		else
		{
			stop("Unknown model: ", model, "\n")
		}
	}
  # force to be array if 1 column
  if (is.null(dim(mat)))
	{
		mat <- as.array(mat)
	}
  if (!is.null(subjects))
  {
    subjects <- as.character(subjects)
    i <- match(subjects, happy.get.subjects(h))
    if (1==length(dim(mat)))
    {
      mat <- mat[i]
    }
    else
    {
      mat <- matrix(mat[i,],
              nrow = length(subjects),
              ncol = ncol(mat),
              dimnames = list(subjects, colnames(mat)))
    }
  }

	if ("dominance"==model)
	{
		mat <- mat[,-(1:length(happy.get.strains(h)))]
	}

  if (as.data.frame)
  {
    mat <- as.data.frame(mat)
    if (1==ncol(mat))
    {
      colnames(mat) <- model
    }
    if (!is.null(subjects))
    {
        rownames(mat) <- subjects
    }
  }
  mat
}

happy.get.design <- function(h, marker,
        model="additive",
        subjects=NULL,
        as.data.frame=TRUE,
        sdp=NULL,
        merge.matrix=NULL)
{
  assert.happy(h)

  which.subjects=NULL
  if (!is.null(subjects))
  {
    subjects  <- as.character(subjects)
    if (!identical(subjects,happy.get.subjects(h)))
    {
      which.subjects <- match(subjects, happy.get.subjects(h))
    }
  }
  else
  {
    subjects = happy.get.subjects(h)
  }
  
  mat=NULL
  if (igrep("genotype", model)) # Genotype models
  {
    gmodel=ifow("genotype"==model, "factor", sub("genotype\\.*", "", model, perl=TRUE))
    mat=happy.get.genotype(h, marker=marker, genotype.model=gmodel)
    mat=as.matrix(mat)
    # subjects
    if (!is.null(which.subjects))
    {
      mat=mat[which.subjects,]
    }
    rownames(mat)=happy.get.subjects(h)
  }
  else if (!is.null(merge.matrix))
  {
    M            = merge.matrix
    num.groups   = ncol(M)
    if (is.null(colnames(M)))
    {
      colnames(M) = paste("merge",sep="",1:ncol(M))
    }
    if (1 >= num.groups | happy.num.strains(h) < num.groups)
    {
      stop("SDP or mergematrix must specify 2-", happy.num.strains(h), " groups\n")
    }
    num.subjects = length(subjects)
    group.names  = colnames(M)

    if ("additive"==model)
    {
      amat = as.matrix(happy.load.marker(h, marker=marker, model="additive"))
      if (!is.null(which.subjects))
      {
        amat=amat[which.subjects,]
      }
      mat = amat %*% M
      rownames(mat) = subjects
    }
    else if ("dominance"==model)
    {
      fmat = happy.get.design(h,
          model="full",
          marker=marker,
          merge.matrix=M,
          subjects=subjects,
          as.data.frame=FALSE)
      if (1==ncol(fmat)-num.groups)
      {
        mat = as.matrix(fmat[,-(1:num.groups)])
        colnames(mat) = colnames(fmat)[-(1:num.groups)]
      }
      else if (1==nrow(fmat))
      {
        mat = t(as.matrix(fmat[,-(1:num.groups)]))
      }
      else
      {
        mat = fmat[,-(1:num.groups)]
      }
    }
    else if (model %in% c("full","full.asymmetric"))
    {
      happy.model=ifow("full.asymmetric"==model, model, "full")

      # make tensor of diplotype matrices
      strains.tensor=happy.get.diplotype.tensor(h, 
          marker=marker, model=happy.model,
          subjects=subjects, memoize=TRUE)
      groups.tensor=array(numeric(0),
          dim=c(num.groups,num.groups,num.subjects),
          dimnames=list(group.names,group.names,subjects))

      # conversion loop
      Mt = t(M) # optimization
      for (i in 1:num.subjects)
      {
        groups.tensor[,,i] = Mt %*% strains.tensor[,,i] %*% M
      }
      
      # flatten to happy models
      if ("full.asymmetric"==model)
      {
        stop("Strain merge on full.asymmetric is not implemented yet\n")
      }
      else
      {
        row1=happy.matrixop.diplotypes.to.full(groups.tensor[,,1], symmetric.X=TRUE, want.names=TRUE)
        row1=t(as.matrix(row1))
        if (1==num.subjects)
        {
          mat=row1
        }
        else
        {
          mat=matrix(numeric(0),
              nrow=num.subjects,
              ncol=length(row1),
              dimnames=list(subjects,colnames(row1)))
          mat[1,]=row1
          for (i in 2:num.subjects)
          {
            mat[i,]=happy.matrixop.diplotypes.to.full(groups.tensor[,,i], 
                symmetric.X=TRUE,
                want.names=FALSE)
          }
        } 
      }
    }
    else
    {
      stop("Can't perform sdp on model '", model, "'\n") 
    }
  }
  else if (!is.null(sdp)) # Merged models
  {
    # error checking
    if (3>happy.num.strains(h))
    {
      stop("sdp requires at least 3 founders\n")
    }
    if (length(sdp)!=happy.num.strains(h))
    {
      stop("sdp must have as many elements as there are founders: expected ",
          happy.num.strains(h), ", got ",length(sdp), "\n")
    }
    # make merge matrix
    M = incidence.matrix(as.factor(sdp))
    mat=happy.get.design(h,
        marker=marker,
        subjects=subjects,
        merge.matrix=M,
        model=model,
        as.data.frame=FALSE)
  }
  else # Standard Happy models
  {
    if ("additive"==model)
    {
      mat <- as.matrix(happy.load.marker(h, marker=marker, model="additive"))
    }
    else if ("dominance"==model)
    {
      mat = happy.load.marker(h, marker=marker, model="full")
  		mat = mat[,-(1:length(happy.get.strains(h)))]
    }
  	else if ("full"==model)
  	{
      mat <- happy.load.marker(h, marker=marker, model="full")
  	}
  	else
  	{
  	  stop("Unknown model '", model, "'\n")
  	}
  	# subjects
  	if (!is.null(which.subjects))
  	{
      if (1==length(subjects))
      {
  	    mat=t(as.matrix(mat[which.subjects,]))
  	  }
  	  else
  	  {
  	    mat=mat[which.subjects,]
  	  }
  	}
	  rownames(mat)=subjects
  }
     
  # force to be array if 1 column
  if (is.null(dim(mat)))
	{
		mat <- as.array(mat)
	}

  # convert to data frame if requested
  if (as.data.frame)
  {
    mat <- as.data.frame(mat)
    if (1==ncol(mat))
    {
      colnames(mat) <- model
    }
  }
  mat
}


happy.get.genotype <- function(h, marker, genotype.model="factor")
{
  mat <- happy.load.marker(h, marker=marker, model="genotype")

  if ("factor"==genotype.model)
  {
    mat <- as.factor(mat)
  }
  else if ("full"==genotype.model)
  {
    mat = incidence.matrix(as.factor(mat))
  }
  else if (genotype.model %in% c("additive", "dominance", "hier") )
  {
    mat <- as.matrix(genotype.to.hier(as.vector(mat)))
  	if ("additive"==genotype.model)
  	{ 
  	  mat <- as.matrix(mat[,1])
  	  colnames(mat) = "additive"
  	}
  	else if ("dominance"==genotype.model)
  	{
  	  mat <- as.matrix(mat[,2])
  	  colnames(mat) = "dominance"
  	}
  }
	else
	{
		stop("Unknown model: ", genotype.model, "\n")
	}
	if (is.null(dim(mat)))
	{
		mat <- as.array(mat)
	}
	mat
}


happy.get.diplotype.tensor <- function(h, marker,  model, subjects=happy.get.subjects(h), simplify=FALSE, memoize=TRUE)
# returns a list of diplotype matrices
# TODO: allow caching of previously requested tensor
{
	assert.happy(h)
	if (! model %in% c("full", "full.asymmetric") )
	{
		stop("happy.get.diplotype.matrix() only implemented for full and full.asymmetric models\n")
	}
  subjects <- as.character(subjects)

  # simple memoizing: getter
  if (happy.reserve.has.scratch(h) & memoize)
  {
    if (happy.reserve.has(h, category="scratch", object.name="scratch.diplotype.tensor"))
    {
      scratch=happy.reserve.get(h, category="scratch", object.name="scratch.diplotype.tensor")
      if (all(scratch$subjects==subjects) & scratch$model==model & scratch$marker==marker)
      {
        return (scratch$tensor)
      }
    }
  }

  # create tensor
  x.mat <- happy.get.design(h,
          marker        = marker,
          model         = model,
          subjects      = subjects,
          as.data.frame = FALSE)
  strains      = happy.get.strains(h)
  num.strains  = length(strains)
  num.subjects = length(subjects)
  mat.tensor   = array(numeric(0),
      dim=c(num.strains, num.strains, length(subjects)),
      dimnames=list(strains, strains, subjects))
  for (i in 1:num.subjects)
  {
  	if ("full"==model)
  	{
      mat.tensor[,,i] = happy.matrixop.full.to.diplotypes(x.mat[i,], num.strains)
  	}
  	else if ("full.asymmetric"==model)
  	{
  		mat.tensor[,,i] <- happy.matrixop.full.asymmetric.diplotypes(x.mat[i,], num.strains)
  	}
  }
  
  # simple memoizing: setter
  if (happy.reserve.has.scratch(h) & memoize)
  {
    scratch=list(tensor=mat.tensor, marker=marker, model=model, subjects=subjects)
    happy.reserve.put(h, category="scratch", object.name="scratch.diplotype.tensor", object=scratch)
  }
  
  # reduce if a single individual and simplifying (no memoizing)
  if (simplify & 1==length(subjects))
  {
      return (mat.tensor[,,1])
  }
  mat.tensor
}

happy.matrixop.full.to.diplotypes <- function(x, num.strains)
{
  m <- matrix(0, ncol=num.strains, nrow=num.strains)
	diag(m) <- x[1:num.strains]
	m[upper.tri(m, diag=FALSE)] = x[(num.strains+1):length(x)]
	0.5 * (m + t(m))
}

happy.matrixop.full.asymmetric.to.diplotypes <- function(x, num.strains)
{
  matrix(x, nrow=num.strains)
}

happy.matrixop.diplotypes.to.full <- function(X, symmetric.X=FALSE, want.names=TRUE)
{
  if (!symmetric.X)
  {
    X=0.5*(X+t(X))
  }
  f = c(diag(X), 2*X[upper.tri(X, diag=FALSE)])
  if (want.names & !is.null(colnames(X)))
  {
    A=matrix(kronecker(colnames(X), colnames(X), FUN=paste, sep="."),
        nrow=ncol(X), byrow=TRUE)
    names(f)=c(diag(A), A[upper.tri(A, diag=FALSE)])
  }
  f
}

happy.get.first.marker <- function(h, chromosome=NULL)
{
    if (!is.null(chromosome))
    {
        x <- character(length(chromosome))
        for (i in 1:length(chromosome))
        {
            x[i] <- happy.get.markers(h, chromosome=chromosome[i])[1]
        }
        return (x)
    }
    else
    {
        return (happy.get.markers(h)[1])
    }
}

happy.get.interval.midpoint <- function(h, markers, scale="bp", fudge.bp=FALSE)
# return the midpoint of the interval
{
    p <- happy.get.location(h, markers, scale=scale)
    p + happy.get.interval.length(h, markers, scale=scale, fudge.bp=fudge.bp)/2
}

happy.get.interval.over <- function(h, chromosome, x,
        scale                = "cM",
        use.nearest.terminus = FALSE,
        boundary.choice      = "l",
        fudge.bp             = FALSE)
# return which intervals are over the specified locations
# x may be a vector
# boundary.choice == l | r | NA
{
  assert.happy(h)
  chromosome  <- rep(chromosome, length.out=length(x))
  boundary.choice <- rep(boundary.choice, length.out=length(x))

  markers <- happy.get.markers(h, chromosome=chromosome)
  chrom   <- happy.get.chromosome(h, markers)
  range   <- happy.get.interval.range(h, markers, scale=scale, fudge.bp=fudge.bp)

  overlap.marker <- rep(NA, length(x))
  for (i in 1:length(x))
  {
      my.chrom <- chromosome[i]
      my.loc   <- x[i]
      overlap.idx <- which( my.chrom==chrom
              & my.loc >= range[,1]
              & my.loc <= range[,2])

      if (2==length(overlap.idx))
      {
          if (is.na(boundary.choice[i]))
          {
              overlap.marker[i] <- NA
          }
          else if ("l"==boundary.choice[i])
          {
              overlap.marker[i] <- markers[overlap.idx[1]]
          }
          else
          {
              overlap.marker[i] <- markers[overlap.idx[2]]
          }
      }
      if (1==length(overlap.idx))
      {
          overlap.marker[i] <- markers[overlap.idx]
      }
      if (0==length(overlap.idx) & use.nearest.terminus)
      {
          is.early <- my.loc < happy.get.location(h, scale=scale,
                  happy.get.first.marker(h, chromosome=my.chrom))
          overlap.marker[i] <- ifelse(is.early,
                  happy.get.first.marker(h, chromosome=my.chrom),
                  happy.get.last.marker(h, chromosome=my.chrom))
      }
  }
  overlap.marker
}

happy.get.interval.range <- function(h, markers, scale="cM", fudge.bp=FALSE)
# get start bp and end bp of requested intervals
{
    r <- happy.get.location(h, markers, scale=scale)
    r <- cbind(r, r + happy.get.interval.length(h, markers, scale=scale, fudge.bp=fudge.bp))
    if ("bp"==scale | "Mb"==scale)
    {
        one.base <- ifelse("Mb"==scale, 1e-6, 1)
        r[,2] <- r[,2]-one.base
    }
    rownames(r) <- markers
    colnames(r) <- c("begin","end")
    return (r)
}


happy.get.intervals <- function(h, chromosome=NULL)
{
    happy.get.markers(h, chromosome=chromosome, as.intervals=TRUE)
}

happy.get.intervals.in.range <- function(h,
        from       = NULL,
        to         = NULL,
        markers    = NULL,
        chromosome = NULL,
        scale      = "interval")
{
    ## deal with requests for all markers or all chromosome markers...
    if (is.null(from) & is.null(to))
    {
        return ( happy.get.markers(h, chromosome=chromosome) )
    }

    ## calculate range

    if (is.null(chromosome) & "interval"!=scale)
    {
        stop("Must specify chromosome= if using ", scale, "\n")
    }

    # specify start of range
    marker1 <- NULL
    if (is.null(from))
    {
        marker1 <- happy.get.first.marker(h, chromosome=chromosome)
    }
    else
    {
        if ("interval"==scale)
        {
            if (!happy.has.markers(h, from))
            {
                stop("Could not find marker ", from, "\n")
            }
            marker1 <- from
        }
        else
        {
            marker1 <- happy.get.interval.over(h,
                    chromosome=chromosome,
                    x=from,
                    scale=scale,
                    use.nearest.terminus=TRUE)
        }
    }

    # specify end of range
    marker2 <- NULL
    if (is.null(from))
    {
        marker2 <- happy.get.last.marker(h, chromosome=chromosome)
    }
    else
    {
        if ("interval"==scale)
        {
            if (!happy.has.markers(h, to))
            {
                stop("Could not find marker ", to, "\n")
            }
            marker2 <- to
        }
        else
        {
            marker2 <- happy.get.interval.over(h,
                    chromosome=chromosome,
                    x=to,
                    scale=scale,
                    use.nearest.terminus=TRUE)
        }
    }
    happy.get.markers.between(h, from=marker1, to=marker2)
}


happy.get.last.marker <- function(h, chromosome=NULL, as.intervals=TRUE)
{
    m <- happy.get.markers(h, chromosome=chromosome, as.intervals=as.intervals)
    m[length(m)]
}

happy.get.location <- function(h,markers, scale="bp")
{
    switch(scale,
            bp = happy.get.bp(h, markers),
            Mb = happy.get.bp(h, markers)/1e6,
            cM = happy.get.position(h, markers))
}

happy.get.markers.between <- function(h, to=NULL, from=NULL, before=NULL,
        after=NULL,
        as.intervals=TRUE)
# return all markers between two specified markers
{
    if (1<length(to) | 1<length(from) | 1<length(before) | 1<length(after))
    {
        stop("Arguments must be of length 1\n")
    }

    markers <- happy.get.markers(h, as.intervals=as.intervals)

    start <- NA
    if (!is.null(after))
    {
        start <- which(after==markers) + 1
    }
    else if (!is.null(from))
    {
        start <- which(from==markers)
    }

    end <- NA
    if (!is.null(before))
    {
        end <- which(before==markers) - 1
    }
    else if (!is.null(to))
    {
        end <- which(to==markers)
    }
    if (!force.logical(start) | !force.logical(end))
    {
        stop("Could not find start and end points for markers from=",
                from,", to=",to,", before=",before,", after=",after,"\n" )
    }
    markers[start:end]
}


happy.get.next.marker <- function(h, markers, as.intervals=TRUE, within.chr=FALSE)
# TODO: fix the fact that chr 10 right after chr 1
{
    found <- happy.has.markers(h, markers)
    if (!all(found))
    {
        stop("No such markers: ", paste(markers[!found], collapse=", "), "\n")
    }
    if (within.chr)
    {
        out <- character(length(markers))
        for (i in 1:length(markers))
        {
            chr.markers <- happy.get.markers(h,
                    as.intervals=as.intervals,
                    chr=happy.get.chromosome(h, markers[i]))
            mi <- match(markers[i], chr.markers)
            out[i] <- chr.markers[mi+1]
        }
        return (out)
    }
    else
    {
        all.markers <- happy.get.markers(h, as.intervals=as.intervals)
        mi <- match(markers, all.markers)
        return (all.markers[mi+1])
    }
}


happy.get.previous.marker <- function(h, marker, as.intervals=TRUE)
{
    markers <- happy.get.markers(h, as.intervals=as.intervals)
    i <- match(marker, markers)
    if (any(is.na(i)))
    {
        stop("No such markers: ", paste(marker[which(is.na(i))], collapse=", "), "\n")
    }
    if (any(1==i))
    {
        stop("No marker previous to ", markers[1==i], "\n")
    }
    markers[i-1]
}


happy.has.chromosomes <- function(h, chroms, model="genotype")
{
    chroms %in% happy.list.chromosomes(h, model=model)
}

happy.has.subjects <- function(h, subjects)
{
    subjects %in% happy.get.subjects(h)
}


happy.has.markers <- function(h, markers, model="additive")
{
    markers %in% happy.get.markers(h, model=model)
}


happy.list.chromosomes <- function(h, sort=TRUE, model="genotype")
{
    assert.happy(h)
    chr <- unique(as.character(h[[model]]$genome$chromosome))
    if (sort)
    {
        ints  <- suppressWarnings(as.integer(chr))
        chars <- chr[is.na(ints)]
        ints  <- ints[!is.na(ints)]
        chr   <- c(as.character(sort(ints)), sort(chars))
    }
    chr
}

happy.make.colnames <- function(strain.names, model)
{
	if (!is.character(strain.names))
	{
		stop("Must pass strain.names as character vector to happy.make.colnames()\n")
	}
	
	num.strains <- length(strain.names)
	
	if ("additive"==model)
	{
		return (strain.names)
	}
	if ("genotype"==model)
	{
		return (NULL)
	}
	
	diplotype.names <- matrix(
			kronecker(strain.names, strain.names, paste, sep = "."),
			nrow = num.strains)
	if ("full"==model)
	{
		return ( c(diag(diplotype.names),
				diplotype.names[upper.tri(diplotype.names, diag = FALSE)])
				)
	}
	if ("full.asymmetric"==model)
	# assumes row major order, ie, (row1, row2, etc), in C object
	{
		return (c(t(diplotype.names)))
	}
	else
	{
		stop("No colnames defined for model ", model, "\n")
	}
}

happy.num.strains<-function(h)
{
	length(happy.get.strains(h))
}
```
