## Supplementary material for "Determinants of QTL mapping power in the realized Collaborative Cross": SPARCC R package used for all analyses: read.happy.genotype.R

```
#### Written by Will Valdar - from bagpipe.backend package

genotype.to.factor <- function(g)
{
    unphased   <- rep(NA, length=nrow(g))

    ok      <- complete.cases(g)
    ordered <- g[,1] <= g[,2]

    mask <- ordered & ok
    unphased[mask] <- paste(g[mask,1], g[mask,2], sep="_")

    mask <- !ordered & ok
    unphased[mask] <- paste(g[mask,1], g[mask,2], sep="_")

    as.factor(unphased)
}

genotype.to.count <- function(g)
{
    unique.g <- unique(c(na.omit(as.character(g))))
	if (any(2!=nchar(unique.g)))
	{
		stop("Genotypes must be 2 characters long or NA\n")
	}

    alleles  <- unique(unlist(strsplit(unique.g, "")))
    if (2 < length(alleles))
    {
        stop("Cannot interpret genotype as additive with than >2 alleles\n")
    }

    count <- rep(NA, length(g))
    count[g==paste(alleles[1],alleles[1],sep="")] <- 0

    if (2==length(alleles))
    {
         count[g==paste(alleles[2], alleles[1], sep="")] <- 1
         count[g==paste(alleles[1], alleles[2], sep="")] <- 1
         count[g==paste(alleles[2], alleles[2], sep="")] <- 2
    }
    return (count)
}

genotype.to.hier <- function(g)
{
    g   <- genotype.to.count(g)

    lo  <- g==0
    het <- g==1
    hi  <- g==2

    # use parameterization of Cordell

    # x describes additive contribution only
    x <- rep(NA, length(g))
    x[lo]  <- -1
    x[het] <- 0
    x[hi]  <- 1

    # z describes dominance contribution only
    z <- rep(NA, length(g))
    z[lo]  <- -0.5
    z[het] <- 1
    z[hi]  <- -0.5

    return (data.frame(additive=x, dominance=z))
}
```
