## Supplementary material for "Determinants of QTL mapping power in the realized Collaborative Cross": SPARCC R package used for all analyses: read.happy.reserve.R

```
#### Written by Will Valdar - from bagpipe.backend package

happy.init.reserve <- function(h,
    memory.limit.Mb=Inf,
    models=happy.get.models(h),
    auto.reserve=TRUE,
    allow.scratch=TRUE)
{
	h$DATA <- list()
	if (allow.scratch)
	{
	  models=c(models, "scratch")
	}
	for (m in models)
	{
		h$DATA[[m]] <- new.hash()
	}
	h$DATA.MAX.MEMORY <- memory.limit.Mb*2^20
	h$DATA.AUTO.ADD	<- auto.reserve
	h
}

happy.clear.reserve <- function(h)
{
  h$DATA=NULL
  h$DATA.MAX.MEMORY=NULL
  h$DATA.AUTO.ADD=NULL
  h
}

happy.is.auto.reserve <- function(h)
{
	if (is.null(h$DATA.AUTO.ADD)) return (FALSE)
	h$DATA.AUTO.ADD
}

happy.reserve.has.scratch <- function(h)
{
  !is.null(h$DATA[["scratch",exact=TRUE]])
}

happy.get.reserve.limit <- function(h)
{
	h$DATA.MAX.MEMORY
}

happy.set.auto.reserve <- function(h, bool)
{
	h$DATA.AUTO.ADD <- bool
	h
}

#---CORE RESERVE ACCESS FUNCTIONS---

happy.reserve.exists <- function(h)
{
	!is.null(h$DATA)
}

happy.reserve.memory.usage <- function(h)
{
  if (!happy.reserve.exists(h))
  {
    stop("Reserve does not exist!\n")
	}
  sum(sapply(h$DATA, hash.memory.usage))
}

happy.reserve.get <- function(h, category, object.name)
{
  hash.get(h$DATA[[category, exact=TRUE]], object.name)
}

happy.reserve.has <- function(h, category, object.name=NULL)
{
  if (!happy.reserve.exists(h))
  {
    return(FALSE)
  }
  if (is.null(h[["DATA", exact=TRUE]][[category, exact=TRUE]]))
  {
    return (FALSE)
  }
  if (is.null(object.name))
  {
    return (TRUE)
  }
  hash.has(h$DATA[[category, exact=TRUE]], object.name)
}

happy.reserve.put <- function(h, category, object.name, object)
{
  if (!happy.reserve.exists(h))
  {
    stop("Cannot reserve object because reserve is not initialized\n")
  }
  if (is.null(h$DATA[[category, exact=TRUE]]))
  {
    stop("Cannot reserve object because category ", category, " does not exist\n")
  }
  hash.put(h$DATA[[category, exact=TRUE]], object.name, object)
}
```
