## Supplementary material for "Determinants of QTL mapping power in the realized Collaborative Cross": SPARCC R package used for all analyses: WVmisc.R

```
#### Written by Will Valdar - from bagpipe.backend package

caught.error <- function(x)
# simple catch for try()
{
    inherits(x, "try-error")
}

dfapply <- function(data, INDICES, FUN, results=list(), results.add.FUN=c, matched.vector=FALSE, pass.key=is.list(INDICES), ...)
# Function for processing each cell of a ragged array when the array is a data frame. An alternative to by()
{ 
  idata <- as.data.frame(INDICES)
  udata <- unique(idata)
  if (matched.vector)
  {
    results <- rep(NA, nrow(data))
  }
  for (ui in 1:nrow(udata))
  {
    di <- rep(TRUE, nrow(data))
    for (cn in colnames(udata))
    {
      di <- di & as.character(udata[ui,cn])==as.character(idata[,cn])
    }
    u <- udata[ui,]
    if (!is.data.frame(u)) {
      u <- data.frame(u)
      colnames(u) = colnames(udata)
    }
    if (pass.key) {
      result <- FUN(data[di,], u, ...)
    } else { # allow for really simple operations
      result <- FUN(data[di,], ...)
    }
    
    if (matched.vector){
      results[which(di)] <- result
    } else {
      results <- results.add.FUN(results, result)
    }
  }
  results
}


elem <- function(x, start=1, end=length(x))
{
  ifow(0==length(x), integer(0), start:end)
}

force.logical <- function(x, null=FALSE, na=FALSE, empty=FALSE, blank=FALSE)
# 
#
# Problem definition: Rs "if" statements evaluate all conditions whether
# or not it is necessary to do so. Eg,
#      x <- NA
#      ...
#      if (!is.na(x) & x > 0)
# The above condition will evaluate as (FALSE & NA) -> (NA) -> throw error,
# whereas in other languages the second component is not evaluated if the
# first is FALSE and no error would be thrown. The default solution to this
# is to use multiple nested if statments like
#      if (!is.na(x)) { if (x>0) {
# However, that is expensive on nesting and brackets.
# A better alternative:
#      if (!is.na(x) & force.logical(x>0))
#  or more explicitly
#      if (!is.na(x) & force.logical(x>0, na=FALSE))
#  or, for this example, most concisely
#      if (force.logical(x>0))
# Whichever, this means the condition requires only one if statement.
{
    if (is.null(x))
    {
        return (null)
    }
    if (0==length(x))
    {
        return (empty)
    }
    if (any(is.na(x)))
    {
        x[is.na(x)] <- na
    }
    if (is.character(x))
    {
        i <- nchar(x)==0
        x[i] <- FALSE
        x[!i] <- TRUE
    }
    as.logical(x)
}

formula.as.string <- function(x)
# Converts a formula object into a character string
# If x is a list of formula objects, a character vector of the same
# length is returned. If x is a string or vector of strings, then x is
# returned unchanged.
{
    if (is.character(x))
    {
        return (x)
    }
    if (is.formula(x))
    {
        return (paste(deparse(x), collapse=""))
    }
    if (is.list(x))
    {
        strings <- NULL
        for (i in 1:length(x))
        {
            strings <- c(strings, formula.as.string(x[[i]]))
        }
        return (strings)
    }
    stop("Cannot convert object of class ", class(x), " to string\n")
}

incidence.matrix <- function(fact)
{
    m=diag(nlevels(fact))[fact,]
    colnames(m)=levels(fact)
    m
}


igrep <- function(pattern, x, ..., value=FALSE, logical=TRUE)
# pass through method for grep that can return match as an indicator
# vector
{
    if (!value & logical)
    {
        indices <- grep(pattern, x, value=value, ...)
        return (1:length(x) %in% indices)
    }
    grep(pattern, x, value=value, ...)
}

interpolate <- function(x, y, xout, project=1, ...)
# wrapper for approx() that linearly projects missing data
# at the front and back. $project determines how many
# terminal non-missing points are used to calculate a
# a projection slope.
{
	yout <- approx(x, y, xout=xout, ...)$y
	
	na <- is.na(yout)
	if (0<project & any(na))
	{
		known <- which(!na)
		if (na[1])
		{
			k     <- known[1]
			front <- 1:(k-1)
			loc   <- yout[k]
			slope <- (yout[k+project]-loc)/project
			yout[front] <- loc + (front - k)*slope
		}
		if (tail(na,1))
		{
			k     <- tail(known,1)
			back  <- length(yout):k
			loc   <- yout[k]
			slope <- (loc - yout[k-project])/project
			yout[back] <- loc + (back - k)*slope
		}
	}
	list(x=xout, y=yout)
}

interpolate.Sys.env <- function(x, stop.on.fail=FALSE)
# interpolates environmental shell variables
# ie, finds each substring x that start with "$" and ends before a "/" or whitespace,
# and substitutes it with Sys.getenv(x). If Sys.getenv(x) returns an empty string
# such that there is no environmental variable with that name, then if stop.on.fail
# is TRUE the function throws an error. Otherwise, the substring is left unsubstituted
{
	if (!is.character(x))
	{
		stop("Argument to interpolate.Sys.env currently handles only character vectors\n")
	}
	if (1!=length(x))
	{
		return ( as.character(sapply(x, interpolate.Sys.env, stop.on.fail=stop.on.fail)) )
	}
	regmatch <- gregexpr("\\$[^\\/]+", x, perl=TRUE)[[1]]
	starts <- as.integer(regmatch)
	lens   <- attr(regmatch, "match.length")
	if (-1==starts[1]) return (x)

	parts <- NULL
	next.pos <- NULL
	for (i in 1:length(starts))
	{
		if (!is.null(next.pos))
		{
			parts <- c(parts, substr(x, next.pos, starts[i]-1) )
		}

		shellvar <- substr(x, starts[i]+1, lens[i] + starts[i] - 1)
		value <- Sys.getenv(shellvar)
		if (""==value)
		{	
			if (stop.on.fail) stop("Could not get shell environment variable \"$", shellvar, "\"\n")
			value <- paste("$", sep="", shellvar)
		}
		parts <- c(parts, value)
		
		next.pos <- starts[i] + lens[i]
	}
	parts <- c(parts, substr(x, next.pos, nchar(x)))	
	paste(parts, collapse="")
}
ENV <- function(...) { interpolate.Sys.env(...) }

ifow=function(test, yes, no)
{
  if (test)
  {
    return (yes)
  }
  no
}

invlogit <- function(x)
{
    exp(x)/(1+exp(x))
}

is.formula <- function(x)
# for some reason absent in R base
{
  inherits(x, "formula")
}

is.wholenumber <- function(x, tol = .Machine$double.eps^0.5)
{
  abs(x - round(x)) < tol
}

list.subdirs <- function(path=".", ..., full.names=FALSE)
{
  path.names <- list.files(path, ..., full.names=TRUE)
	files      <- list.files(path, ..., full.names=full.names)
  files[ file.info(path.names)$isdir ]
}

logit <- function(p)
{
  log(p/(1-p))
}

map.eq <- function(x, lookup=NULL, from=NULL, to=NULL)
# maps values in vector from $from to $to using == to test equality
{
    y <- x
    if (!is.null(from) & !is.null(to))
    {
		if (length(from)!=length(to))
		{
			stop("Arguments $from and $to must be the same length\n")
		}
        for (k in 1:length(from))
        {
            i <- x==from[k]
            y[i] <- to[k]
        }
    }
	if (!is.null(lookup))
	{
		y <- map.eq(y, from=names(lookup), to=unlist(lookup))
	}
    return (y)
}

object.sizes <- function(env=parent.frame(), sort="sd", format="Mb")
{
	obs <- objects(name=env)
	sizes <- numeric(length(obs))
	for (i in 1:length(obs))
	{
		sizes[i] <- eval(parse(text=paste("object.size(",obs[i], ")")), env=env)
	}
	names(sizes) <- obs
	
	if ("sd"==sort)
	{
		sizes <- sizes[order(-sizes)]
	}
	if ("Mb"==format)
	{
		sizes <- round(sizes/2^20, 3)
	}
	sizes   
}

se.mean <- function(x, na.rm=FALSE) # std error of the mean
{
    n <- sum(!is.na(x))
    if (n < 2) return (NaN)
    sd(x, na.rm=na.rm)/sqrt(n)
}

split.formula <- function(x, simplify=FALSE)
{
  terms.object = terms(as.formula(x), simplify=simplify)
  formula.env  = attr(terms.object, ".Environment")
  formula      = paste(string.trim(deparse(terms.object)), collapse="")

  vars <- all.vars(terms.object)
  facts <- rownames(attr(terms.object, "factors"))

  has.response   <- 1==attr(terms.object, "response")
  has.predictors <- !is.null(facts)
  has.intercept  <- 1==attr(terms.object, "intercept")

  response <- ""
  response.vars <- NULL
  if (has.response)
  {
    response = string.trim(sub("~.*", "", formula))
    response.terms.object = terms(
        as.formula(paste(response,"~1"), env=formula.env)
        )
    num.responses = length(all.vars(response.terms.object))
    response.vars = vars[1:num.responses]
  }
    
  predictor.string <- ""
  predictors       <- NULL
  predictor.vars   <- NULL
  if (has.predictors)
  {
    predictor.string = string.trim(sub(".*~", "", formula))
    predictors.terms.object = terms(
        as.formula(paste("~",predictor.string), env=formula.env)
        )
    predictors     = rownames(attr(predictors.terms.object, "factors"))
    predictor.vars = all.vars(predictors.terms.object)
  }
  else
  {
    predictor.string = ifelse(has.intercept, "1", "-1")
  }

  list(
      formula          = formula,
      response         = response,
      response.vars    = response.vars,
      predictor.string = predictor.string, # formerly `predictors'
      predictors       = predictors,
      predictor.vars   = predictor.vars
      )
}

split.pathname <- function(x)
{
    list(
            base = basename(x),
            dir  = dirname(x),
            ext  = sub(".*\\.", "", x),
            core = sub("\\.[^\\.]+", "", basename(x)))
}

strcat<-function(...,sep=""){paste(sep=sep,...)}

string.trim <- function(s)
# trims leading and trailing whitespace from a character vector
{
    gsub("^[[:space:]]+", "", gsub("[[:space:]]+$", "", s) )
}

tr <- function(mat)
{
	sum(diag(mat))
}

write.delim <- function(..., quote=FALSE, row.names=FALSE, sep="\t")
{
    write.table(..., sep=sep, quote=quote, row.names=row.names)
}
```
