## Supplementary material for "Determinants of QTL mapping power in the realized Collaborative Cross": SPARCC R package used for all analyses: README.md

sparcc
====

**S**imulated **P**ower **A**nalysis in the **R**ealized **C**ollaborative **C**ross

This package provides two primary functions:

1. Simulate Collaborative Cross (CC) phenotype data from the actual CC lines
- Potential parameters:
  - QTL effect size
  - Overall strain effect size
  - Number of CC lines
  - Number of replicates
  - Number of functional alleles at the QTL
  - Strain distribution pattern of the functional alleles
2. Estimate power based on rapid genome scans of simulated data 
- Genome scans are highly optimized by storing QR decompositions when appropriate
- Signicance thresholds determined through rapid permutations
- Additional statistics of interest possible, such as false positive rate

Finally, sparcc also includes functionality to visualize results. It can also be extended to other recombinant inbred line (RIL) panels.
