## Supplementary material for "Determinants of QTL mapping power in the realized Collaborative Cross": Overview of data and Supplement

### Data and Supplement details

Data and analysis files in the **Supplement** include:

- File_S1: data_supplement_details.pdf - Overview of data and **Supplement**.
- File_S2: sparcc_1.1.1.tar.gz - SPARCC R package used for all analyses.
- File_S3: sparcc_cache.zip - CC haplotype mosaics formatted for SPARCC.
- File_S4: sparcc_powersim.R - Example R script to perform large-scale power analysis.
- File_S5: sparcc_powersim.sh - Example shell script to coordinate separate calls to sparcc_powersim.R.
- File_S6: collapse_results.R - Example R script to aggregate results from multiple calls to sparcc_powersim.R.
- File_S7: sparcc_options_tutorial.pdf - Provides description of options in SPARCC, and provides simple tutorials for using the package.
- File_S8: generate_figures.R - R script to generate figures in manuscript and **Supplement**.
- File_S9: supplement_tables_figures.pdf - The supplementary tables and figures.

#### *SPARCC Package*

The static version of the SPARCC R package (1.1.1) used in this paper is provided in the **Supplement** as File_S2. R and shell scripts are provided to generate the same results as File_S4, File_S5, File_S6, and File_S8. Static versions of these files have been placed on figshare. The current version of the SPARCC package is available here: https://github.com/gkeele/sparcc. SPARCC can be installed using the command 'R CMD INSTALL' at the terminal. The current version can be conveniently installed using the devtools R package and the following command within R: 'install_github("gkeele/sparcc")'.

#### *Data objects included in SPARCC package*

- r1.dat: Data frame of power and FPR results for Definition B from 1,000 simulations per combination of
  - Number of strains: [(10-70 by 5), 72]
  - Single observation per strain
  - Number of alleles: [2, 3, 8]
  - $h^2_{\text{strain}} = 0$
  - $h^2_{\text{QTL}}$: [0.01, (0.05-0.95 by 0.05)]
- r1.damb.dat: Data frame of power and FPR results for Definition DAMB from 1,000 simulations per combination of
  - Number of strains: [(10-70 by 5), 72]
  - Single observation per strain
  - Number of alleles: [2, 3, 8]
  - $h^2_{\text{strain}} = 0$
  - $h^2_{\text{QTL}}$: [0.01, (0.05-0.95 by 0.05)]
- r1.4v4.dat: Data frame of power and FPR results for Definition B with bi-allelic series forced to be balanced (4 founders per functional allele) from 1,000 simulations per combination of
  - Number of strains: [(10-70 by 5), 72]
  - Single observation per strain
  - Number of alleles: 2
  - $h^2_{\text{strain}} = 0$
  - $h^2_{\text{QTL}} = 0.5$
- r1.7v1.dat: Data frame of power and FPR results for Definition B with bi-allelic series forced to be imbalanced (singler founder with one allele) from 1,000 simulations per combination of
  - Number of strains: [(10-70 by 5), 72]
  - Single observation per strain
  - Number of alleles: 2
  - $h^2_{\text{strain}} = 0$
  - $h^2_{\text{QTL}} = 0.5$
- r1.dist.dat: Data frame of mean location error results for Definition B from 1,000 simulations per combination of
  - Number of strains: [(10-70 by 5), 72]
  - Single observation per strain
  - Number of alleles: [2, 3, 8]
  - $h^2_{\text{strain}} = 0$
  - $h^2_{\text{QTL}}$: [0.01, (0.05-0.95 by 0.05)]
- r1.exchange.dat: Data frame of FPR results for Definition B from 10,000 null simulations with correlated strain effects based on **K** per combination of
  - Number of strains: [71 (excluding CC059), 72]
  - Single observation per strain
  - Number of alleles: 0
  - $h^2_{\text{strain}}$: [0-1 by 0.2]
  - $h^2_{\text{QTL}} = 0$
- r1.beavis.dat: Data frame of QTL effect size estimations for Definition DAMB from 1,000 simulations per combination of
  - Number of strains: [(40-60 by 10), 72]
  - Single observation per strain
  - Number of alleles: 2
  - $h^2_{\text{strain}} = 0$
  - $h^2_{\text{QTL}}$: [(20-70 by 10)]
- K: Matrix of dimension $72 \times 72$ representing the realized genetic relationship matrix of the 72 CC strains, calculated from the founder mosaics.

#### *File types*

- *.R - These are R scripts used for the analyses and figures.
- *.sh - These are bash scripts used to run large-scale analysis.
- *.RData - These files are contained in the happy_formated sparcc_cache directory that can be loaded in R and are necessary for SPARCC to run.
