## Supplementary material for "Determinants of QTL mapping power in the realized Collaborative Cross": Example R script to aggregate results from multiple calls to sparcc_powersim.R: collapse_results.R

```
directory <- "./results/"
#n.strains <- c(seq(10, 70, 5), 72)
n.strains <- 72
n.replicates <- 1
#n.alleles <- c(2,3,8)
n.alleles <- 8
#h.qtl <- c(0.01, seq(0.05, 0.95, 0.05))
h.qtl <- 0.5
h.strain <- 0
n.jobs <- 1:10

collapse.results <- function(directory, n.strains, n.replicates, n.alleles, h.qtl, h.strain, n.jobs){
  #function to collapse results and report missing jobs
  missing <- c()
  full.results <- matrix(NA, length(n.strains)*length(n.replicates)*length(n.alleles)*length(h.qtl)*length(h.strain)*length(n.jobs), 9)
  row.index <- 0
  
  for (i in n.strains){
    for (j in n.replicates){
      for (k in n.alleles){
        for (l in h.qtl){
          for (m in h.strain){
            for (n in n.jobs){
              row.index <- row.index + 1
              
              job.id <- paste("powersim_n.strains_", i, "_n.reps_", j, "_n.alleles_", k, "_h.qtl_", format(l, nsmall=2), "_h.strain_", m, "_job_", n, sep="")
              job.path <- paste(directory, job.id, ".RData", sep="")
              job.call <- paste("bsub Rscript sparcc_powersim.R", job.id, "./CC_genomecache_l2norm0.1_build37/", i, j, 100, k, format(l, nsmall=2), m, "NA", sep=" ")

              #store parameters for job
              full.results[row.index,1:6] <- c(i,j,k,l,m,n)
              
              if (file.exists(job.path)){
                #store results for job
                load(job.path)
                full.results[row.index,7] <- mean(results$results)
                full.results[row.index,8] <- mean(results$results.window)
                full.results[row.index,9] <- mean(results$results.false)
              } else {
                #record if job is missing
                missing <- c(missing, job.call)
              }
            }
          }
        }
      }
    }
  }
  
  #convert results to data frame and collapse
  full.results <- data.frame(full.results)
  names(full.results) <- c("n.strains", "n.replicates", "n.alleles", "h.qtl", "h.strain", "n.jobs", "power", "power.window", "false")
  
  full.results <- aggregate(full.results, by=list(full.results$n.strains, full.results$n.replicates, full.results$n.alleles, full.results$h.qtl, full.results$h.strain), mean)
  full.results <- full.results[-c(1:5, 11)]
  
  #return missing list and collapsed results
  list("missing"=missing, "full.results"=full.results)
}


full.results <- collapse.results(directory, n.strains, n.replicates, n.alleles, h.qtl, h.strain, n.jobs)
full.results$full.results$h.qtl <- round(full.results$full.results$h.qtl, 2)

save(full.results, file="full_results.RData")
```
