## Supplementary material for "Determinants of QTL mapping power in the realized Collaborative Cross": Example R script to perform large-scale power analysis: sparcc_powersim.R

```
args <- commandArgs(TRUE)

job.id <- as.character(args[1])
genomecache <- as.character(args[2])
n.strains <- as.numeric(args[3])
n.replicates <- as.numeric(args[4])
n.sims <- as.numeric(args[5])
n.alleles <- as.numeric(args[6])
h.qtl <- as.numeric(args[7])
h.strain <- as.numeric(args[8])
seed <- suppressWarnings(as.numeric(args[9]))

#job.id <- "powercurve_n.strains_72_n.reps_50_n.alleles_8_h.qtl_0.5_h.strain_0_job_10"
#genomecache <- "../SPARCC/CC_genomecache_l2norm0.1_build37/"
#n.strains <- 72
#n.replicates <- 10
#n.sims <- 5
#n.alleles <- 3
#h.qtl <- 0.1
#h.strain <- 0
#seed <- NA

########################################################

#load packages
#devtools::install_github("gkeele/miqtl")
#devtools::install_github("gkeele/sparcc")
library(miqtl)
library(sparcc)

#set seed
if (is.na(seed)){
  seed <- ceiling(runif(1)*10^9)
}
set.seed(seed, kind = "Mersenne-Twister", normal.kind = "Inversion")
seed

#object to store results
results <- rep(NA, n.sims)
results.false <- rep(NA, n.sims)
results.window <- rep(NA, n.sims)
gev.flag <- rep(NA, n.sims)
peak.dist <- rep(NA, n.sims)

#iterate simulation
for (i in 1:n.sims){
  
  if (i%%5 == 0){
    print(i)
  }
  
  #simulate dataset
  simulated.data <- sim.CC.data(genomecache = genomecache,
                                num.lines = n.strains, 
                                num.replicates = n.replicates,
                                num.sim = 1,
                                num.alleles = n.alleles,
                                qtl.effect.size=h.qtl, 
                                strain.effect.size=h.strain)
  
  #perform QR decomposition
  compact.CC.sample.qr <- extract.compact.qr(genomecache = genomecache, 
                                             CC.lines.matrix = as.matrix(simulated.data$data$SUBJECT.NAME.1),
                                             use.progress.bar = F)
  
  #scan
  simulated.scan <-  run.sim.scans(sim.data = simulated.data,
                                   use.progress.bar = F,
                                   return.all.sim.qr = F,
                                   all.sim.qr = compact.CC.sample.qr)
  
  #permutation thresholds using GEV
  permutation.index <- generate.perm.matrix(num.lines = n.strains, 
                                                        num.perm=100)
  
  simulated.threshold.scans <- run.perm.scans(perm.matrix = permutation.index,
                                                               sim.CC.scans = simulated.scan,
                                                               sim.CC.object = simulated.data,
                                                               all.sim.qr = pull.qr.from.compact(compact.qr.list=compact.CC.sample.qr,
                                                                                                 qr.index=1),
                                                               phenotype.index = 1,
                                                               use.progress.bar = F)
  
  simulated.threshold <- tryCatch(get.thresholds(simulated.threshold.scans), error=function(e){NULL})
  
  if (!is.null(simulated.threshold)){
    #check if true locus is above threshold
    results[i] <- pull.power(sim.scans = simulated.scan, thresh = simulated.threshold, window.mb = 0)
    results.window[i] <- pull.power(sim.scans = simulated.scan, thresh = simulated.threshold, window.mb = 5)
    results.false[i] <- pull.false.positive.prob(sim.scans = simulated.scan, thresh = simulated.threshold)
    gev.flag[i] <- 0
    peak.dist[i] <- pull.dist.from.locus(sim.scans = simulated.scan, thresh = simulated.threshold)
  } else {
    results[i] <- 0
    results.window[i] <- 0
    results.false[i] <- 0
    gev.flag[i] <- 1
    peak.dist[i] <- NA
  }
}

results <- list("job.id"=job.id, "genomecache"=genomecache, "n.strains"=n.strains, "n.replicates"=n.replicates, "n.sims"=n.sims, "n.alleles"=n.alleles, "h.qtl"=h.qtl, h.strain=h.strain, "seed"=seed, "results"=results, "results.window"=results.window, "results.false"=results.false, "gev.flag"=gev.flag, "peak.dist"=peak.dist)
save(results, file=paste("results/", job.id, ".RData", sep=""))
```
