## Supplementary material for "Determinants of QTL mapping power in the realized Collaborative Cross": R script to generate figures in manuscript and Supplement: generate_figures.R

```
##############################################################################
##############################################################################
########
########  Title: File S8: generate_figures.R
########  Purpose: Generate figures for SPARCC manuscript
########  Authors: Greg Keele and Wes Crouse
########
##############################################################################
##############################################################################

##############################################################################
########  
########  Necessary Packages
########
##############################################################################
library(sparcc) 
## - Purpose: Perform power simulations and analysis for Collaborative Cross (CC)
## - Available at https://github.com/gkeele/sparcc
## - If you use the R package devtools, simply use:
##   - install_github("gkeele/sparcc")

library(akima)
## - Purpose: Interpolation across a grid
## - Available on CRAN

library(pBrackets)
## - Purpose: Add brackets to allelic series grid
## - Available on CRAN

library(randomcoloR)
## - Purpose: Generate random and distinct colors
## - Available on CRAN

library(ggplot2)
## - Purpose: Used for the violin plots
## - Available on CRAN

library(superheat)
## - Purpose: Generate table/heatmap combo for Table 2
## - Available on CRAN

library(tidyverse)
## - Purpose: Convenient re-shaping of data tables
## - Available on CRAN
##############################################################################
########  
########  CC genomes
########
##############################################################################
## - Available at https://github.com/gkeele/sparcc_cache
## - Consists of a directory tree that contains founder haplotype probabilities 
##   for the 72 CC strains (available at http://csbio.unc.edu/CCstatus/index.py?run=FounderProbs})
## - Using Git with GitHub:
##   - git clone https://github.com/gkeele/sparcc_cache
##   into your personal directory, here abbreviated to the home directory: ~/


##############################################################################
########  
########  Figure 1: Example allelic series grid
########
##############################################################################
set.seed(1000)
## May not produce exactly the same results across different computer architectures
full.palette <- randomcoloR::distinctColorPalette(k=8, runTsne=TRUE) 

## Generate allelic series matrix
all.series <- matrix(c(c(0,1,2,3,4,5,6,7), #8
                       c(0,0,1,2,3,4,5,6),
                       c(0,1,1,2,3,4,5,6), #7
                       c(0,0,0,1,2,3,4,5),
                       c(0,0,1,1,2,3,4,5), #6
                       c(0,0,0,0,1,2,3,4),
                       c(0,0,1,1,2,2,3,4), #5
                       c(0,0,0,0,0,1,2,3),
                       c(0,0,1,1,2,2,3,3), #4
                       c(0,0,0,0,0,0,1,2),
                       c(0,0,0,1,1,1,2,2), #3
                       c(0,0,0,0,0,0,0,1),
                       c(0,0,0,0,1,1,1,1)),
                     ncol=8, byrow=TRUE)
all.series <- all.series[rev(1:nrow(all.series)),]
colnames(all.series) <- LETTERS[1:8]

## Generate Figure 1 from manuscript
image(x=1:ncol(all.series), y=1:nrow(all.series), z=t(all.series), 
      col=full.palette, 
      xaxt="n", yaxt="n", xlab="", ylab="")
grid(nx=8, ny=nrow(all.series), col="gray30", lty=1)
axis(1, at=1:ncol(all.series), labels=colnames(all.series), srt=45, tick=FALSE, cex.axis=1)
brackets(x1=0.4, x2=0.4, y1=2.5, y2=4.5, lwd=3, xpd=TRUE, h=0.25, type=4)
mtext(2, at=3.5, adj=1.4, text="3 alleles", las=1, xpd=TRUE, cex=0.9)
brackets(x1=0.4, x2=0.4, y1=6.5, y2=8.5, lwd=3, xpd=TRUE, h=0.25, type=4)
mtext(2, at=7.5, adj=1.4, text="5 alleles", las=1, xpd=TRUE, cex=0.9)
brackets(x1=0.4, x2=0.4, y1=10.5, y2=12.5, lwd=3, xpd=TRUE, h=0.25, type=4)
mtext(2, at=11.5, adj=1.4, text="7 alleles", las=1, xpd=TRUE, cex=0.9)
lines(x=c(0, 10), y=c(2.5, 2.5), lwd=3, col="gray30")
lines(x=c(0, 10), y=c(4.5, 4.5), lwd=3, col="gray30")
lines(x=c(0, 10), y=c(6.5, 6.5), lwd=3, col="gray30")
lines(x=c(0, 10), y=c(8.5, 8.5), lwd=3, col="gray30")
lines(x=c(0, 10), y=c(10.5, 10.5), lwd=3, col="gray30")
lines(x=c(0, 10), y=c(12.5, 12.5), lwd=3, col="gray30")
## Legend
plot.new()
legend("right", legend=paste("Allele", c(1,5,2,6,3,7,4,8)), 
       fill=c(full.palette[1], full.palette[5], full.palette[2], full.palette[6],
              full.palette[3], full.palette[7], full.palette[4], full.palette[8]), 
       border="gray30", bty="n", ncol=4)


##############################################################################
########  
########  Figure 2: Small scale demonstration for paper
########
##############################################################################
## Useful functions for parsing haplotype data
h <- DiploprobReader$new("./sparcc_cache/")
## May not produce exactly the same results across different computer architectures
set.seed(100)
these.cc.lines <- sample(h$getSubjects(), size=65)

## Simulation of five realizations
simple.data <- sim.CC.data(genomecache="~/sparcc_cache/", 
                           CC.lines=these.cc.lines,
                           num.replicates=5,
                           num.sim=5,
                           locus=NULL,
                           num.alleles=2,
                           qtl.effect.size=0.1, 
                           strain.effect.size=0)

## Genome scan
simple.scans <- run.sim.scans(sim.data=simple.data,
                              all.sim.qr=NULL,
                              return.all.sim.qr=TRUE, print.scans.progress=TRUE)

## Generation permutations and corresponding scans
permutation.index <- generate.perm.matrix(num.lines=65, num.perm=100)
thresh.scans <- run.perm.scans(perm.matrix=permutation.index, 
                               sim.CC.object=simple.data, 
                               sim.CC.scans=simple.scans,
                               print.scans.progress=FALSE)

## Pulling significance thresholds
all.thresh <- get.thresholds(thresh.scans=thresh.scans)

## Calculating power from scans and thresholds
pull.power(sim.scans=simple.scans, thresh=all.thresh)

## Calculating distance from detected hits and simulated QTL
pull.dist.from.locus(sim.scans=simple.scans, thres=all.thresh)

## Generate Figure 2 from manuscript
single.sim.plot(simple.scans, thresh=all.thresh, phenotype.index=1, y.max.manual=7,
                mark.locus.col="blue")
single.sim.plot(simple.scans, thresh=all.thresh, phenotype.index=2, y.max.manual=7,
                mark.locus.col="blue")
single.sim.plot(simple.scans, thresh=all.thresh, phenotype.index=3, y.max.manual=7,
                mark.locus.col="blue")
single.sim.plot(simple.scans, thresh=all.thresh, phenotype.index=4, y.max.manual=7,
                mark.locus.col="red")
single.sim.plot(simple.scans, thresh=all.thresh, phenotype.index=5, y.max.manual=7,
                mark.locus.col="blue")

##############################################################################
########  
########  Figure 3: Power panel
########
##############################################################################
## May not produce exactly the same results across different computer architectures
set.seed(100)
these.col <- distinctColorPalette(k=7, runTsne=TRUE)
these.effect.size <- c(0.01, 0.05, round(seq(0.1, 0.5, by=0.1), 1))

## Generate Figure 3 subplot with 1 observation and 2 alleles
power.plot(results=r1.dat, 
           qtl.effect.size=these.effect.size[1], 
           n.alleles=2, 
           n.replicates=1, 
           col=these.col[1], 
           alpha=0.5)
for (i in 2:7) {
  add.curve.to.power.plot(results=r1.dat, 
                          qtl.effect.size=these.effect.size[i], 
                          n.alleles=2, 
                          n.replicates=1, 
                          col=these.col[i], 
                          pch=20, 
                          alpha=0.5)
}
abline(h=0.8, col="red", lty=3)
abline(v=58, lty=2, col="black")
legend("left", legend=c("50%", "40%", "30%", "20%", "10%", "5%", "1%"), 
       fill=rev(these.col), border=rep(NA, 7), bty="n", ncol=2)

## Generate Figure 3 subplot with 1 observation and 3 alleles
power.plot(results=r1.dat, 
           qtl.effect.size=these.effect.size[1], 
           n.alleles=3, 
           n.replicates=1, 
           col=these.col[1], 
           alpha=0.5)
for (i in 2:7) {
  add.curve.to.power.plot(results=r1.dat, 
                          qtl.effect.size=these.effect.size[i], 
                          n.alleles=3, 
                          n.replicates=1, 
                          col=these.col[i], 
                          pch=20, 
                          alpha=0.5)
}
abline(h=0.8, col="red", lty=3)
abline(v=58, lty=2, col="black")

## Generate Figure 3 subplot with 1 observation and 8 alleles
power.plot(results=r1.dat, 
           qtl.effect.size=these.effect.size[1], 
           n.alleles=8, 
           n.replicates=1, 
           col=these.col[1], 
           alpha=0.5)
for (i in 2:7) {
  add.curve.to.power.plot(results=r1.dat, 
                          qtl.effect.size=these.effect.size[i], 
                          n.alleles=8, 
                          n.replicates=1, 
                          col=these.col[i], 
                          pch=20, 
                          alpha=0.5)
}
abline(h=0.8, col="red", lty=3)
abline(v=58, lty=2, col="black")

## Generate Figure 3 subplot with 5 replicates and 2 alleles
r5.a2.dat <- interpolate.table(r1.results=r1.dat, 
                               num.replicates=5, 
                               n.alleles=2)
power.plot(results=r5.a2.dat, 
           qtl.effect.size=these.effect.size[1], 
           n.alleles=2, 
           n.replicates=5, 
           col=these.col[1], 
           alpha=0.5, 
           pch=1)
for (i in 2:7) {
  add.curve.to.power.plot(results=r5.a2.dat, 
                          qtl.effect.size=these.effect.size[i], 
                          n.alleles=2, n.replicates=5, 
                          col=these.col[i], 
                          alpha=0.5, 
                          pch=1)
}
abline(h=0.8, col="red", lty=3)
abline(v=58, lty=2, col="black")

## Generate Figure 3 subplot with 5 replicates and 3 alleles
r5.a3.dat <- interpolate.table(r1.results=r1.dat, 
                               num.replicates=5, 
                               n.alleles=3)
power.plot(results=r5.a3.dat, 
           qtl.effect.size=these.effect.size[1], 
           n.alleles=3, 
           n.replicates=5, 
           col=these.col[1], 
           alpha=0.5, 
           pch=1)
for (i in 2:7) {
  add.curve.to.power.plot(results=r5.a3.dat, 
                          qtl.effect.size=these.effect.size[i], 
                          n.alleles=3, 
                          n.replicates=5,
                          col=these.col[i], 
                          alpha=0.5, 
                          pch=1)
}
abline(h=0.8, col="red", lty=3)
abline(v=58, lty=2, col="black")

## Generate Figure 3 subplot with 5 replicates and 8 alleles
r5.a8.dat <- interpolate.table(r1.results=r1.dat, 
                               num.replicates=5, 
                               n.alleles=8)
power.plot(results=r5.a8.dat, 
           qtl.effect.size=these.effect.size[1], 
           n.alleles=8, 
           n.replicates=5, 
           col=these.col[1], 
           alpha=0.5, 
           pch=1)
for (i in 2:7) {
  add.curve.to.power.plot(results=r5.a8.dat, 
                          qtl.effect.size=these.effect.size[i], 
                          n.alleles=8, 
                          n.replicates=5, 
                          col=these.col[i], 
                          alpha=0.5)
}
abline(h=0.8, col="red", lty=3)
abline(v=58, lty=2, col="black")

## Generate Figure 3 subplot with 5 replicates, 2 alleles, and background strain effect
r5.a2.h50.dat <- interpolate.table(r1.results=r1.dat, 
                                   num.replicates=5, 
                                   strain.effect.size=0.5-these.effect.size[1], 
                                   n.alleles=2)
power.plot(results=r5.a2.h50.dat, 
           qtl.effect.size=these.effect.size[1], 
           n.alleles=2, 
           n.replicates=5, 
           col=these.col[1], 
           alpha=0.5, 
           pch=1)
for (i in 2:7) {
  this.dat <- interpolate.table(r1.results=r1.dat, 
                                num.replicates=5, 
                                strain.effect.size=0.5-these.effect.size[i],
                                n.alleles=2)
  add.curve.to.power.plot(results=this.dat, 
                          qtl.effect.size=these.effect.size[i], 
                          n.alleles=2, 
                          n.replicates=5, 
                          col=these.col[i], 
                          alpha=0.5, 
                          pch=1)
}
abline(h=0.8, col="red", lty=3)
abline(v=58, lty=2, col="black")

## Generate Figure 3 subplot with 5 replicates, 3 alleles, and background strain effect
r5.a3.h50.dat <- interpolate.table(r1.results=r1.dat, 
                                   num.replicates=5, 
                                   strain.effect.size=0.5-these.effect.size[1], 
                                   n.alleles=3)
power.plot(results=r5.a3.h50.dat, 
           qtl.effect.size=these.effect.size[1], 
           n.alleles=3, 
           n.replicates=5, 
           col=these.col[1], 
           alpha=0.5, 
           pch=1)
for (i in 2:7) {
  this.dat <- interpolate.table(r1.results=r1.dat, 
                                num.replicates=5, 
                                strain.effect.size=0.5-these.effect.size[i],
                                n.alleles=3)
  add.curve.to.power.plot(results=this.dat, 
                          qtl.effect.size=these.effect.size[i], 
                          n.alleles=3, 
                          n.replicates=5, 
                          col=these.col[i], 
                          alpha=0.5, 
                          pch=1)
}
abline(h=0.8, col="red", lty=3)
abline(v=58, lty=2, col="black")

## Generate Figure 3 subplot with 5 replicates, 8 alleles, and background strain effect
r5.a8.h50.dat <- interpolate.table(r1.results=r1.dat, 
                                   num.replicates=5, 
                                   strain.effect.size=0.5-these.effect.size[1], 
                                   n.alleles=8)
power.plot(results=r5.a8.h50.dat, 
           qtl.effect.size=these.effect.size[1], 
           n.alleles=8, 
           n.replicates=5, 
           col=these.col[1], 
           alpha=0.5, 
           pch=1)
for (i in 2:7) {
  this.dat <- interpolate.table(r1.results=r1.dat, 
                                num.replicates=5, 
                                strain.effect.size=0.5-these.effect.size[i],
                                n.alleles=8)
  add.curve.to.power.plot(results=this.dat, 
                          qtl.effect.size=these.effect.size[i], 
                          n.alleles=8, 
                          n.replicates=5, 
                          col=these.col[i], 
                          alpha=0.5, 
                          pch=1)
}
abline(h=0.8, col="red", lty=3)
abline(v=58, lty=2, col="black")

##############################################################################
########  
########  Figures 4 and S3: Location error table
########
##############################################################################
########## QTL distance table with and without regularization
n.mean.pseudo <- 10
n.ci.pseudo <- 10
n.alleles <- 8

n.strains <- sort(unique(r1.dist.dat$n.strains))
h.qtl <- sort(unique(r1.dist.dat$h.qtl))

results.mean <- matrix(NA, length(n.strains), length(h.qtl))
colnames(results.mean) <- h.qtl
rownames(results.mean) <- n.strains

results.ci <- results.mean
results.mean.pseudo <- results.mean
results.ci.pseudo <- results.mean

n.per.cell <- matrix(NA, length(n.strains), length(h.qtl))

for (i in n.strains){
  for (j in h.qtl){
    current.results <- r1.dist.dat[r1.dist.dat$n.strains==i & r1.dist.dat$h.qtl==j & r1.dist.dat$n.alleles==n.alleles,]
    
    n <- nrow(current.results)
    
    n.per.cell[which(i==n.strains), which(j==h.qtl)] <- n
    
    abs.dist <- abs(current.results$dist)
    
    if (nrow(current.results)>=1){
      results.mean[which(i==n.strains), which(j==h.qtl)] <- mean(abs.dist)
      results.ci[which(i==n.strains), which(j==h.qtl)] <- quantile(abs.dist, 0.95)
    }
    
    results.mean.pseudo[which(i==n.strains), which(j==h.qtl)] <- mean(c(abs.dist, rep(2.5, n.mean.pseudo)))
    results.ci.pseudo[which(i==n.strains), which(j==h.qtl)] <- mean(c(rep(4.75, n.ci.pseudo), rep(quantile(abs.dist, 0.95), n)))
  }
}

## Figure 4A
X_text <- formatC(results.mean.pseudo, format = "f", digits = 1)
width_text <- gsub(x = formatC(se, format = "f", digits = 2), pattern = " ", replacement = "")
colnames(X_text) <- colnames(results.mean.pseudo)
rownames(X_text) <- rownames(results.mean.pseudo)
superheat(results.mean.pseudo, 
          heat.pal = colorRampPalette(c("white", "salmon"))(30),
          X.text = X_text, 
          X.text.col = "black",
          X.text.size = 5,
          heat.na.col = "black",
          order.cols = 1:ncol(results.mean.pseudo),
          order.rows = rev(1:nrow(results.mean.pseudo)),
          column.title = "QTL effect size",
          row.title = "Number of strains",
          title = "Mean location error of detected QTL",
          left.label.size = 0.05,
          bottom.label.size = 0.05,
          grid.vline = FALSE,
          grid.hline = FALSE,
          grid.vline.col = "gray",
          grid.hline.col = "gray")

## Figure 4B
X_text <- formatC(results.ci.pseudo, format = "f", digits = 1)
width_text <- gsub(x = formatC(se, format = "f", digits = 2), pattern = " ", replacement = "")
colnames(X_text) <- colnames(results.ci.pseudo)
rownames(X_text) <- rownames(results.ci.pseudo)
superheat(results.ci.pseudo, 
          heat.pal = colorRampPalette(c("white", "skyblue"))(30),
          X.text = X_text, 
          X.text.col = "black",
          X.text.size = 5,
          heat.na.col = "black",
          order.cols = 1:ncol(results.ci.pseudo),
          order.rows = rev(1:nrow(results.ci.pseudo)),
          column.title = "QTL effect size",
          row.title = "Number of strains",
          title = "95% quantile location error of detected QTL",
          left.label.size = 0.05,
          bottom.label.size = 0.05,
          grid.vline = FALSE,
          grid.hline = FALSE,
          grid.vline.col = "gray",
          grid.hline.col = "gray")

## Figure S3A
X_text <- formatC(results.mean, format = "f", digits = 1)
width_text <- gsub(x = formatC(se, format = "f", digits = 2), pattern = " ", replacement = "")
colnames(X_text) <- colnames(results.mean)
rownames(X_text) <- rownames(results.mean)
superheat(results.mean, 
          heat.pal = colorRampPalette(c("white", "salmon"))(30),
          X.text = X_text, 
          X.text.col = "black",
          X.text.size = 5,
          heat.na.col = "black",
          order.cols = 1:ncol(results.mean),
          order.rows = rev(1:nrow(results.mean)),
          column.title = "QTL effect size",
          row.title = "Number of strains",
          title = "Row mean location error of detected QTL",
          left.label.size = 0.05,
          bottom.label.size = 0.05,
          grid.vline = FALSE,
          grid.hline = FALSE,
          grid.vline.col = "gray",
          grid.hline.col = "gray")

## Figure S3B
X_text <- formatC(results.ci, format = "f", digits = 1)
width_text <- gsub(x = formatC(se, format = "f", digits = 2), pattern = " ", replacement = "")
colnames(X_text) <- colnames(results.ci)
rownames(X_text) <- rownames(results.ci)
superheat(results.mean, 
          heat.pal = colorRampPalette(c("white", "skyblue"))(30),
          X.text = X_text, 
          X.text.col = "black",
          X.text.size = 5,
          heat.na.col = "black",
          order.cols = 1:ncol(results.ci),
          order.rows = rev(1:nrow(results.ci)),
          column.title = "QTL effect size",
          row.title = "Number of strains",
          title = "Raw 95% quantile location error of detected QTL",
          left.label.size = 0.05,
          bottom.label.size = 0.05,
          grid.vline = FALSE,
          grid.hline = FALSE,
          grid.vline.col = "gray",
          grid.hline.col = "gray")

##############################################################################
########  
########  Figure 5: Generate power heatmap for number of 
########            CC strains vs number of replicates
########
##############################################################################
## Colors
color.palette <- colorRampPalette(c("darkslategray3", "firebrick3"))

## Function for interpolating the grid
interpolate.qtl.power.grid <- function(num.replicates = c(1:10),
                                       n.strains = c(seq(10, 70, by = 5), 72),
                                       qtl.effect.size=c(0.01, seq(0.05, 0.95, by = 0.05)),
                                       this.qtl.effect.size = 0.2,
                                       strain.effect.size = 0,
                                       n.alleles,
                                       use.window = TRUE,
                                       r1.results) {
  
  final.data <- NULL
  for(i in 1:length(n.strains)) {
    temp <- sapply(1:length(num.replicates), function(x) interpolate.qtl.power(qtl.effect.sizes = qtl.effect.size,
                                                                               strain.effect.size = strain.effect.size,
                                                                               num.replicates = num.replicates[x],
                                                                               n.alleles = n.alleles,
                                                                               use.window = TRUE,
                                                                               n.strains = n.strains[i], 
                                                                               r1.results = r1.results))
    rownames(temp) <- qtl.effect.size
    colnames(temp) <- num.replicates
    temp.data <- reshape2::melt(temp)
    temp.data <- cbind(rep(n.strains[i], nrow(temp.data)), temp.data)
    names(temp.data) <- c("n.strains", "h.qtl", "n.replicates", "power")
    final.data <- rbind(final.data, temp.data)
  } 
  
  ## Processing evaluated power
  power <- final.data[final.data$h.qtl == this.qtl.effect.size,]
  if (use.window) { z <- power$power }
  else { z <- power$power.window }
  x <- power$n.strains
  y <- power$n.replicates
  a <- x*y
  
  power.grid <- akima::interp(a, y, z,
                              xo=seq(1, 720, by=0.1),
                              yo=seq(1, 10, by=0.1),
                              linear = TRUE) 
  
  return(power.grid)
}

## Generate Figure 5 for manuscript
power.grid.plot <- function(power.grid,
                            col=colorRampPalette(c("darkslategray3", "firebrick3"))(30),
                            line.col = "gray40",
                            xlab = "Number of mice",
                            ylab = "Number of replicates",
                            zlab = "Number of strains") {
  image(power.grid$x, power.grid$y, power.grid$z, 
        xlim = c(1,720),
        ylim = c(1,10), 
        xlab = xlab, 
        ylab = ylab, 
        col = col, 
        las = 1,
        yaxt = "n")
  axis(side=2, at=1:10, las=1)
  axis(side=3, at=seq(100, 700, by=100), labels=seq(10, 70, by=10))
  mtext(side=3, zlab, line=3, cex=1)
  
  reps <- 1:10
  
  these.lwd <- ifelse(seq(10, 70, by=5) %% 10 == 0, 1, 0.3)
  strains <- seq(10, 70, by=5)
  
  for (i in 1:length(strains)) {
    these.x <- reps*strains[i]
    if (any(these.x > 240)) {
      low.x <- these.x[these.x < 240]
      low.x <- c(low.x, 240)
      low.y <- reps[these.x < 240]
      low.y <- c(low.y, (1/strains[i])*240)
      lines(low.x, low.y, lwd=0.5, lty=1, col=line.col)
      
      high.x <- these.x[these.x > 260]
      high.x <- c(260, high.x)
      high.y <- reps[these.x > 260]
      high.y <- c((1/strains[i])*260, high.y)
      lines(high.x, high.y, lwd=0.5, lty=1, col=line.col)
    }
    else {
      lines(reps*strains[i], reps, lwd=0.5, lty=1, col=line.col)
    }
    lines(reps*strains[i])
  }
  
  lines(x=c(240, 240), y=c((1/72)*240, 10), lwd=2, col=line.col, lty=1)
  lines(x=c(260, 260), y=c((1/72)*260, 10), lwd=2, col=line.col, lty=1)
  abline(coef=c(0,1/72), lwd=6, lty=1)
  abline(coef=c(0,1/10), lwd=6, lty=1)
}

## Power grid
grid.power.02 <- interpolate.qtl.power.grid(strain.effect.size=0,
                                            n.alleles=2,
                                            r1.results=r1.dat)
power.grid.plot(grid.power.02)

## Color ramp for Figure 5
plot.new()
color.palette <- colorRampPalette(c("darkslategray3", "firebrick3"))
these.colors <- color.palette(30)
barplot(height=rep(0.1, length(these.colors)), width=1/length(these.colors), density=1000,
        angle=90, col=these.colors, 
        border=FALSE, space=FALSE, axes=FALSE, add=TRUE)
axis(1, at=0:1, labels=0:1, tick=TRUE, cex.axis=1.2)
mtext(side=1, "Power", line=2)


##############################################################################
########  
########  Figure 6: Generate FPR by strain effect (with K) plot
########
##############################################################################
exchange_results_72 <- r1.exchange.dat %>% filter(n.strains == 72)
exchange_results_71 <- r1.exchange.dat %>% filter(n.strains == 71)

plot(y = exchange_results_72$results.false, 
     x = exchange_results_72$h.strain * 100, 
     type = "b", 
     ylim = c(0, 0.2), 
     las = 1, 
     frame.plot = FALSE,
     ylab = "False positive rate",
     xlab = "Strain effect size (%)",
     lwd = 4, 
     pch = 20)
points(y = exchange_results_71$results.false,
       x = exchange_results_71$h.strain * 100,
       lwd = 4,
       type = "b",
       pch = 20,
       col = "gray")
abline(h = 0.05, col = "red", lty = 2, lwd = 3)
legend("topright", legend = c("72 strains", "71 strains (One cousin)"),
       bty = "n", lty = c(3, 3), lwd = c(3, 3), pch = c(20, 20), col = c("black", "gray"))

##############################################################################
########  
########  Figure 7: Beavis effect in the CC
########
##############################################################################
## Converting number of strains and effect size to factors
r1.beavis.dat$n.strains <- as.factor(r1.beavis.dat$n.strains)
r1.beavis.dat$h.qtl <- as.factor(r1.beavis.dat$h.qtl * 100)
## Ratio of estimated effect size to simulated effect size
r1.beavis.dat$effect_size_ratio <- (r1.beavis.dat$effect.size * 100)/as.numeric(as.character(r1.beavis.dat$h.qtl))
beavis_plot <- ggplot(data = r1.beavis.dat %>% filter(power.window == 1), 
                       aes(x = h.qtl, 
                           y = effect_size_ratio, 
                           fill = n.strains, 
                           color = n.strains))
beavis_plot <- beavis_plot + scale_fill_discrete(guide = FALSE)
beavis_plot <- beavis_plot + scale_color_manual(name = "Number of strains", values = colorRampPalette(colors = c("tomato", "cadetblue"))(4))
beavis_plot <- beavis_plot + geom_point(color = "black", position = position_jitterdodge(), size = 0.95, alpha = 0.4)
beavis_plot <- beavis_plot + geom_boxplot(fill = "white", alpha=1, size = 1.1, position = pd, outlier.shape = NA)
beavis_plot <- beavis_plot + stat_summary(fun.y = mean, 
                                            color = "black", 
                                            geom = "point",
                                            position = pd,
                                            shape = 18, size = 3)
beavis_plot <- beavis_plot + geom_hline(yintercept = 1, color = "red", linetype = "longdash", size = 1.25)
beavis_plot <- beavis_plot + ylab("Estimated QTL effect size / Simulated QTL effect size") + xlab("Simulated QTL effect size (%)")
beavis_plot <- beavis_plot + theme(panel.grid.major = element_blank(), 
                                     panel.grid.minor = element_blank(),
                                     panel.background = element_blank(), 
                                     axis.line = element_line(colour = "black", size = 1),
                                     plot.title = element_text(hjust = 0.5), 
                                     axis.title = element_text(size = 12),
                                     axis.text = element_text(size = 12))
beavis_plot

##############################################################################
########  
########  Figure 8: QTL effect size in different reference populations
########
##############################################################################
## May not produce exactly the same results across different computer architectures
set.seed(100)
## Effect sizes in 10,000 simulations of bi-allelic QTL
two.allele.data <- sim.CC.data(genomecache="~/Documents/Wes_DOQTL/sparcc_cache/",
                               num.lines=72,
                               num.replicates=1,
                               num.sim=10000,
                               locus=NULL,
                               num.alleles=2,
                               qtl.effect.size=0.5, 
                               strain.effect.size=0,
                               scale.qtl.mode="B")
## Effect sizes in 10,000 simulations of QTL with eight alleles
eight.allele.data <- sim.CC.data(genomecache="~/Documents/Wes_DOQTL/sparcc_cache/",
                                 num.lines=72,
                                 num.replicates=1,
                                 num.sim=10000,
                                 locus=NULL,
                                 num.alleles=8,
                                 qtl.effect.size=0.5, 
                                 strain.effect.size=0,
                                 scale.qtl.mode="B")

## Grab only the effect sizes scaled from balanced population
two.data.B <- two.allele.data$properties$var.tables$B
eight.data.B <- eight.allele.data$properties$var.tables$B
## Grab only the effect sizes scaled from observed population
two.data.DAMB <- two.allele.data$properties$var.tables$DAMB
eight.data.DAMB <- eight.allele.data$properties$var.tables$DAMB
## Combine into a single data set
dat1 <- rbind(data.frame(B=two.data.B[,"DAMB.ve"], n.alleles="2", population="Mapping", scale="2 in balanced"),
              data.frame(B=eight.data.B[,"DAMB.ve"], n.alleles="8", population="Mapping", scale="8 in balanced"),
              data.frame(B=two.data.DAMB[,"B.ve"], n.alleles="2", population="Balanced", scale="2 in mapping"),
              data.frame(B=eight.data.DAMB[,"B.ve"], n.alleles="8", population="Balanced", scale="8 in mapping"))

## Generate Figure 7 A from manuscript
p <- ggplot(dat1, aes(x=n.alleles, y=B, fill=scale))#, color=n.alleles)) 
p <- p + ggtitle("QTL effect size")
p <- p + geom_jitter(shape=16, position=position_jitter(0.2), col=scales::alpha("gray", 0.6)) + ylim(0, 1)
p <- p + geom_violin(size=1.1, adjust=1, bw=0.03, draw_quantiles=c(0.25, 0.5, 0.75)) 
p <- p + scale_fill_manual(values=c("#FB9A99", "#E31A1C", "#A6CEE3", "#1F78B4"))
p <- p + scale_color_manual(values=c("black", "black", "black", "black"))
p <- p + theme_bw() + theme(panel.border = element_blank(), panel.grid.major = element_blank(),
                            panel.grid.minor = element_blank(), axis.line = element_line(colour = "black"),
                            #axis.text.x=element_blank(), axis.ticks.x=element_blank(), axis.title.x=element_blank(),
                            plot.title = element_text(hjust = 0.5), axis.text=element_text(color="black"))
p <- p + ylab("QTL effect size") + xlab("Number of alleles")
p <- p + guides(fill=guide_legend(title="Scaled in:", override.aes = list(shape = NA, col=NA)), col=FALSE)
p <- p + facet_wrap(~population)
p

## Generate Figure 7 B from manuscript
power.plot(results=r1.dat,
           qtl.effect.size=0.5, n.alleles=2, n.replicates=1, col="#FB9A99", alpha=0.5)
add.curve.to.power.plot(results=r1.dat,
                        qtl.effect.size=0.5, n.alleles=8, n.replicates=1, col="#E31A1C", pch=20, alpha=0.5)
add.curve.to.power.plot(results=r1.damb.dat,
                        qtl.effect.size=0.5, n.alleles=2, n.replicates=1, col="#A6CEE3", pch=20, alpha=0.5)
add.curve.to.power.plot(results=r1.damb.dat,
                        qtl.effect.size=0.5, n.alleles=8, n.replicates=1, col="#1F78B4", pch=20, alpha=0.5)
abline(h=0.8, col="red", lty=3)
abline(v=58, lty=2, col="black")

##############################################################################
########  
########  Figure 9: Allelic balance in power to map bi-allelic QTL
########
##############################################################################
allelic.series <- as.matrix(expand.grid(c(1,0), c(1,0), c(1,0), c(1,0), c(1,0), c(1,0), c(1,0), c(1,0)))[2:128,]
colnames(allelic.series) <- LETTERS[1:8]

for (i in 1:nrow(allelic.series)) {
  if (sum(allelic.series[i,]) > 4) {
    allelic.series[i,] <- ifelse(allelic.series[i,] == 1, 0, 1)
  }
  if (sum(allelic.series[i,]) == 4 & allelic.series[i,1] == 1) {
    allelic.series[i,] <- ifelse(allelic.series[i,] == 1, 0, 1)
  }
}

## Balance: 4v4
combo.4v4 <- allelic.series[rowSums(allelic.series) == 4,]
## Balance: 7v1
combo.7v1 <- allelic.series[rowSums(allelic.series) %in% c(1,7),]
## Balance: 6v2
combo.6v2 <- allelic.series[rowSums(allelic.series) %in% c(2,6),]
## Balance: 5v3
combo.5v3 <- allelic.series[rowSums(allelic.series) %in% c(3,5),]
## Combine all allelic series
all.combo <- rbind(combo.7v1, combo.6v2, combo.5v3, combo.4v4)

## Generate Figure 8 A from manuscript
image(x=1:ncol(allelic.series), y=1:nrow(all.combo), z=t(rbind(combo.7v1, combo.6v2, combo.5v3, combo.4v4)), 
      col=c(full.palette[1], full.palette[2]), 
      xaxt="n", yaxt="n", xlab="", ylab="")
grid(nx=8, ny=nrow(all.combo), col="gray30", lty=1)
axis(1, at=1:ncol(combo.5v3), labels=colnames(combo.5v3), srt=45, tick=FALSE, cex.axis=1.2)
brackets(x1=0.4, x2=0.4, y1=0.5, y2=8.5, lwd=3, xpd=TRUE, h=0.25, type=1)
mtext(2, at=4.5, adj=1.5, text="7v1", las=1, xpd=TRUE, cex=1.5)
brackets(x1=0.4, x2=0.4, y1=8.5, y2=36.5, lwd=3, xpd=TRUE, h=0.25, type=1)
mtext(2, at=22.5, adj=1.5, text="6v2", las=1, xpd=TRUE, cex=1.5)
brackets(x1=0.4, x2=0.4, y1=36.5, y2=92.5, lwd=3, xpd=TRUE, h=0.25, type=1)
mtext(2, at=64.5, adj=1.5, text="5v3", las=1, xpd=TRUE, cex=1.5)
brackets(x1=0.4, x2=0.4, y1=92.5, y2=127.5, lwd=3, xpd=TRUE, h=0.25, type=1)
mtext(2, at=110, adj=1.5, text="4v4", las=1, xpd=TRUE, cex=1.5)
## Legend
plot.new()
legend("right", legend=c("Allele 1", "Allele 2"), 
       fill=c(full.palette[1], full.palette[2]), border="gray30", bty="n", ncol=2)

## Generate Figure 8 B from manuscript
## Make color spectrum
magenta.cyan.intermediate <- colorRampPalette(c("magenta", "cyan"))(3)[2]
power.plot(results=r1.4v4.dat,
           qtl.effect.size=0.5,
           n.alleles=2, 
           col="magenta")
add.curve.to.power.plot(results=r1.dat,
                        qtl.effect.size=0.5,
                        n.alleles=2,
                        col=magenta.cyan.intermediate,
                        pch=20)
add.curve.to.power.plot(results=r1.7v1.dat,
                        qtl.effect.size=0.5,
                        n.alleles=2,
                        col="cyan",
                        pch=20)
legend("left", legend=c("4v4", "Uniform", "7v1"),
       col=c("magenta", magenta.cyan.intermediate, "cyan"),
       fill=c("magenta", magenta.cyan.intermediate, "cyan"),
       border=c(NA, NA, NA),
       bty="n")
abline(h=0.8, col="red", lty=3)
abline(v=58, lty=2, col="black")

##############################################################################
########  
########  Figure S1: Power interpolation of experiments with replicates
########
##############################################################################
## Interpolate power for experiment with 3 replicates
r3.interp.dat <- interpolate.table(r1.results=r1.dat,
                                   num.replicates=3,
                                   n.alleles=2,
                                   use.window=TRUE)
## Interpolate power for experiment with 5 replicates
r5.interp.dat <- interpolate.table(r1.results=r1.dat,
                                   num.replicates=5,
                                   n.alleles=2,
                                   use.window=TRUE)

## Generate Figure S1 from manuscript
power.plot(results=r1.dat,
           qtl.effect.size=0.3,
           n.alleles=2)
add.curve.to.power.plot(results=r3.interp.dat,
                        qtl.effect.size=0.3,
                        n.alleles=2,
                        col="skyblue")
add.curve.to.power.plot(results=r5.interp.dat,
                        qtl.effect.size=0.3,
                        n.alleles=2,
                        col="gray")
abline(h=0.8, col="red", lty=3)
legend("left", legend=c("5 rep", 
                        "3 rep", 
                        "1 obs"), 
       fill=c("gray", "skyblue", "firebrick3"),
       border=c(NA, NA, NA),
       bty="n")
abline(v=58, lty=2, col="black")

##############################################################################
########  
########  Figure S2: Generate FPR pandel
########
##############################################################################
## May not produce exactly the same results across different computer architectures
set.seed(100)
these.col <- distinctColorPalette(k=7, runTsne=TRUE)
these.effect.size <- c(0.01, 0.05, round(seq(0.1, 0.5, by=0.1), 1))

## Generate Figure S2 subplot with 1 observation and 2 alleles
fpr.plot(results=r1.dat, 
         qtl.effect.size=these.effect.size[1], 
         n.alleles=2, 
         n.replicates=1, 
         col=these.col[1], 
         alpha=0.5)
for (i in 2:7) {
  add.curve.to.fpr.plot(results=r1.dat, 
                        qtl.effect.size=these.effect.size[i], 
                        n.alleles=2, 
                        n.replicates=1, 
                        col=these.col[i], 
                        pch=20,
                        alpha=0.5)
}
abline(h=0.05, col="red", lty=2, lwd=3)
abline(v=58, lty=2, col="black")
legend("left", legend=c("50%", "40%", "30%", "20%", "10%", "5%", "1%"), 
       fill=rev(these.col), border=rep(NA, 7), bty="n", ncol=2)

## Generate Figure S2 subplot with 1 observation and 3 alleles
fpr.plot(results=r1.dat, 
         qtl.effect.size=these.effect.size[1], 
         n.alleles=3, 
         n.replicates=1, 
         col=these.col[1], 
         alpha=0.5)
for (i in 2:7) {
  add.curve.to.fpr.plot(results=r1.dat, 
                        qtl.effect.size=these.effect.size[i], 
                        n.alleles=3, 
                        n.replicates=1, 
                        col=these.col[i], 
                        pch=20,
                        alpha=0.5)
}
abline(h=0.05, col="red", lty=2, lwd=3)
abline(v=58, lty=2, col="black")

## Generate Figure S2 subplot with 1 observation and 8 alleles
fpr.plot(results=r1.dat, 
         qtl.effect.size=these.effect.size[1], 
         n.alleles=8, 
         n.replicates=1, 
         col=these.col[1], 
         alpha=0.5)
for (i in 2:7) {
  add.curve.to.fpr.plot(results=r1.dat, 
                        qtl.effect.size=these.effect.size[i], 
                        n.alleles=8, 
                        n.replicates=1, 
                        col=these.col[i], 
                        pch=20,
                        alpha=0.5)
}
abline(h=0.05, col="red", lty=2, lwd=3)
abline(v=58, lty=2, col="black")

##############################################################################
########  
########  Figure S4: Generate mean location error plot with increasing
########             number of replicates for sample of 350 mice
########
##############################################################################
## Function for interpolating the distance grid
interpolate.distance.grid <- function(num.replicates=c(1:10),
                                      n.strains=c(seq(10, 70, by=5), 72),
                                      qtl.effect.size=c(0.01, seq(0.05, 0.95, by=0.05)),
                                      this.qtl.effect.size=0.2,
                                      strain.effect.size=0,
                                      n.alleles,
                                      use.window=TRUE,
                                      xo = seq(1, 720, by=0.1),
                                      yo = seq(1, 10, by=0.1),
                                      r1.results) {
  
  final.data <- NULL
  for(i in 1:length(n.strains)) {
    temp <- sapply(1:length(num.replicates), function(x) interpolate.qtl.distance(qtl.effect.sizes=qtl.effect.size,
                                                                                  strain.effect.size=strain.effect.size,
                                                                                  num.replicates=num.replicates[x],
                                                                                  n.alleles=n.alleles,
                                                                                  n.strains=n.strains[i], 
                                                                                  r1.results=r1.results))
    rownames(temp) <- qtl.effect.size
    colnames(temp) <- num.replicates
    temp.data <- reshape2::melt(temp)
    temp.data <- cbind(rep(n.strains[i], nrow(temp.data)), temp.data)
    names(temp.data) <- c("n.strains", "h.qtl", "n.replicates", "distance")
    final.data <- rbind(final.data, temp.data)
  } 
  
  ## Processing evaluated power
  distance <- final.data[final.data$h.qtl == this.qtl.effect.size,]
  z <- abs(distance$dist)
  x <- distance$n.strains
  y <- distance$n.replicates
  a <- x*y
  
  distance.grid <- akima::interp(a, y, z,
                                 xo = xo,
                                 yo = yo,
                                 linear = TRUE) 
  
  return(distance.grid)
}

## Distance grid
fix_250 <- interpolate.distance.grid(strain.effect.size=0,
                                     xo = 250,
                                     n.alleles=8,
                                     this.qtl.effect.size = 0.5,
                                     r1.results=r1.dist.dat)

plot(x = fix_250$y[fix_250$y >= 4], 
     y = fix_250$z[fix_250$y >= 4], 
     xlim = c(4, max(fix_250$y)),
     #ylim = c(0, max(fix_250$z, na.rm = TRUE)),
     type = "l",
     frame.plot = FALSE,
     lwd = 2,
     xlab = "Number of replicates",
     ylab = "Mean location error (Megabase)",
     las = 1,
     main = "Total sample size: 250 mice")
abline(h = min(fix_250$z[fix_250$y >= 4]), col = "red", lty = 2)

##############################################################################
########  
########  Figure S5: Generate comparison of singular values of kinship plot
########
##############################################################################
## Subfigure A
## Calculate average off-diagonal of realized K
av_offdiag <- mean(K[upper.tri(K)])
## Make perfectly balanced K to compare to realized K
balanced_K <- (diag(72)*(1 - av_offdiag)) + av_offdiag
## Taken eigen decompositions
eigen_K <- eigen(K, symmetric = TRUE)
eigen_balanced_K <- eigen(balanced_K, symmetric = TRUE)
## Plot singlular values
plot(x = 1:72, y = eigen_K$value, xlab = "Components", ylab = "Eigenvalues", 
     pch = 19, las = 1, frame.plot = FALSE, cex = 1.1, ylim = c(0, 12), col = "firebrick1",
     main = "Deviation from balance in realized K")
points(x = 1:72, y = eigen_balanced_K$value, pch = 18, cex = 1.1)
abline(v = min(which(cumsum(eigen_K$value)/sum(eigen_K$value) > 0.95)), lty = 2, col = "firebrick1")
abline(v = min(which(cumsum(eigen_balanced_K$value)/sum(eigen_balanced_K$value) > 0.95)), lty = 2)
legend("topright", legend = c("Realized K", "Balanced K"), 
       pch = c(19, 18), col = c("red", "black"), bty = "o",
       box.lwd = 2, box.lty = 1, bg = "white")

## Subfigure B
## Remove CC059 (cousin strain)
reduced_K <- K[colnames(K) != "CC059", colnames(K) != "CC059"]
red_av_offdiag <- mean(reduced_K[upper.tri(reduced_K)])
## Make perfectly balanced K to compare to realized K
reduced_balanced_K <- (diag(71)*(1 - red_av_offdiag)) + red_av_offdiag
## Taken eigen decompositions
eigen_reduced_K <- eigen(reduced_K, symmetric = TRUE)
eigen_reduced_balanced_K <- eigen(reduced_balanced_K, symmetric = TRUE)
## Plot singlular values
plot(x = 1:71, y = eigen_reduced_K$value, xlab = "Components", ylab = "Eigenvalues", 
     pch = 19, las = 1, frame.plot = FALSE, cex = 1.1, ylim = c(0, 12), col = "darkturquoise",
     main = "Deviation from balance in realized K (CC059 omitted)")
points(x = 1:71, y = eigen_reduced_balanced_K$value, pch = 18, cex = 1.1)
abline(v = min(which(cumsum(eigen_reduced_K$value)/sum(eigen_reduced_K$value) > 0.95)), lty = 2, col = "darkturquoise")
abline(v = min(which(cumsum(eigen_reduced_balanced_K$value)/sum(eigen_reduced_balanced_K$value) > 0.95)), lty = 2)
legend("topright", legend = c("Realized K", "Balanced K"), 
       pch = c(19, 18), col = c("darkturquoise", "black"), bty = "o",
       box.lwd = 2, box.lty = 1, bg = "white")


##############################################################################
########  
########  Figure S6: Generate distance between detected QTL signal 
########             and simulated QTL
########
##############################################################################
set.seed(100)
## May not produce exactly the same results across different computer architectures
## Generate Figure S6 A for distance densities by number of strains
color.palette <- colorRampPalette(c("seagreen3", "royalblue3"))
colScale <- scale_colour_manual(name="h.qtl", 
                                values=color.palette(length(unique(r1.dist.dat$n.strains))))
fillScale <- scale_fill_manual(name="h.qtl", 
                               values=color.palette(length(unique(r1.dist.dat$n.strains))))
r1.dist.dat$n.strains <- as.factor(r1.dist.dat$n.strains)
p <- ggplot(r1.dist.dat %>% filter(h.qtl == 0.5, n.alleles == 8), 
            aes(x=n.strains, y=dist, fill=n.strains)) 
p <- p + geom_jitter(shape=16, position=position_jitter(0.2), col=scales::alpha("gray", 0.6))
p <- p + geom_violin(size=0.5, adjust=1.5, draw_quantiles=c(0.25, 0.75)) 
p <- p + stat_summary(fun.y=mean, geom="point", shape=20, size=1.5)
p <- p + theme_bw() + theme(panel.border = element_blank(), panel.grid.major = element_blank(),
                            panel.grid.minor = element_blank(), axis.line = element_line(colour = "black"),
                            axis.text=element_text(color="black"))
p <- p + fillScale + xlab("Number of strains") + ylab("Distance from QTL position (Megabase)")
p <- p + guides(fill=guide_legend(title="Number of strains", override.aes = list(shape = NA)))
p

## Generate Figure S6 B for distance densities by number of alleles
color.palette <- randomcoloR::distinctColorPalette(k=3, runTsne=TRUE)
colScale <- scale_colour_manual(name="n.alleles", 
                                values=color.palette)
fillScale <- scale_fill_manual(name="n.alleles", 
                               values=color.palette)
r1.dist.dat$n.alleles <- as.factor(r1.dist.dat$n.alleles)
p <- ggplot(r1.dist.dat %>% filter(h.qtl == 0.5, n.strains == 72), 
            aes(x=n.alleles, y=dist, fill=n.alleles)) 
p <- p + geom_jitter(shape=16, position=position_jitter(0.2), col=scales::alpha("gray", 0.6))
p <- p + geom_violin(size=0.5, adjust=1.5, draw_quantiles=c(0.25, 0.75)) 
p <- p + stat_summary(fun.y=mean, geom="point", shape=20, size=1.5)
p <- p + theme_bw() + theme(panel.border = element_blank(), panel.grid.major = element_blank(),
                            panel.grid.minor = element_blank(), axis.line = element_line(colour = "black"),
                            axis.text=element_text(color="black"))
p <- p + fillScale + xlab("Number of alleles") + ylab("Distance from QTL position (Megabase)")
p <- p + guides(fill=guide_legend(title="Number of alleles", override.aes = list(shape = NA)))
p

## Generate Figure S6 C for distance densities by QTL effect size
color.palette <- colorRampPalette(c("darkgoldenrod1", "cyan4"))
colScale <- scale_colour_manual(name="h.qtl", 
                                values=color.palette(length(unique(r1.dist.dat$h.qtl))))
fillScale <- scale_fill_manual(name="h.qtl", 
                               values=color.palette(length(unique(r1.dist.dat$h.qtl))))
r1.dist.dat$h.qtl <- as.factor(r1.dist.dat$h.qtl)
p <- ggplot(r1.dist.dat %>% filter(n.strains == 72, n.alleles == 8), 
            aes(x=h.qtl, y=dist, fill=h.qtl)) 
p <- p + geom_jitter(shape=16, position=position_jitter(0.2), col=scales::alpha("gray", 0.6))
p <- p + geom_violin(size=0.5, adjust=1.5, draw_quantiles=c(0.25, 0.75)) 
p <- p + stat_summary(fun.y=mean, geom="point", shape=20, size=1.5)
p <- p + theme_bw() + theme(panel.border = element_blank(), panel.grid.major = element_blank(),
                            panel.grid.minor = element_blank(), axis.line = element_line(colour = "black"),
                            axis.text=element_text(color="black"))
p <- p + fillScale + xlab("QTL effect size") + ylab("Distance from QTL position (Megabase)")
p <- p + guides(fill=guide_legend(title="QTL effect size", override.aes = list(shape = NA)))
p
```
